# NMR Reveals the Synergistic Roles of Bivalent Metal Ions in Norovirus Infections

**DOI:** 10.1101/2024.07.10.602906

**Authors:** Thorben Maass, Leon Torben Westermann, Linda Sharotri, Leon Blankenhorn, Miranda Sophie Lane, Maryna Chaika, Stefan Taube, Thomas Peters, Alvaro Mallagaray

## Abstract

Norovirus is the most common cause of acute gastroenteritis worldwide. Murine noroviruses (MNV) are often used as model systems for human noroviruses (HuNoV). Therefore, it is important to identify common and divisive properties. Here, we compare the interactions of human and murine norovirus P-domains with bivalent metal ions. Binding of bivalent metal ions and bile acids to MNV P-domains have been shown to stabilize a contracted (“resting”) as opposed to an extended (“raised”) capsid conformation. This conformational change has been linked to infectivity, diarrheagenic potential, and immune escape. Likewise, the interaction of bivalent metal ions with human norovirus capsids results in contraction, suggesting a similar underlying mechanism. We used methyl TROSY NMR experiments to study the thermodynamics and kinetics of metal ion binding to P-domains, revealing a highly synergistic interaction with the bile acid glycochenodeoxycholic acid (GCDCA) for MNV. Neutralization assays support this synergistic behavior. It turns out that bivalent metal ion binding to MNV and HuNoV P-domains differs significantly. Therefore, although the transition between “raised” and “resting” capsid conformations and consequential modulation of infectivity appears to be triggered by bivalent metal ions in murine and human noroviruses, the underlying mechanisms must be different.

## Introduction

Noroviruses are non-enveloped, positive-strand RNA viruses, belonging to the family of *Caliciviridae*. Human noroviruses (HuNoVs) cause highly contagious gastroenteritis, and resulting epidemics represent a substantial burden to healthcare systems worldwide. To date, neither licensed vaccines nor antiviral drugs are available. The development of vaccines or antiviral drugs would benefit from comprehensive knowledge about the structural and dynamic features of the viral capsid proteins that directly interact with host cell receptors and with molecules of the host immune system such as monoclonal antibodies. Studies of the structure of norovirus capsids have shown that norovirus particles undergo a variety of conformational changes in the protruding (P) domain of the VP1 capsid protein as well as in the linker region between the protruding (P) and shell (S) domains^[1-15]^. The conformational transitions have been correlated with infectivity^[1,12]^ and immune escape^[6-7, 9]^.

An additional dimension of plasticity exists with respect to the formation of the VP1 dimers that make up the icosahedral viral capsid. In a previous study^[16]^ we have shown that the stability of isolated P-domain dimers (P-dimers) differs substantially between HuNoVs and murine noroviruses (MNVs). In that study we have also shown that dimerization of the P-domain of the VP1 capsid protein of MNV-1 is strongly promoted by binding of the bile acid glycochenodeoxycholic acid (GCDCA) whereas HuNoV P-domains form very stable P-dimers independent of the presence of GCDCA or other bile acids. In parallel, cryo-EM studies have shown that GCDCA also affects the overall shape of MNV virions by inducing a large-scale conformational rearrangement as reflected by a contraction of the P-domains onto the S-domains^[5]^. Follow-up studies have revealed that binding of bivalent metal ions and/or low pH induce the same overall conformational rearrangements within the capsid^[2]^. A combined cryo-EM and crystallographic study has shown that metal ions also induce a contracted conformation of GII.4 human norovirus capsids, suggesting similar underlying mechanisms for the transition from the extended to the contracted conformation^[11]^. The bivalent metal ions bind to the P-domains of the VP1 capsid protein. The binding sites identified in the crystal and cryo-EM structures are schematically visualized in Figure 1a.

**Figure 1.**
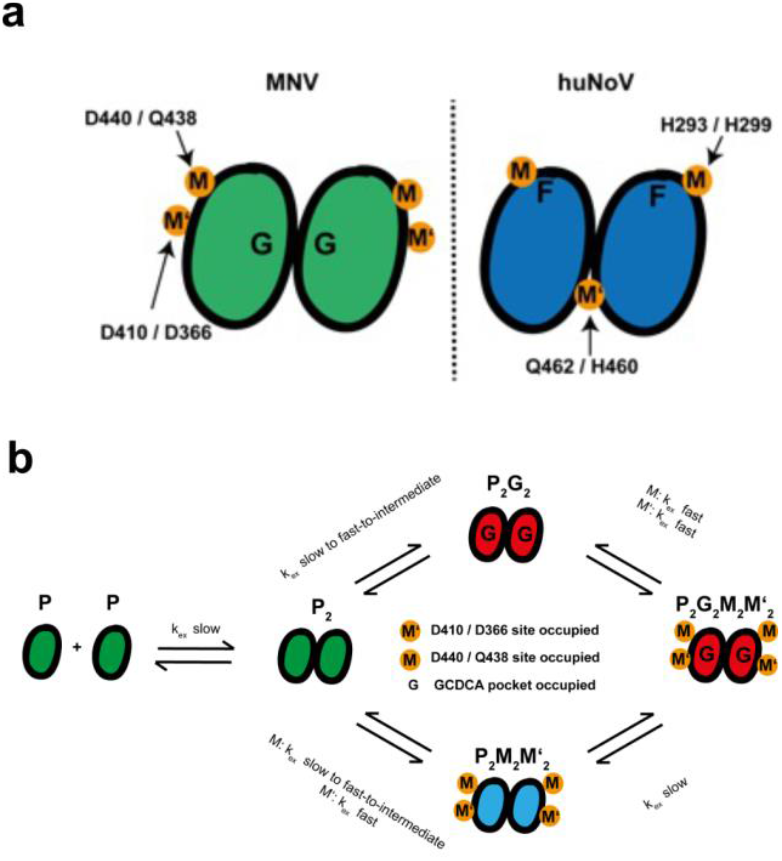
a) Cartoon representation of known binding sites for bivalent metal ions (M, M’) in MNV and HuNoV P-domain dimers (P-dimers). The green (MNV) and blue (HuNoV) ellipsoids symbolize P-domain monomers. G and F are the GCDCA and L-fucose binding sites, respectively. MNV: Mg^2+^ and Ca^2+^ bind to the D440/Q438 site (M, adjacent to the G site), and to the D410/D366 (M’, part of the receptor (CD300lf) binding site)^[17]^. HuNoV: Zn^2+^ binds to the H293/H299 site (M) of GII.2 Snow Mountain Virus (SMV) P-domains^[18]^. This site is in a loop adjacent to the F site. Cd^2+^ binds to the Q462/H460 site (M’) between the two GII.4 HOV P-domains. b) Chemical exchange between MNV P-domain states. P denotes P-domain, P2 stands for P-domain dimers (P-dimers), and M, M’ symbolize Ca^2+^ or Mg^2+^. Likewise, the extensions, e.g., “G_2_” reflect binding of the respective ligand. The different colors used for the P-domains in the different states reflect subtle overall conformational or dynamic changes as reflected by long-range CSPs associated with the formation of the different complexes. N.b.: HuNoV P-dimers studied in this work are stable and monomers are not observed under any conditions^[16]^.

An obvious question is whether the metal ion induced contraction in human and murine noroviruses follows similar or different mechanisms. Another question is whether the binding of bivalent metal ions and GCDCA is independent or not. To this end, we used methyl-TROSY NMR experiments to directly monitor the binding of bivalent metal ions and GCDCA to HuNoV and to MNV P-domains. More specifically, we used chemical shift perturbations (CSP) to obtain binding topologies and affinities. Under suitable conditions global two-dimensional line shape analysis^[19]^ provided access to both, thermodynamic and kinetic parameters of binding.

## Results

Protein-based NMR studies of ligand binding usually require an assignment of protein NMR signals as a first step. In the case of methyl TROSY experiments this means an assignment of ^13^C-methyl signals. In this study we targeted the P-domains of human and murine noroviruses. In the case of HuNoVs we used the P-domains of two strains, the GII.4 Saga and the GII.17 Kawasaki strains, as examples. For MNV we used the CW1 strain and mutants thereof. For the GII.4 Saga P-domains an assignment was available from prior work^[20]^. The GII.17 Kawasaki P-dimers were used without any further assignment as will be explained further below. For MNV P-domains assignments were also available^[16, 21]^. However, in the case of MNV P-dimers we were facing the problem that depending on the presence of co-factors the stability of P-dimers varies^[16]^ and cross peak positions shift in hard-to-predict directions. Therefore, the first step was to assign the ^1^H,^13^C-methyl groups of the MNV P-domains in the presence of bivalent metal ions.

### Assignment of ^1^H,^13^C-methyl groups of the MNV P-domain in the presence of Ca^2+^ or Mg^2+^

To study the binding of Ca^2+^ and Mg^2+^ ions to MNV P-domains by methyl TROSY NMR experiments a comprehensive assignment of ^1^H,^13^C-methyl groups of free (apo) and metal-ion bound forms of the P-domain is required. In a previous study^[21]^, we used side chain ^13^C-methyl group labeling against a highly deuterated background^[22-29]^ to obtain MNV P-domain samples specifically enriched with ^13^C-methyl labeled Ala, Ile, Leu^proS^, Met, and Val^proS^ residues, abbreviated as “MILVA”-labeled samples (for the nomenclature of specifically ^13^C-methyl labeled samples see the materials and methods section). We had been able to assign about 70% of all methyl groups for MNV P-domains in the presence of GCDCA. In a follow-up study we also obtained a partial assignment of the apo-form of the MNV P-domain^[16]^. Based on these data, we were able to assign spectral changes such as chemical shift perturbations (CSPs), or changes of signal intensities upon binding of Ca^2+^ or Mg^2+^, or due to changes in pH. From crystal structures it is known that each P-dimer has two symmetric GCDCA binding pockets (G ≔GCDCA binding pocket occupied) and two symmetric pairs of bivalent metal ion binding sites (M, M’ ≔M, M’ binding pockets occupied). Therefore, the following states are theoretically possible: apo P-domain monomers (P-monomers, P), P-monomers with GCDCA bound (PG), P-monomers with metal ion binding sites occupied (PM, PM’, or PMM’), apo P-dimers (P_2_), GCDCA bound P-dimers (P_2_G_2_), metal ion bound P-dimers (P_2_M_2_, P_2_M′_2_, P_2_M_2_M′_2_), and the P-dimer with metal ions and GCDCA bound (P_2_G_2_M_2_, PG_2_M′_2_, PG_2_M_2_M′_2_). Theoretically, single bound species like P_2_M_1_ or P_2_G_2_M_1_ may also exist. However, practical dissection of single and double-bound species in methyl TROSY spectra proved impossible. All these states are in chemical exchange, and the position of the exchange equilibria changes with the concentrations of GCDCA, metal ions, and with pH. Whether a state is spectroscopically observable depends on the respective equilibrium constants and exchange kinetics. We have noted before that monomeric P-domains complexed with GCDCA (PG) cannot be observed, and it is very likely that GCDCA does not bind to P-monomers^[16]^. Likewise, in our present study we were unable to detect P-monomers with metal ions bound (PM or PM’), and it turned out that spectroscopic discrimination of binding to the two sites is impossible. Therefore, only the five states P, P_2_, P_2_G_2_, P_2_M_2_M′_2_, and P_2_G_2_M_2_M′_2_ can be observed when using wildtype MNV P-domains. As will be shown below, inactivation of either metal ion binding site by site directed mutagenesis allowed the observation of P_2_M_2_, P_2_G_2_M_2_ and P_2_G_2_M′_2_. The chemical exchange equilibria connecting the spectroscopically observable states of wild type MNV P-domains are shown in Figure 1b.

The starting point for the assignment of methyl TROSY cross peaks to individual ^1^H,^13^C-methyl groups of the dimeric states P_2_M_2_M′_2_ and P_2_G_2_M_2_M′_2_ was based on our previous (partial) assignments for P_2_G_2_^18^ and P_2_/P^[16]^. The assignment of methyl resonance signals of P_2_G_2_M_2_M′_2_ was straightforward since the binding of Ca^2+^ or Mg^2+^ to P_2_G_2_ was fast on the chemical shift time scale, allowing continuous tracking of cross peaks during CSP titration experiments and transfer of 75 out of 77 available assignments. Only two of the 77 assignments for P_2_G_2_ could not be transferred due to spectral overlap of the methyl TROSY cross peaks of L359 and L384 (Figure S1). Assignment of methyl resonance signals for P_2_M_2_M′_2_ was not possible by tracking cross peaks during a CSP titration because binding of Ca^2+^ or Mg^2+^ to P_2_ as well as binding of GCDCA to P_2_M_2_M′_2_ was slow to intermediate on the NMR chemical shift time scale (see Figure 1). However, a comparison of the methyl TROSY spectra of the three states P_2_G_2_, P_2_G_2_M_2_M′_2_, and P_2_M_2_M′_2_ employing the nearest neighbor approach^[30-33]^ yielded an unambiguous assignment of 74 cross peaks of P_2_M_2_M′_2_. Again, a transfer of assignment of L359 and L384 was impossible due to severe spectral overlap. In addition, in the Mg^2+^-bound state (P_2_M_2_M′_2_) one of the two methyl resonances of I405 (there are two isomers of each P-domain due to a mixture of E and Z configurations of P361 ^[21]^) split further into two peaks with low intensities preventing an unambiguous assignment. This was not observed for the Ca^2+^-bound state, but with Ca^2+^ another methyl resonance, V374, was broadened beyond detection (Figure S1). Currently, we have not attempted to identify the source of these effects. Finally, using the nearest neighbor approach and comparing the assignments of the metal-bound states P_2_G_2_M_2_M′_2_ and P_2_M_2_M′_2_ to the apo state P_2_, we obtained additional assignments for the apo state.

### Binding of Ca^2+^ or Mg^2+^ to MNV P-domains causes long-range CSPs and stabilizes P-dimers

The titration of apo MNV P-domains, which consist of an equilibrium mixture of monomers (P) and dimers (P_2_), with solutions of MgCl_2_ or CaCl_2_ led to a shift of the monomer-dimer equilibrium towards the dimeric form (P_2_M_2_M′_2_). Methyl TROSY spectra reflect a depletion of the monomeric form P during titration (Figure 2a and Figure S2). The stabilization of P-dimers was also monitored using size exclusion chromatography (SEC) (Figure 2b). As explained above, at this point it was still unclear which metal binding site was responsible for the promotion of dimerization.

**Figure 2.**
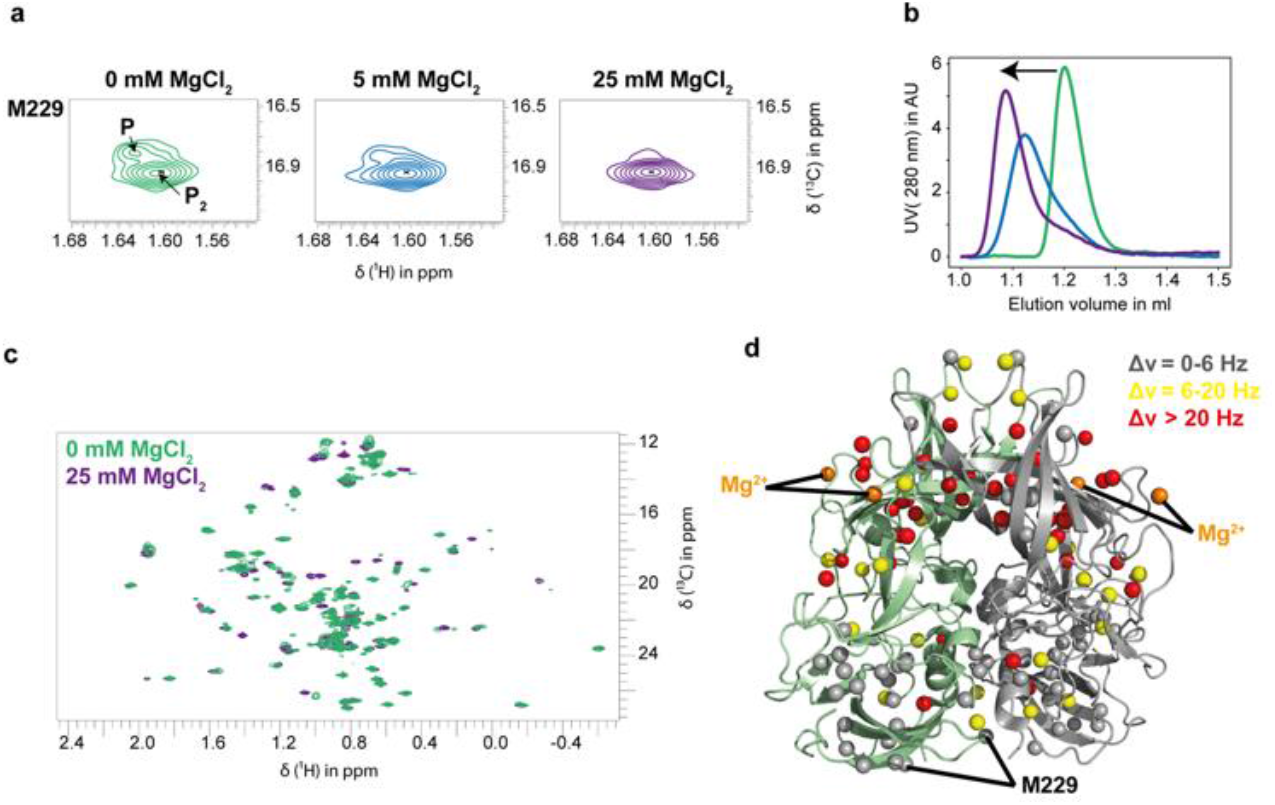
Mg^2+^ cations stabilize MNV P-dimers (P_2_M_2_M′_2_) and cause long range CSPs. (a) Addition of MgCl_2_ leads to the monomeric P-domain (P) cross peak of M229 disappearing in the methyl TROSY spectrum, while the corresponding cross peak of the P-dimer (P_2_ and P_2_M_2_M′_2_) stays. The methyl TROSY spectra were acquired at 298 K using a 75 μM MILVA labeled MNV P-domain sample. (b) The formation of P-dimers upon addition of MgCl_2_ was confirmed using size exclusion chromatography (SEC). The color coding of MgCl_2_ concentrations is as given in (a). The elution profile shifts to the left with increasing concentrations of Mg^2+^ reflecting an increase in molecular weight. (c) Comparison of methyl TROSY spectra in the absence (green) and in the presence of 25 mM MgCl_2_ (violet). (d) CSPs plotted on a structural model^[17]^ (pdb 6e47) of MNV P-dimers. Large CSPs are observed (Δν_Eucl_ > 20 Hz, red) for methyl groups close to the metal ion binding sites, as expected. Many weaker CSPs (6 Hz < Δν_Eucl_ < 20 Hz, yellow) reflect long-range effects. CSPs with Δν_Eucl_ < 6 Hz were not considered as significant. Corresponding CSPs are tabulated in Table S1 as Euclidian distances.

Further analysis of CSPs upon titration with Mg^2+^ or Ca^2+^ revealed long-range effects like those observed upon titration with GCDCA. Comparison of the 62 assigned methyl TROSY resonances of P-dimers (P_2_) with the corresponding resonances of the Mg^2+^ or Ca^2+^ bound P-dimers (P_2_M_2_M′_2_) (Figure 2c) yielded the CSPs listed in Table S1 and visualized in Figure 2d. CSPs were only determined when the cross peaks of the apo and of the bound form were well resolved. In cases where a well-defined cross peak was observed for the bound form but was missing in the corresponding apo form, we assumed that CSPs were larger than 20 Hz. The threshold level for considering CSPs as significant was set at 6 Hz ^[16]^.

To round out the picture, we analyzed the “direction” of long-range CSPs caused by GCDCA and metal ions. This comparison included the lanthanide ion La^3+^, which was used as a diamagnetic reference for measuring ^1^H pseudo contact shifts (PCSs) as will be discussed below. We observed that the CSPs followed linear vectors as this is shown in Figure S3 for ten representative resonance signals of methyl groups adjacent to the G′H′ loop (A442, A444), at the C’D’ loop (V352), in the vicinity of the CD300lf receptor^[34]^ binding site (A365), at the dimer interface (I514, I281), and at other sites (V339, L499, L232). Such linear chemical shift patterns reflect ligand similarity in inducing long-range structural or dynamic protein changes and may identify allosteric networks^[35-37]^.

### Dissecting binding of bivalent metal cations via site-directed mutagenesis of MNV P-domains

The crystal structure of MNV P-dimers complexed with the murine receptor protein CD300lf (pdb 6e47) displays three bound Mg^2+^ ions. One cation binds in the vicinity of the GCDCA binding pocket and makes polar contacts to the side chain carboxyl group of D440 and to the side chain amide CO of Q438. Another Mg^2+^ ion binds to the side chain carboxyl groups of D410 and D366 of two adjacent loops participating in CD300lf recognition. A third Mg^2+^ cation is bound to a loop structure of CD300lf ^[17, 34]^ of a complex between P-dimer and CD300lf (pdb 5ffl and pdb 6c74). This third binding pocket does not exist in the present case where CD300lf is absent. In a previous study we have used the two Mg^2+^ binding sites per P-domain for the assignment of methyl groups in MILVA labeled samples of MNV P-dimers by measuring pseudo contact shifts (PCSs) in the presence of lanthanide ions^[21]^. Our data were consistent with the D440/Q438 binding pocket (Figure S5b) having a higher affinity for metal ions than the D410/D366 binding site. In the present study we dissected metal ion binding to the two sites by deactivating either binding site using site-directed mutagenesis. The corresponding P-domain mutants, D440A and D410A, were subjected to NMR experiments. It turned out that mutation of D440 had a detrimental effect on the appearance of the NMR spectra and resulted in poor P-domain yields. In methyl TROSY spectra of the D440A P-domain many peaks were broadened beyond detection (Figure S4a). In addition, no monomer peaks could be identified for the apo state of the D440A P-domain, and methyl TROSY spectra in the presence and absence of MgCl_2_ showed no major changes. The only assigned resonance clearly showing CSPs is V414 (Figure S4b), which is close to the remaining D410/D366 metal ion binding site. For the D410A P-domain all resonances were well-resolved allowing a detailed analysis of CSPs (Figure S5a) caused by binding of Mg^2+^ to the D440/Q438 site, which is part of the G′H′ loop (Figure S5b). CSPs observed upon adding Mg^2+^ ions to the apo form of the D410A P-domain were very similar to those observed for the wild-type MNV CW1 P-domain (Figure 2). As observed for the wild-type protein, monomer cross peaks disappear in the presence of Mg^2+^ ions reflecting a stabilization of P-dimers (Figures 2a and S5e). This observation shows that the D440/Q438 site is responsible for the dimer-stabilizing effect.

### Binding of bivalent metal cations to the D440/Q438 site of MNV P-domains

For the D410A mutant we could safely exclude binding to the D410/D366 metal cation binding site and the CSPs observed (Figure S6) were solely due to binding to the D440/Q438 binding site in the G′H′ loop (Figure S5b and S5d). We titrated MNV CW1 wild type and D410A mutant P-domains with MgCl_2_ and CaCl_2_ in the absence and in the presence of GCDCA and monitored the resulting CSPs using methyl TROSY spectra. CSPs were measured as shifts of resonance frequencies (Δν_Eucl_) of cross peaks in the case of fast exchange on the chemical shift time scale. In the case of slow exchange, we measured the changes of cross peak intensities to obtain binding isotherms. Alternatively, we performed a 2D line shape analysis using the program TITAN^[19]^.

In the absence of GCDCA we observed exchange that was slow-to-intermediate on the NMR chemical shift time scale (Figure 3a). For cross peaks in slow exchange, we measured the increasing intensities of the metal ion bound form (P_2_M_2_) and obtained corresponding binding isotherms (Figure 3b; see also Figures S7 and S8). Dissociation constants *K*_*D*_ were obtained from fitting the law of mass action to the binding isotherms and yielded values in the low mM range for both cations, Mg^2+^ and Ca^2+^ (Table 1). However, considering that the apo form of P-domains exists as a mixture of P-domain monomers (P) and P-dimers (P_2_) some care must be taken in the analysis. As shown above, addition of Mg^2+^ or Ca^2+^ stabilizes P-dimers and causes long-range CSPs (Figure 2 and Figure S5). Therefore, the changes of cross-peak intensities were not only due to metal ion binding but at the same time reflected other changes within the P-dimers. On the other hand, dissociation of MNV P-dimers is characterized by a dissociation constant *K*_*D*_ of 7 μM ^[16]^, reflecting that at the P-domain concentrations used in our experiments, which were five to ten times the *K*_*D*_, P-domains exist almost exclusively as P-dimers (P_2_). Therefore, it is reasonable to ignore the binding of metal ions to P-domain monomers in our data analysis. In addition, the *K*_*D*_ values for binding of Mg^2+^ or Ca^2+^ turned out to be in the mM range, three orders of magnitude above the dissociation constant for P-dimer (P_2_) dissociation. Therefore, the effects of small inaccuracies in protein concentration due to omission of monomers (P) should not noticeably affect the result of the fitting procedure. Nevertheless, we refer to the dissociation constants for metal cation binding to the apo P-domain as “apparent dissociation constants” *K*_*D,app*_ (Table 1), highlighting the fact that these data reflect more than just metal cation binding. Finally, we attempted a 2D line shape analysis of the corresponding methyl TROSY data sets with TITAN^[19]^, which allowed us to apply a three-state model^[16]^, explicitly considering the monomer-dimer equilibrium (Figure S9). However, only a rather small subset of resonances allowed identification of the three states, and the corresponding cross peaks resulting from P-dimers (P_2_) and P-dimers with metal ions bound (P_2_M_2_) were poorly resolved. As a result, the dissociation rate constants *k*_*off*_ resulting from the line shape analysis were associated with a very large covariance so that these data were not used for further interpretation. On the other hand, the dissociation constants derived from the 2D line shape analysis agreed with the *K*_*D,app*_ values obtained from the fitting of binding isotherms (Figure 3b).

**Table 1.** Dissociation constants *K*_*D*_ for binding of Mg^2+^ and Ca^2+^ to the D440/Q438 site. This site is part of the G′H′ loop of the P-domain of MNV CW1. Data were obtained in the absence and presence of saturating amounts of GCDCA (at pH^*^ 5.3). *K*_*D*_ values in the absence of GCDCA are considered “apparent” because the process of metal ion binding is accompanied by a stabilization of P-dimers and the observation of long-range CSPs (see main text). The dissociation constants were obtained from titrations of P-domains with metal cations, monitored by methyl TROSY experiments. In cases of slow-to-intermediate exchange CSPs were determined as changes of cross peak intensities (I) or 2D line shape analysis (II). In the case of fast exchange CSPs were measured as frequency shifts (Δν_Eucl_) of cross peaks (III).

| P-Domain | Cation | GCDCA | $K_{D,app}$ / mM | Method |
| --- | --- | --- | --- | --- |
| MNV CW1 | $Mg^{2+}$ | + | $0.76 \pm 0.09^a$ | III |
| | | - | $5.7 \pm 1.7^a$ | I |
| MNV CW1 (D410A) | | + | $0.62 \pm 0.04$ | III |
| | | - | $11 \pm 6$ | I |
| | | - | $3.8 \pm 0.2$ | II |
| MNV CW1 | $Ca^{2+}$ | + | 0.138 | Ref. [38] |
| | | - | $7.8 \pm 3^b$ | I |
| MNV CW1 (D410A) | | - | $2.7 \pm 2.4^b$ | I |
| | | - | $1.7 \pm 0.2$ | II |

<sup>a</sup> cf. Figure S13; <sup>b</sup> cf. Figure S14.

**Figure 3.**
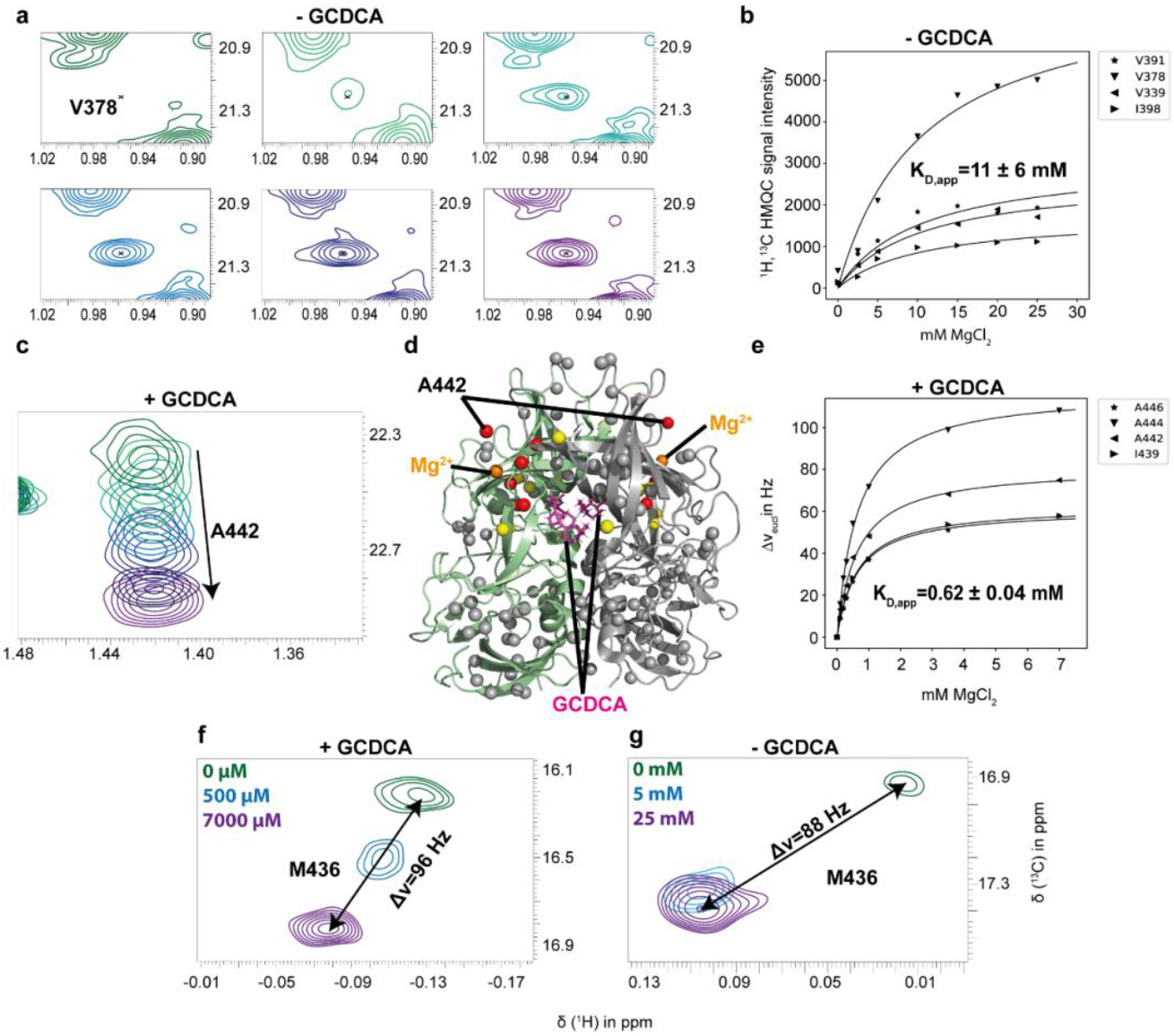
Titrations of the D410A MNV P-domain with Mg^2+^ in the presence and absence of GCDCA. a) Section of a methyl TROSY spectrum of the D410A MNV P-domain in the absence of GCDCA at increasing concentrations of Mg^2+^ (0 to 25 mM, for details see materials and methods), showing the emerging V378 cross peak as an example. The position of this cross peak in the apo-form could not be determined. b) Fitting the law of mass action to the intensities of methyl TROSY cross peaks as a function of the Mg^2+^ concentration yields an apparent dissociation constant *K*_*D,app*_ of 11 mM for the binding of Mg^2+^ in the absence of GCDCA. c-e) In the presence of saturating amounts of GCDCA chemical exchange is fast and allows measurement of CSPs (c) as a function of the Mg^2+^ concentration, yielding an apparent dissociation constant *K*_*D,app*_ of 0.62 mM for the binding of Mg^2+^ in the presence of saturating amounts of GCDCA (e). (d) CSPs observed in the presence of GCDCA plotted on a structural model^[17]^ (pdb 6e47) of MNV P-dimers. For the color coding see the legend to Figure 2. In (c), the peak corresponding to a concentration of 1 mM MgCl_2_ has been omitted for clarity. (f, g) The M436 cross peak under fast (f) and slow (g) chemical exchange conditions was used to estimate upper limits for *k*_*ex*_ as explained in the main text. The methyl TROSY spectra were acquired at 298 K on a 600 MHz spectrometer with cryo probe using a sample of 62 μM MI*LVA labeled D410A MNV CW1 P-domain (see methods section) in the absence of GCDCA (a, b) and a sample of 36 μM MI*LVA labeled D410A MNV CW1 P-domain in the presence of 250 μM GCDCA (c, e). One data point at a concentration of 3.5 mM MgCl_2_ in the presence of GCDCA (c and e) was measured separately using a sample containing 18 instead of 36 μM MI*LVA labeled D410A MNV CW1 P-domain.

Although in this case the 2D line shape analysis failed to provide a reliable value for *k*_*off*_ we were able to estimate an upper limit for the exchange rate constant *k*_*ex*_ for metal ion binding directly from the methyl TROSY spectra. The Euclidian distance between the cross peaks of M436 in the metal-ion bound state and the apo state is 88 Hz (Figure 3g). Since the exchange is slow on the chemical shift time scale, the exchange rate constant *k*_*ex*_ must be substantially smaller than 88 s^-1^ (*k*_*ex*_ << 88 s^-1^). The dissociation rate constant *k*_*off*_ is always smaller than or equal to the exchange rate constant *k*_*ex*_ :

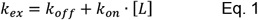

with *k*_*on*_ being the association rate constant, and *[L]* being the ligand concentration, i.e., the Mg^2+^ concentration. Therefore, we can safely assume *k*_*off*_ << 88 s^-1^. Given a dissociation constant *K*_*D*_ in the range between 1 and 10 mM for binding of Mg^2+^ (Table 1) the upper limit for the association rate constant *k*_*on*_ is around 10^4^ M^-1^ s^-1^.

Simpler spectra were obtained from titrations with Mg^2+^ and Ca^2+^ in the presence of saturating amounts of GCDCA, where the P-domains are found exclusively in the dimeric form (P_2_G_2_), with excellent resolution of all methyl TROSY cross peaks. In addition, binding of bivalent metal cations is fast on the chemical shift time scale, allowing the analysis of continuous frequency shifts (Δν_Eucl_) of cross peaks. This procedure is much less error prone than the measurement of cross peak intensities. From Table 1 it is obvious that the dissociation constants for Mg^2+^ and Ca^2+^ are more than a magnitude smaller than in the absence of GCDCA. The values were almost identical for wild type and D410A mutant P-dimers (P_2_G_2_), showing that the presence of the intact D410/D366 metal binding site in the wild type P-dimers did not affect the analysis. Since the system was in the fast exchange regime, we estimated the association rate constant for the binding of Mg^2+^ as above using the Euclidian distance between the cross peaks of M436 of the D410A mutant P-domain in the GCDCA saturated state (P_2_G_2_) and the corresponding metal ion bound state (P_2_G_2_M_2_) as a measure. The cross peak of M436 shifts with increasing Mg^2+^ concentration yielding a maximum value of Δν_Eucl_ of 96 Hz (Figure 3f), which in this case serves as a lower limit for the exchange rate constant *k*_*ex*_. A statement concerning the association rate constant *k*_*on*_ is less straightforward than in the absence of GCDCA since Eq. 1 yields a lower limit for *k*_*on*_ that depends on the ligand concentration, i.e., the Mg^2+^ concentration:

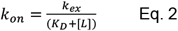

Using a *K*_*D*_ value of 0.62 mM (Table 1) and assuming Mg^2+^ concentrations ranging from 0 to 7 mM (Figure 3f), the association rate constant *k*_*on*_ is estimated to be larger than 10^4^ to 10^5^ M^-1^ s^-1^.

### Binding of bivalent metal cations to the D410/D366 site of MNV P-domains

Binding affinities for binding of Mg^2+^ and Ca^2+^ to the D410/D366 site, which is very close to the CD300lf receptor binding site were obtained from methyl TROSY titration experiments, employing the wild type P-domain and its D440A mutant. Of the assigned cross peaks of the mutant only the resonance of V414 showed unambiguous CSPs. We used the corresponding methyl TROSY cross peak to obtain a binding isotherm from a titration with Mg^2+^ (Figure S4). The binding reaction was fast on the chemical shift scale and a dissociation constant in the high mM range was obtained (Table 2). In the presence of GCDCA the quality of the methyl TROSY spectrum of the D440A mutant P-domain improved dramatically (Figure S10). For the metal ion titration experiments we used the methyl TROSY cross peaks of the three amino acids, V414, A365, and I358 that were in the vicinity of the D410/D366 binding site, and that were unaffected by metal ion binding to the D440/Q438 site allowing to also use the wild type P-domain for titration experiments (Figure S10). All dissociation constants were in the high mM range, and we could not detect an effect of GCDCA binding on metal ion binding to this site.

**Table 2.**
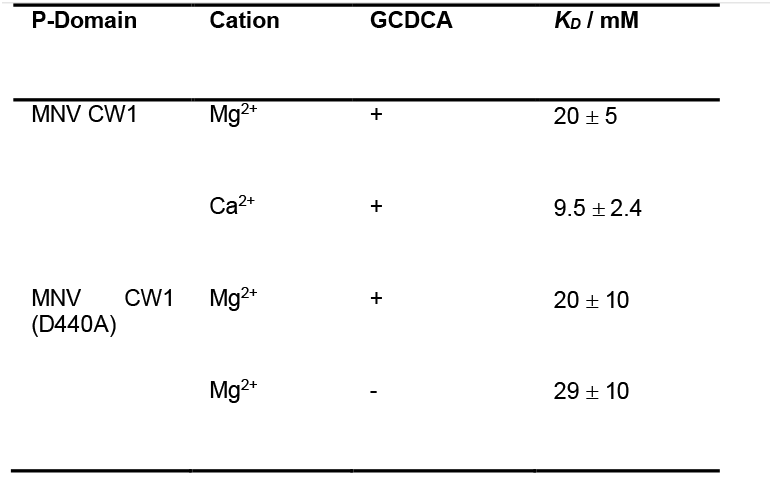
Dissociation constants *K*_*D*_ for binding of Mg^2+^ and Ca^2+^ to the D410/D366 site. This site is close to the CD300lf receptor binding site. Data were obtained in the absence and presence of saturating amounts of GCDCA (at pH^*^ 5.3). *K*_*D*_ values in the absence of GCDCA are considered as apparent (cf. legend to Table 1). CSPs were measured as frequency shifts (**Δν**_Eucl_) to obtain binding isotherms. For the titrations in the presence of GCDCA the resonances of V414, A365 and I358 were followed (cf. Figures S4 and S10). For the titration of the apo form of the D440A mutant P-domain with Mg^2+^ only the V414 resonance was used.

| P-Domain | Cation | GCDCA | $K_D$ / mM |
| --- | --- | --- | --- |
| MNV CW1 | $Mg^{2+}$ | + | $20 \pm 5$ |
| | $Ca^{2+}$ | + | $9.5 \pm 2.4$ |
| MNV CW1 (D440A) | $Mg^{2+}$ | + | $20 \pm 10$ |
| | $Mg^{2+}$ | - | $29 \pm 10$ |

### Binding of GCDCA to MNV P-dimers in the presence of bivalent metal cations

Next, we studied the effects of binding of Mg^2+^ and Ca^2+^ on the binding of GCDCA to MNV P-domains. In our previous study we performed 2D line shape analysis to determine the dissociation constant *K*_*D*_ as well as the dissociation rate constant *k*_*off*_ for binding of GCDCA in the absence of bivalent metal ions. Here, we have repeated these experiments in the presence of saturating concentrations of Mg^2+^ and Ca^2+^. We identified nine spectral regions in the methyl TROSY spectrum of MILVA-labeled wild type MNV P-dimers saturated with Mg^2+^ or Ca^2+^ that were suitable for 2D line shape analysis (Figure 4), using a two-state binding model to yield corresponding *K*_*D*_ and *k*_*off*_ values (Table 3). Obviously, Mg^2+^ or Ca^2+^ ions increase the affinity of P-dimers for GCDCA by slowing down dissociation and accelerating binding of GCDCA. The effect is more pronounced for Ca^2+^ resulting in a dissociation constant two orders of magnitude smaller than in the absence of any bivalent metal cations.

**Table 3:**
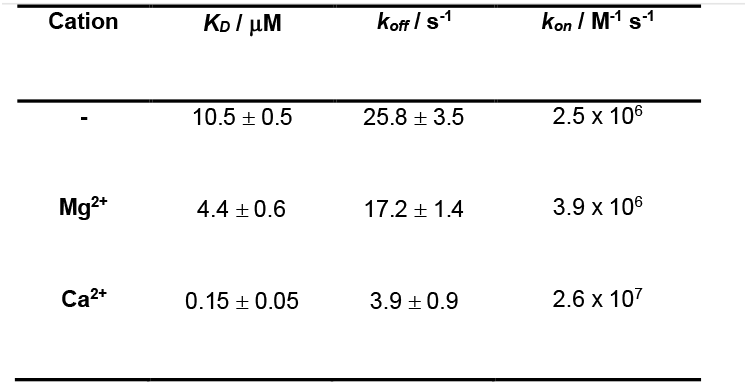
Dissociation constants *K*_*D*_ and dissociation rate constants *k*_*off*_ for binding of GCDCA to the MNV CW1 P-domain. The data were obtained using 2D line shape analysis in the absence and presence (25 mM) of bivalent metal cations at pH^*^ 5.3. The spectral regions used for the line shape analysis are shown in Figure 4. The data in the absence of bivalent metal ions were taken from our previous study^**[16]**^.

**Figure 4:**
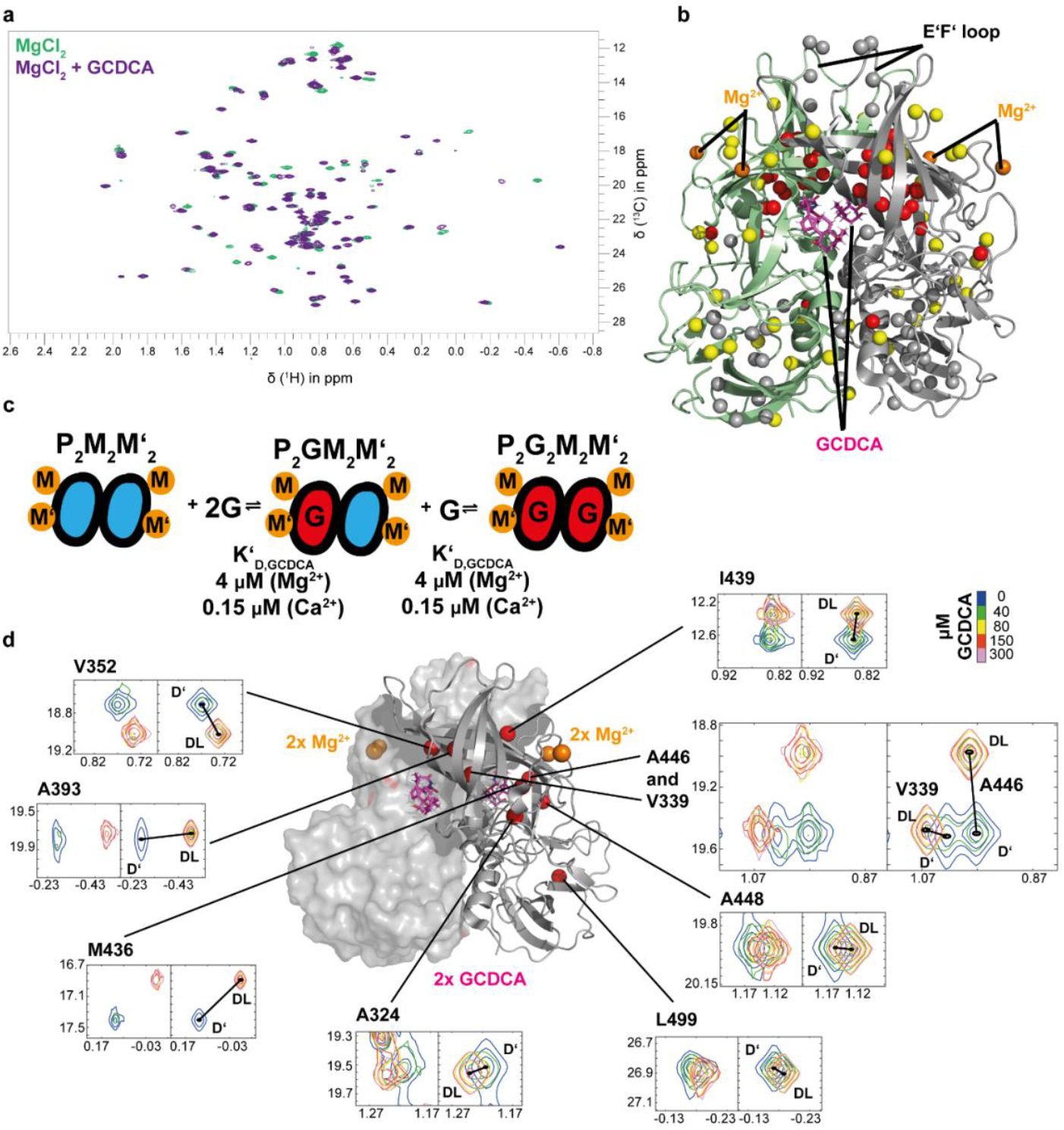
2D line shape analysis of methyl TROSY cross peaks of MNV P-dimers titrated with GCDCA in the presence of Mg^2+^. (a) Comparison of methyl TROSY spectra of MILVA-labeled MNV CW1 P-dimers (75 μM) saturated with Mg^2+^ (25 mM MgCl_2_) in the absence and presence of GCDCA. (b) Mapping of CSPs on a structural model from crystal structure analysis^[17]^ (pdb 6e47). The color coding is identical with the coding used in Figure 2d. No long-range CSPs are observed in the E’F’ loop. (c) Methyl TROSY cross peaks used for 2D line shape analysis with TITAN and the location of the corresponding amino acids in the structural model. For the line shape analysis, a simple two-state model was used, assuming saturation with Mg^2+^ already had stabilized P-dimers (cf. Figures 2 and 3) with two equivalent GCDCA binding pockets (cf. Figure 1a).

In our previous study we reported many long-range CSPs upon binding of GCDCA to MNV CW1 P-dimers in the absence of bivalent metal cations (cf. Fig. 4 of ref.^[16]^). A subset of these long-range effects was observed in the E’F’ loop, for which a cryo-EM study had shown that neutralizing antibodies can only bind to the so-called open conformation of this loop^[3]^, i.e., in the absence of GCDCA. In the presence of saturating amounts of Mg^2+^ or Ca^2+^ we also observed many long-range CSPs upon addition of GCDCA (Mg^2+^: Figure 4b; Ca^2+^: Figure S11). However, no CSPs were observed within the E’F’ loop.

### Dimerization of MNV P-domains and metal ion binding depend on pH

Methyl TROSY spectra were recorded at different pH values. We report pH readings in D_2_O solutions as pH* values. To compare to data from other studies, usually performed in H_2_O solution, we added corrected pH values, pH_corr_^[39]^. We measured the intensity ratios of monomer to dimer cross peaks, which reflect the degree of dimerization. The pH dependence of the intensity ratios (Figure S12a) indicated that dimer stability is optimal at pH values between 4 and 5. At more acidic or basic pH values the intensities of monomer cross peaks increased, reflecting diminished dimer stability. SEC runs reflected a lower apparent molecular weight at higher pH values (Figure S12b), supporting the NMR observations. In our previous study^[16]^ we performed a 2D line shape analysis of methyl TROSY spectra of samples with increasing MNV P-dimer concentrations under slightly acidic conditions, at pH^*^ 5.3 (pH_corr_ 5.3). The analysis yielded a dissociation constant *K*_*D*_ of 7 μM and a dissociation rate constant *k*_*off*_ of 1.2 s^-1^. Here, we repeated these experiments at pH_corr_ 7.4 (pH* 7.5). A dissociation constant *K*_*D*_ of 45 μM and a dissociation rate constant *k*_*off*_ of 4.7 s^-1^ reflect the reduced stability of P-dimers under near-neutral conditions.

We then tested the binding of Ca^2+^ ions to the MNV P-domain at pH* 3.9 (pH_corr_ 4.0) and at pH* 7.5 (pH_corr_ 7.4). At pH* 3.9 (pH_corr_ 4.0) no CSPs associated with binding to the D440 site were observed upon titration of P-domain samples with Ca^2+^ (Figure S13a). At pH* 7.4, however, we were able to identify cross peaks of P-domain monomers, of P-dimers, and of P-dimers with Ca^2+^ bound (Figure S13b), allowing to perform a TITAN line shape analysis employing a three-state model (Figures S13c and S13d). For Ca^2+^-binding we obtained a dissociation constant of *K*_*D*_ = 1.09 ± 0.02 mM and a dissociation rate constant of *k*_*off*_ = 27.5 ± 0.4 s^-1^.

Finally, we asked whether we could analyze the protonation state of D440, which is in the G′H′ loop and is essential for metal ion chelation. We monitored the ^1^H and ^13^C NMR chemical shifts of alanine residues A442, A444, and A446 as a function of the pH (Figure S14). These amino acids are in the same G′H′ loop as D440 and should be indirectly affected by the protonation state of D440. Apart from D440 there are two other acidic amino acid residues, D443 and E447 in the G′H′ loop. Therefore, we could not attribute the effect of pH solely to the protonation state of D440, but rather observed a bulk effect, which explained the observation that the chemical shifts did not follow straight vectors, but rather curved lines. Nevertheless, we obtained apparent *pK*_*a*_ values that indicated a change of the protonation status of the said amino acids between pH 4 and 6.

### Virus binding assays reveal the synergistic interplay between Mg^2+^, Ca^2+^, and GCDCA binding to MNV virions

The observation that Mg^2+^ or Ca^2+^ binding and binding of GCDCA are synergistic raised the question of the biological impact of this finding (Figure 5). In the absence or at low (1 mM) ion concentration, the bile acid GCDCA only marginally blocked antibody binding (Figures 5a and 5b). However, high concentrations of the bivalent cations Mg^2+^ or Ca^2+^ (50 mM) strongly inhibited antibody binding in the presence of bile acid. Sodium ions did not affect antibody binding at any concentration tested, and no synergistic effect with bile acid was observed here (Figure 5c). A dose-dependent synergistic effect on the loss of antibody binding was observed at ion concentrations ranging between 1– 100 mM. The lowest magnesium concentration that showed an effect on antibody binding in the absence of GCDCA was ∼50 mM (Figure 5d). To address which ion-binding site (D410 or D440) plays a role, recombinant virions were generated by reverse genetics with the ion-binding position D410 or D440 mutated to alanine. The ELISA with the recombinant virions, showed that the D410 position did not impact antibody escape while losing the ion-binding site at the D440 position prevented the synergistic effect. In conclusion, both Ca^2+^ and Mg^2+^ blocked antibody binding to MNV in a dose-dependent manner, indicating that this is highly biologically relevant. On a molecular level, GCDCA and Mg^2+^ or Ca^2+^ decrease the affinity of a neutralizing antibody to the virion, correlating with the observed shift of the conformational equilibrium of the E’F’-loop towards the closed-loop conformation.

**Figure 5:**
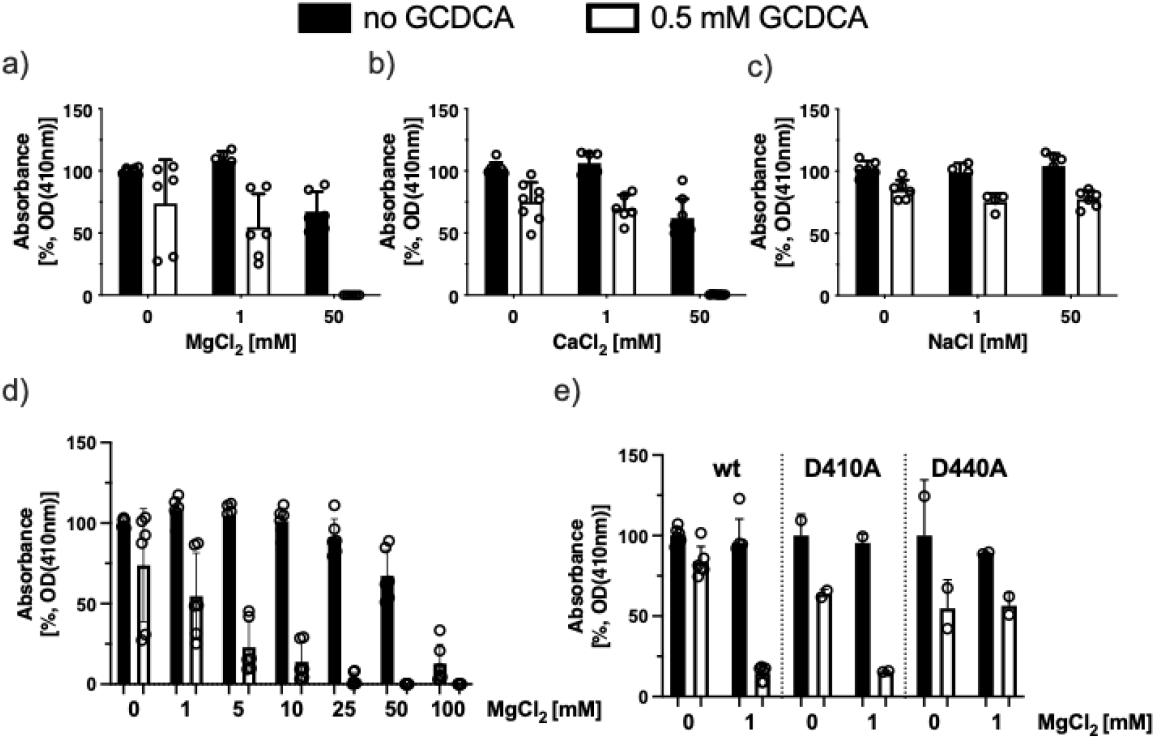
The bile acid GCDCA synergistically acts with bivalent cations preventing murine norovirus (MNV-1) virions from binding to a neutralizing antibody. An ELISAs was performed under defined ion conditions, with specific metal ions Mg^2+^, Ca^2+^, or Na^+^ ions and bile acid GCDCA added to coated infectious MNV virions during antibody (A6.2) binding (Figure 5a-c). Antibody binding was determined in the absence and presence of 0.5 mM GCDCA. ELISA data was normalized to no-ion and non-GCDCA treatment and shown a mean of three independent experiments with error bars representing the standard deviation (SD).

### Probing for conformational changes of MNV P-domains upon metal ion binding

The long-range CSPs observed in methyl TROSY spectra upon addition of bivalent metal ions (see Figures 2 and 3) indicate conformational changes. It is known that the conformational changes associated with such long-range CSPs can be very subtle. To test whether the CSPs reflect only subtle local changes, such as reorientation of amino acid side chains, or are associated with larger structural rearrangements, we measured ^1^H pseudo-contact shifts (PCSs) caused by lanthanide ions. We have previously shown that lanthanide ions bind to the higher affinity metal ion binding site and used PCSs in conjunction with crystal structure data to extend the ^13^C methyl group assignment of MNV P-dimers saturated with GCDCA^[21]^. PCSs depend directly on the distance and orientation of a methyl group relative to the paramagnetic lanthanide ion and are very sensitive to structural changes. Therefore, we measured ^1^H PCSs to analyze the effect of metal ion binding on the conformation of MNV P-dimers. A methyl TROSY spectrum of MNV P-dimers in the presence of La^3+^ is quite like that in the presence of Mg^2+^ (Figure S15), documenting that the effects of lanthanide ion binding are comparable to those of Mg^2+^ or Ca^2+^ binding. To define the direction of the PCS vectors connecting the cross peaks in the presence of the diamagnetic reference La^3+^ and in the presence of paramagnetic lanthanide ions, we used Sa^3+^, which causes only small PCSs and is easy to trace back to the diamagnetic reference. The PCSs were then measured in the presence of Ce^3+^, which causes much larger PCSs.

First, we correlated ^1^H PCSs in the absence and presence of GCDCA yielding an almost perfect linear correlation (Figure S16). Outliers were only found for methyl groups close to the GCDCA binding site and close to the metal ion binding site in the G′H′ loop. These effects were expected as GCDCA binding may easily cause reorientations of amino acid side chains in the G′H′ loop. Second, we compared the experimental PCSs with theoretical PCSs predicted from three different X-ray and cryo-EM structures. Theoretical PCSs were calculated using the Paramagpy program^[40]^, as described in detail previously^[41]^. The differences between experimental and theoretical values were quantified using the Q-factor, which takes values between 0 and 1. A Q-factor of 0 indicates perfect agreement between experimental and theoretical values, and a Q-factor of 1 indicates no agreement. For comparison, we used a crystal structure of the apo-form of the MNV P domain (PDB 3lq6), a cryo-EM structure of the MNV P domain complexed with a monoclonal antibody binding to the E’F’ loop (PDB 7l5j), and a crystal structure of the MNV P domain in the presence of Mg^2+^ and GCDCA and complexed with the CD300lf receptor (PDB 6e47). The main difference between the structures is the position of the E’F’ loop, which is in the “open” conformation in the first two structures and in the “closed” conformation in the receptor-bound P-domain. The correlations between experimental and theoretical PCSs are shown in Figure S17. The Q-factor is reasonable for the two “open” conformations and is much better for the “closed” conformation (Figures S17 and Table S2). Interestingly, calculated and experimental PCSs for A382, which resides in the E’F’ loop, agree only for the “closed” conformation.

### Changes of methyl side chain mobilities of MNV P-domains upon GCDCA binding

Finally, we asked how the side chain mobility is affected by GCDCA binding. To this end, we applied a semi-quantitative approach that utilizes the ratios of cross peak intensities in HMQC and in HSQC spectra to estimate methyl side chain order in perdeuterated ^13^C-methyl labeled samples of large proteins^[42]^. It has been shown^[42]^ that for large proteins this approach leads to methyl 3-fold axis order parameters, *S*^*2*^_*axis*_, closely matching order parameters obtained from the measurement of buildup of “forbidden” ^1^H double or triple quantum magnetization^[43]^. We performed this analysis for MNV P-domains in the absence and presence of GCDCA as this is described in detail in the supporting information. The comparison of methyl group order parameters of MNV P-dimers with and without GCDCA bound showed that binding of GCDCA leads to an overall decrease of *S*^*2*^_*axis*_ values. Only few methyl groups show an increase of *S*^*2*^_*axis*_ (Figure S18).

### Bivalent metal ions bind to two prototypic human genogroup II norovirus P-domains

We examined the binding of bivalent metal ions to P-dimers of two prototypic genogroup II human norovirus (HuNoV) strains, GII.4 Saga and GII.17 Kawasaki. We had previously studied the P-dimers of the GII.4 Saga strain in detail by NMR^[14, 16, 20-21, 38, 44-46]^. An important finding was that N373 in wild-type GII.4 Saga P-dimers undergoes spontaneous and fast deamidation^[14, 44]^. Therefore, here we used the stable N373D mutant^[38]^ instead of the wild type P-dimers for all NMR binding experiments. All results reported in the following have been obtained using MILVA-labeled N373D Saga P-dimers. The ^13^C-methyl assignment obtained previously for wild-type Saga P-dimers^[20]^ was straightforwardly transferred to the N373D mutant. MILVA-labeled Kawasaki P-dimers were used without NMR signal assignments.

Titrations of N373D GII.4 Saga P-dimers with Mg^2+^ or Ca^2+^ ions caused no CSPs in methyl TROSY spectra, showing that these ions do not bind under these conditions (Figure S20). For GII.17 Kawasaki P-dimers we obtained the same result.

Upon addition of Zn^2+^ to Saga P-dimers we observed CSPs as well as line-broadening and resulting signal intensity reduction for specific resonances (Figure 6), reflecting binding. The effects are partially reversible upon addition of EDTA (Figures 6a and 6b). A plot of signal intensity reductions at a concentration of 800 μM Zn^2+^ as a function of amino acid sequence clearly shows that the reduction of signal intensities is site specific (Figure 6c), matching the corresponding bar graph for the amino acid specific increase of transverse relaxation rates *R*_*2*_ (Figure 3 d). The perturbations can be mapped to the P-dimer surface as this is also shown in Figures 6c and 6d. Along with the reduction of signal intensities we observed CSPs (Figures 6e and 6f). We performed a titration with Zn^2+^ ions to extract dissociation constants *K*_*D*_ from concentration dependent CSPs. At a maximum concentration of 3 mM Zn^2+^ we observed a significant overall loss of signal intensities (Figure 6b), most likely due to aggregation and partial unfolding. Yet, the loss of signal intensity remained site specific. Despite this site-specific loss of signal intensities, we could follow CSPs for several amino acids (Figure 6e) located in two distinct regions *1* and *2* (Figure 3f). Global fitting of the law of mass action to the resulting binding isotherms provided two dissociation constants *K*_*D*_ of 3.5 mM (for region *1* in Figure 6f) and 1 mM (for region *2* in Figure 6f). After reaching the maximum concentration of 3 mM Zn^2+^ we added EDTA to reverse the effects of signal reduction. The original spectral appearance could not be fully restored, and some additional cross peaks remained, possibly due to irreversible partial unfolding (Figure 6a).

**Figure 6:**
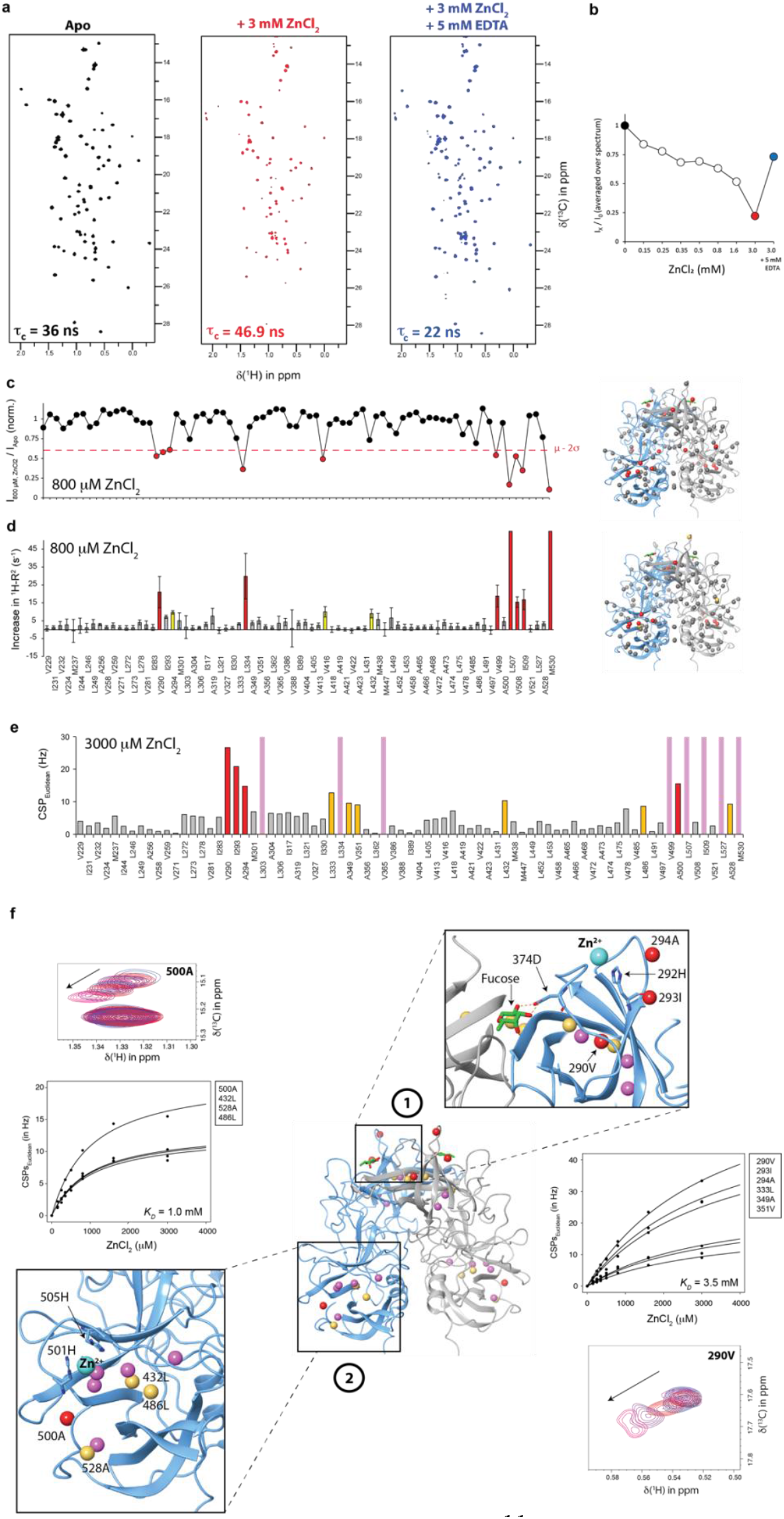
Addition of ZnCl_2_ causes perturbations in methyl TROSY spectra of MILVA-labeled N373D HuNoV GII.4 Saga P-dimers. a) Methyl TROSY spectra of Saga P-dimers in the absence of ZnCl_2_ (black), in the presence of 3 mM ZnCl_2_ (red), and upon addition of 5 mM EDTA (blue). ZnCl_2_ leads to a strong signal decrease along with an increase in τ_c_ (cf. Figure S5), suggesting protein aggregation. Signal intensities can be partially recovered by addition of 5 mM EDTA (blue). However, the appearance of new peaks and a strong decrease of τ_c_ as compared to apo P-dimers suggests partial protein unfolding. τ_c_ values were obtained from ^15^N TRACT experiments (Figure S5). b) Normalized sum of intensities of all cross peaks of methyl TROSY spectra of Saga P-dimers at increasing ZnCl_2_ concentrations and, finally, after the addition of 5 mM EDTA. c) *Left panel*: Methyl group specific decrease of methyl group signal intensities in a methyl TROSY spectrum of Saga P-dimers in the presence of 800 mM ZnCl_2_. Red dots indicate a signal decrease of at least μ-2σ. *Right panel:* Signal intensity decrease mapped onto the crystal structure of Saga P-dimers (pdb 4X06). d) *Left panel*: Increase of ^1^H_M_-Г_2_ rates of methyl groups in Saga P-dimers at 800 mM ZnCl_2_. Yellow and red bars indicate an increase of μ+σ and μ+2σ, respectively (for the calculation of errors see the Methods section). *Right panel*: ^1^H_M_-Г_2_ mapped onto the crystal structure of GII.4 Saga P-dimers (pdb 4X06). e) Methyl group CSPs observed at 3 mM ZnCl_2_. Methyl signals that were broadened beyond detection are shown in pink. Orange and red denote CSPs larger than μ+σ and μ+2σ, respectively. Gray indicates non-significant CSPs. f) CSPs at 3 mM ZnCl_2_ mapped onto the crystal structure (pdb 4X06) of Saga P-dimers (color code as in e)). Fitting the law of mass action to the experimental binding isotherms yields dissociation constants *K*_*D*_ for the binding of Zn^2+^ within the binding regions *1* and *2*. Amino acid labels next to the binding isotherms are sorted from the largest to the smallest CSPs at 3 mM ZnCl_2_. CSPs as a function of ZnCl_2_ concentration a shown for A500 and V290 as examples. The positions of the metal ions are only to be understood symbolically. For binding region *1* the cryo-EM structure of Jung et al.^[18]^ (pdb 6OUC) served as a guide, and for region *2* the position was chosen manually to illustrate H501 and H505 as the potentially chelating amino acids.

TRACT experiments yielded overall motional correlation times *τ*_*c*_ for the apo form of Saga P-dimers at the start of the titration, for the conditions found at 3 mM Zn^2+^, and for the state after adding EDTA (Figure S21). These values match well with the data that we reported previously^[14]^, where we found that *τ*_*c*_ depends strongly on the P-dimer concentration, the pH and the overall salt concentration. In general, a decrease of the pH and an increase of the overall salt concentration caused an increase of *τ*_*c*_ (cf. Figure S6 of ref.^[14]^). The increase at P-dimer concentrations above ca. 100 μM was due to protein aggregation. Under the conditions chosen here, at a P-dimer concentration of 30 μM and a pH of 6.9 aggregation can be excluded. For the apo-form we measured a value of *τ*_*c*_ = 36 ns, as expected. A value of *τ*_*c*_ = 47 ns was obtained in the presence of 3 mM ZnCl_2_, suggesting Zn^2+^-induced aggregation. This matches the observation of decreasing overall signal intensities in methyl TROSY spectra upon addition of Zn^2+^ (Figure 6a). Upon addition of EDTA, TRACT experiments yielded a value of *τ*_*c*_ = 22 ns (Figure S21 and Figure 6a), slightly lower than the value obtained for the apo-form in the beginning, and likely reflecting the presence of irreversibly misfolded species with different motional properties (see above). A very similar picture unfolded upon addition of Cd^2+^ to Saga P-dimers (Figure S22). The same two binding regions *1* and *2* were identified with somewhat lower dissociation constants *K*_*D*_ of 0.3 and 2.5 mM, respectively (Figure S22e). For Cd^2+^ a third binding region *3* with a dissociation constant of 1.3 mM was identified (Figure S22e).

We then tested whether Zn^2+^ or Cd^2+^ affected the attachment of histo blood group antigens (HBGAs) to P-dimers, which is thought to be essential for viral infection. Specifically, we tested binding of methyl-α-L-fucopyranoside, which constitutes the minimal binding element of HBGAs^[47]^, to Saga P-dimers in the presence and absence of Zn^2+^ or Cd^2+^ (Figure S23). The experiments showed that the presence of either of the two metal ions slightly improved the binding affinity of methyl-α-L-fucopyranoside up to a factor of two.

Finally, we tested for binding of the transition metal ions Cr^3+^, Mn^2+^, Fe^2+^, Co^2+^, and Cu^2+^ as well as of the lanthanide ions La^3+^, Lu^3+^, Eu^3+^, and Ce^3+^. The paramagnetic Cr^3+^ and Fe^2+^ ions had no effect on the methyl TROSY spectra (Figure S20). For the paramagnetic metal ions Mn^2+^, Co^2+^, Cu^2+^, Eu^3+^, and Ce^3+^ we observed paramagnetic relaxation enhancements (PREs) that were quantified as a reduction of methyl TROSY cross-peak intensities or an increase of ^1^H relaxation rates Г_2_ (Figures S24 and S25). The diamagnetic lanthanides La^3+^ and Lu^3+^ caused CSPs. It turns out that all these metal ions except for the paramagnetic Cu^2+^ and the diamagnetic Lu^3+^ bind within binding region *3* (Figure S24). For Cu^2+^ we observed PREs and CSPs at the same time (Figure S25). The perturbations caused by Cu^2+^ indicated binding within a region adjacent to the HBGA site (region *1*).

In the case of Cd^2+^ and Cu^2+^, the quality of the methyl TROSY spectra allowed direct simulation with TITAN yielding kinetic rate constants *k*_*on*_ and *k*_*off*_ for the binding of these metal ions (Table S3).

## Discussion

The binding of bivalent metal ions to MNV and HuNoV P-domains has been studied by following changes of the positions, shapes, and intensities of NMR resonance lines in methyl TROSY spectra. In some cases, we also measured the effects of metal ion binding on PCSs, transverse relaxation rates, and on ^13^C methyl group order parameters. The datasets revealed metal ion binding regions as well as thermodynamic and kinetic binding parameters, leading to a detailed picture for the binding of bivalent metal ions to the P-domains of human and murine noroviruses, extending our current structural models.

For MNV P-domains the NMR binding experiments allowed the determination of dissociation constants, and under favorable conditions also of dissociation rate constants. The NMR experiments show that binding of Mg^2+^ or Ca^2+^ to the D440/Q438 site of MNV P-dimers is highly synergistic with GCDCA binding. At the same time the dimeric state of MNV P-domains is synergistically stabilized by these co-factors. In addition, the dissociation of MNV P-dimers strongly depends on the pH, with an optimum stability in the range of pH 4-5. In the absence of any co-factors MNV P-domains exist as a mixture of monomers and dimers. Depending on the presence of the co-factors Mg^2+^, Ca^2+^, or GCDCA we observed all possible cases of chemical exchange, ranging from fast to intermediate to slow exchange (cf. Figure 1b), offering different options for the analysis of methyl TROSY titration experiments. Cases where a 2D lineshape analysis with TITAN was possible yielded affinities as well as dissociation rate constants for the binding of co-factors (cf. Tables 1-3). In the fast exchange regime, dissociation constants were obtained from CSP titrations, and for slow chemical exchange we measured respective peak intensities. Depending on the experimental conditions, we were facing a varying degree of spectral complexity. For example, when titrating metal ions to the apo-form of the P-dimers at pH* 5.3 it was impossible to dissect dimerization and binding, yielding so-called apparent dissociation constants *K*_*D,app*_ (cf. Table 1 and Figure 3). In contrast, titration of P-dimers in the presence of saturating concentrations of bivalent metal ions with GCDCA allowed straightforward 2D-lineshape analysis based on a two-state model (cf. Figure 4) and yielded dissociation constants *K*_*D*_ as well as dissociation rate constants *k*_*off*_ for binding of GCDCA to P-dimers (cf. Table 3).

Although some of the equilibria examined could not be fully analyzed due to experimental constraints, we obtained a complete thermodynamic cycle with changes of the Gibbs free energy ΔG for each individual step starting from MNV P-domain monomers and leading to MNV P-dimers saturated with GCDCA and Mg^2+^ or Ca^2+^, with kinetic parameters annotated where available (Figure 7).

**Figure 7:**
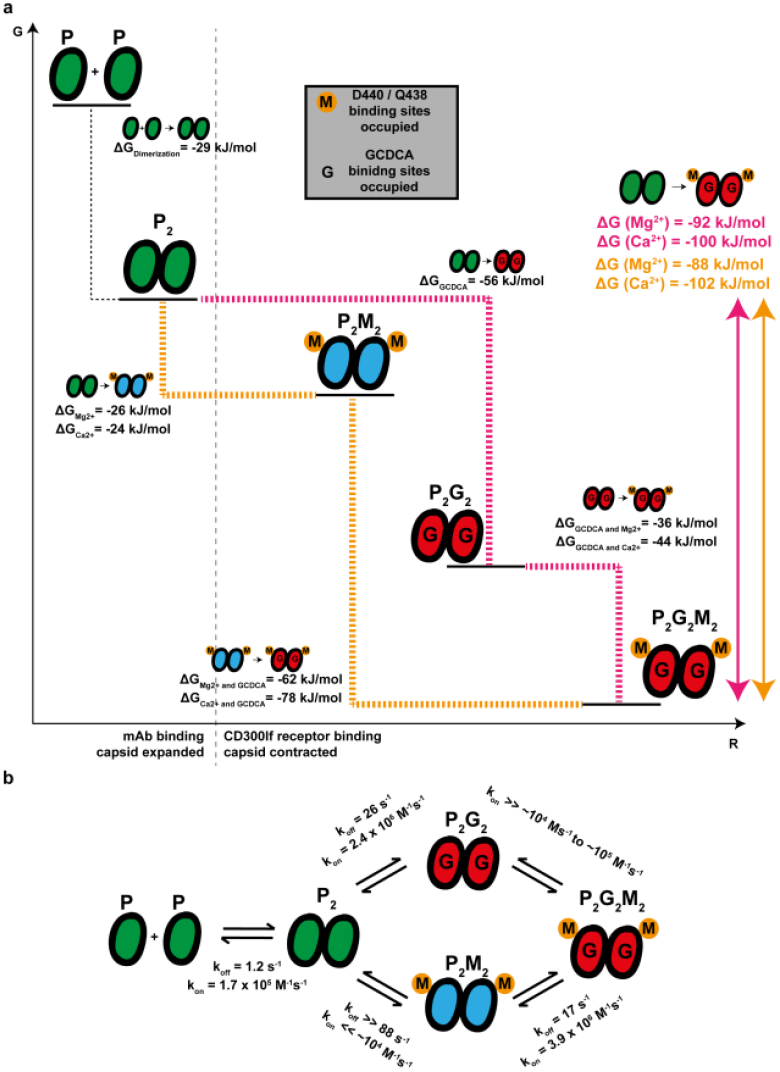
a) Thermodynamic cycle for the dimerization of MNV P-domains and for binding of GCDCA, Mg^2+^, and Ca^2+^ (pH* 5.3). Dissociation constants *K*_*D*_ are converted into changes of free energy ΔG. The abscissa labeled with “R” symbolizes the progression of complex formation. Dashed lines denote addition of GCDCA first and Mg^2+^ or Ca^2+^ subsequently (magenta) or the inverse order (orange). Overall ΔG values for either path are in magenta or in orange, corresponding to the order of addition. The vertical black dashed line separates the “open” conformation of the E’F’ loop from the “closed” conformation, which is associated with capsid contraction. b) Kinetic rate constants *k*_*on*_ and *k*_*off*_ for the dimerization of MNV P-Domains and for binding of Mg^2+^ and GCDCA.

Our experiments show that the first step (Figure 7b), the dimerization of MNV P-domain monomers, strongly depends on the pH (cf. Figure S12). Optimal dimer stability is found in the range of pH 4-5. For the dimerization of apo P-domains we were able to determine dissociation constants *K*_*D*_ and dissociation rate constants *k*_*off*_ under slightly acidic (pH* 5.3)^[16]^ and under near-neutral conditions (pH* 7.4) (Figure S12). The dissociation constant at pH* 7.4 (*K*_*D*_ = 45 μM) is 6.5-fold larger than at pH 5.3* (*K*_*D*_ = 7 μM). This decrease of dimer-stability is mainly due to an increase of the dissociation rate constant, which is four times higher at pH* 7.4 (*k*_*off*_ = 4.7 s^-1^) than at pH* 5.3 (*k*_*off*_ = 1.2 s^-1^). It should be noted that at an acidic pH below about 4 we were not able to detect Ca^2+^ binding to the higher affinity metal ion binding site (see Figure S13), which means that acidic pH alone has a dimer-stabilizing effect. Whether there is a synergism between pH and metal ion binding cannot be determined from the present data. The free energy diagram in Figure 7 reflects that GCDCA as well as bivalent metal ions stabilize MNV P-dimers. Of note, only the D440/Q438 metal ion binding site (cf. Figure 3d) is responsible for the stabilization. This parallels the synergism between metal ion binding to the D440/Q438 site and GCDCA binding.

The findings that apo MNV P-dimers show optimum stability in a pH window between 4 and 5 and that binding of GCDCA and Ca^2+^ and/or Mg^2+^ significantly stabilize P-dimers are correlated with the overall stability and/or shape of MNV capsids. In fact, cryo-EM studies have shown that each of the three co-factors, GCDCA, bivalent metal ions and acidic pH causes an overall contraction of virus capsids, associated with a reorientation of the E’F’-loop such that recognition by neutralizing antibodies is impeded whereas binding to the CD300lf receptor is promoted^[1-3, 5, 48]^. Our data from virus neutralization assays (cf. Figure 5) now directly exhibit that the crosstalk between binding of metal ions and GCDCA to MNV virions has a synergistic impact on virus neutralization by neutralizing mAbs. At the time of writing of this manuscript, another study has been published^[48]^ that also reports a synergistic effect of the binding of Ca^2+^ on the binding of GCDCA to MNV P-domains. The values have been obtained using isothermal titration calorimetry (ITC) at a pH of 7.4 where the equilibrium between P-domain monomers and dimers is shifted towards P-domain monomers (ca. 70% monomers, cf. Figure S12). At the concentration of Ca^2+^ ions of 2 mM used in that study an equilibrium mixture of monomers, dimers, and dimers with Ca^2+^ bound is expected. Therefore, the heats measured upon addition of GCDCA reflect the cumulative effect of binding of GCDCA and dimerization. Here, we have chosen conditions where the metal ion binding pockets are saturated with Ca^2+^ or Mg^2+^, and the P-domains almost exclusively exist as dimers. Thus, the respective dissociation constants (Table 3) reflect binding of GCDCA without any contribution from the dimerization process. Binding of GCDCA in the absence of any bivalent metal ions always carries a contribution from P-domain dimerization. At a pH* of 5.3 chosen for our studies more than 70% of the P-domains are found as dimers (Figure S12) and dimerization contributes less. As a matter of fact, in our previous study^[16]^ we had explicitly taken into account the coupling of binding of GCDCA and dimerization of P-domains, applying a three state model (cf. Fig. 5 of ref.^[16]^). Therefore, the resulting dissociation constant *K*_*D*_, reported here in Table 3, exclusively reflects binding of GCDCA and we can conclude that the presence of Ca^2+^ ions reduce the *K*_*D*_ for binding of GCDCA by almost two orders of magnitude (Table 3 and Figure 8). Interestingly, the effect of Mg^2+^ is much weaker, suggesting that *in vivo* the ratio of Ca^2+^ to Mg^2+^ may fine-tune infectivity.

**Figure 8:**
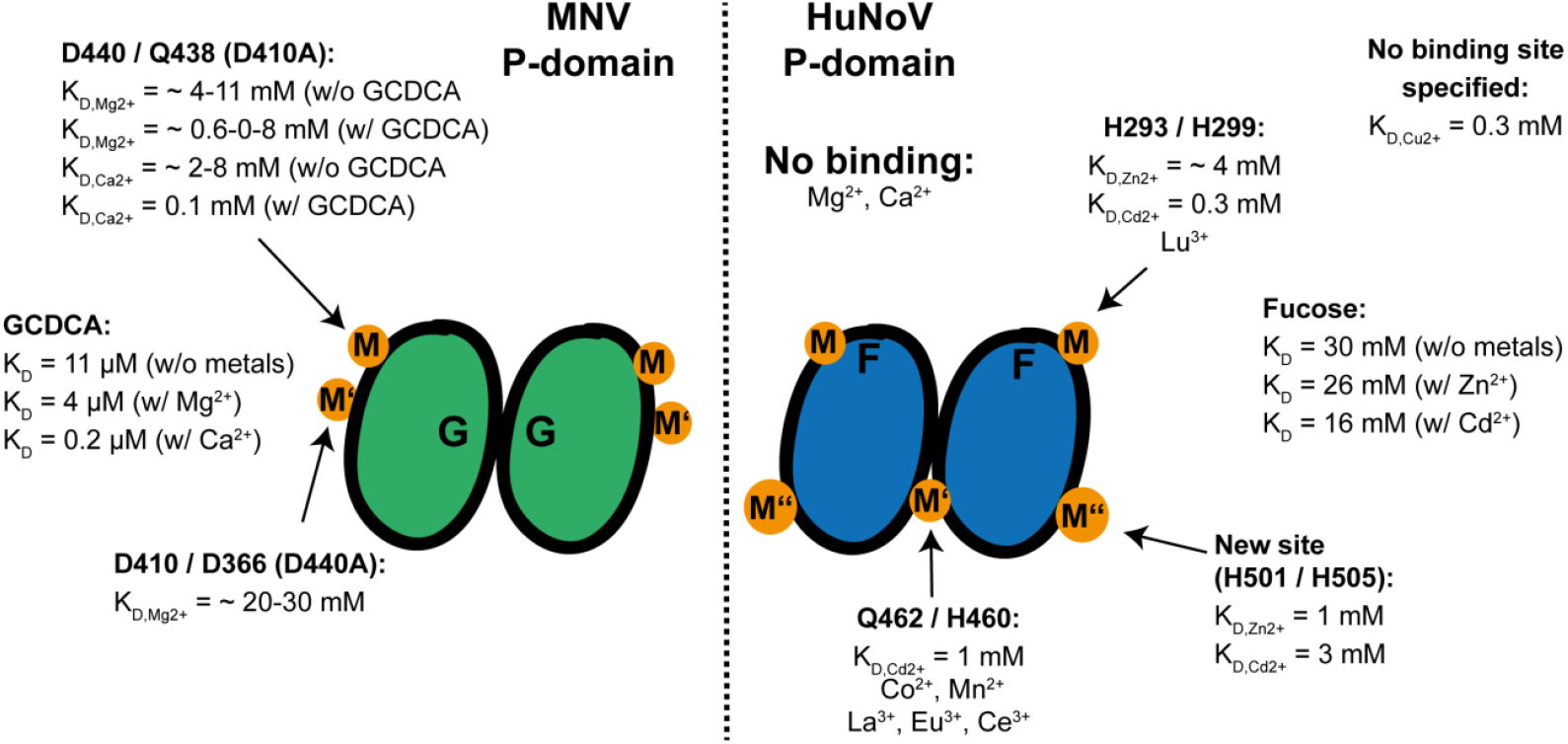
Binding of bivalent metal ions to MNV and HuNoV P-domains. Binding of Mg^2+^ or Ca^2+^ and GCDCA to MNV P-domains independently stabilize P-dimers. In addition, binding of GCDCA is synergistically affected by Mg^2+^ or Ca^2+^, and *vice versa*. HuNoV GII.4 Saga P-domains form stable P-dimers in solution, independent of the presence of bivalent metal ions or GCDCA. Neither Mg^2+^ nor Ca^2+^ bind to Saga P-dimers, but several other bivalent metal cations, some being biologically relevant trace metals. Zn^2+^ or Cd^2+^ enhance the binding affinity for L-fucose, which is the minimal recognition element of HBGAs.

The binding of bivalent metal ions to MNV P-dimers causes long-range CSPs. Interestingly, the long-range effects in the E’F’ loop are similar to the ones observed upon addition of GCDCA^[16]^. When GCDCA is added to P-domains saturated with bivalent metal ions, a similar pattern of long-range effects is observed, except for effects in the E’F’ loop. When divalent metal ions are added to P-domains saturated with GCDCA, no long-range CSPs are observed at all. This indicates that the conformational transition towards the closed conformation as observed by cryo-EM is complete with either of the two co-factors alone. It is important to note that the PCS NMR experiments have shown that the structures of the GCDCA and metal ion bound forms of the P-domain are almost identical. Therefore, we attribute the long-range CSPs and other long-range effects such as line broadening to subtle conformational changes of amino acid side chains. In this respect it is interesting to note that our measurements of methyl order parameters S^2^ indicate an increase of methyl side chain mobility upon addition of GCDCA. This may be interpreted as a “melting” of the respective side chains and likely of other amino acid side chains upon binding of GCDCA. This observation leads to the conclusion that there is a favorable entropic contribution to co-factor binding stemming from internal side chain motions as this has been discussed already for other cases^[49-51]^. Taken together, for MNV P-domains we have underpinned some of the changes of affinities with kinetic data from 2D-NMR line shape analysis. The binding kinetics of bivalent metal ions as well as of GCDCA is substantially altered when the other co-factor is already present in saturating amounts, as this is obvious from the transition from slow to fast exchange of metal ions upon addition of GCDCA (cf. Figure 3). This may be interpreted as conformational pre-organization of, e.g., the GCDCA binding site by binding of Mg^2+^ or Ca^2+^.

Binding of bivalent metal ions to P-dimers of two predominant HuNoV genotypes, GII.4 Saga and GII.17 Kawasaki, is in stark contrast to this complex scenario where dimerization of MNV P-domain monomers and binding of GCDCA is synergistically coupled with bivalent metal ion binding. The most striking observation is that neither Mg^2+^ nor Ca^2+^ caused CSPs or changes in line intensities of methyl TROSY spectra of MILVA-labeled HuNoV P-dimers (Figure S20a and b), ruling out binding under these conditions. However, we observed CSPs upon addition of Zn^2+^ or Cd^2+^ (Figures 6 and S22), with dissociation constants in the low mM range. Along with the CSPs, overall signal intensities decreased. EDTA partially restored the original spectra (Figures 6a, 6b, S22a, and S22b) but in the case of Zn^2+^, additional peaks not observed in the apo-form remained. We suggest that binding of Zn^2+^ or Cd^2+^ caused partial unfolding and aggregation. The CSPs, changes in signal intensities, and the increase of transverse relaxation rates R2 were mapped to two different regions *1* and *2* in the case of Zn^2+^ and Cd^2+^ (Figures 6c-f), and to a third region *3* in the case of Cd^2+^ only (Figures S22c-e). The binding regions *1* and *3* comprise two known binding sites for bivalent metal ions. For region *2* no metal ion binding site has been reported so far. The potential binding sites will be discussed in the following.

Region *1* comprises a binding site for Zn^2+^ that has been observed in a cryo-EM structure of GII.2 snow mountain virus (SMV) VLPs^[18]^, with the P-domains residing in the so-called resting conformation. The chelating amino acids were His293 and His299, located in a surface loop adjacent to the loop that holds the fucose residue of HBGAs. The superposition of the crystal structure of GII.4 Saga P-domain (PDB 4X06) with the cryo-EM structure (PDB 6OUC) suggests that in the GII.4 Saga P-dimers, His292 and Tyr299 may substitute for His293 and His299. This matches well with the observed spectral perturbations in this region *1* of the P-dimers (Figures 3f and S6e). We conclude that the binding site in region *1* corresponds to the site observed in the cryo-EM structure^[18]^. The authors of the cryo-EM study have hypothesized that binding of Zn^2+^ allosterically enhances binding of HBGAs by stabilizing a conformation of Asp382 that is required for binding to fucose. Therefore, we tested whether the addition of Zn^2+^ or Cd^2+^ would affect HBGA binding. Indeed, the presence of Zn^2+^ or Cd^2+^ slightly improved the affinity of Saga P-dimers for methyl-α-L-fucopyranoside (Figure S23), supporting the hypothesis.

For binding region *2* (Figures 3f and S6e) that was identified for both, Zn^2+^ and Cd^2+^, we could not find a known binding site. The perturbations map to amino acids (Val499, Leu507, Val508, Ile509, and M530) that constitute the GCDCA binding site^[8]^. Within this region in GII.4 Saga P-dimers two histidine residues, H501 and H505, could form a binding site for bivalent metal ions (Figure 6f). These two amino acids are highly conserved in GII.4 genotypes.

Within binding region *3* a binding site for bivalent metal ions has been identified from high-resolution crystal and cryo-EM structures of VLPs of the GII.4 Houston 2002 strain^[11]^. In these structures the P-domains as parts of the VP1 capsid protein are also in the so-called resting conformation. The bivalent metal ion, in this case Cd^2+^, is located at the interface between the monomeric P-domains and is held by symmetrically arranged His and Glu residues (His460, His460’, Glu463, and Glu463’)^[11]^. Zn^2+^ did not cause any spectral perturbations in the vicinity of this site (Figure 3) but in the case of Cd^2+^ we observed significant CSPs (I231) or extensive line broadening (V232) in this region *3* (cf. Figures S6c and S6d). Interestingly, the paramagnetic metal ions studied here (Figure S24) almost all cause perturbations within region *3*, except for Cu^2+^ that binds in the vicinity of the HBGA binding site (Figure S25).

It may be hypothesized that the essential trace element ions Zn^2+^, Mn^2+^, and Cu^2+^ play a biological role in infection with HuNoV as these ions bind to P-dimers and in the case of Zn^2+^ binding to Saga P-dimers we even detected a synergistic influence on the binding of L-fucose. From our experiments we cannot exclude that under *in vivo* conditions Mg^2+^ and Ca^2+^ also play a role since we have been studying isolated P-dimers in aqueous solution instead of the complete VP1 as part of an intact virion. For instance, the two P-domains forming the binding site within region *3* may be in a slightly different orientation in complete virions, allowing binding of Mg^2+^ and Ca^2+^. Taken together, the metal ion binding sites identified by cryo-EM and crystallography in intact virions and P-dimers have been quantitatively characterized by NMR experiments, and the comparison of crystallographic and cryo-EM studies with NMR data suggests that binding of bivalent metal ions is modulated by arranging VP1 domains in intact virions. Our observations highlight the conformational flexibility inherent to HuNoV P-domains. To fully understand the role of bivalent metal ions in the transition between resting and raised conformations in HuNoVs more studies are needed.

## Conclusion

For MNV P-dimers it is shown how binding of Mg^2+^ and Ca^2+^ synergistically interacts with the binding of GCDCA. Binding of either partner pre-organizes conformations that facilitate binding of the other ligand (Figure 8). Beyond this synergistic behavior, both types of ligands induce long-range CSPs suggesting allosteric effects as evidenced by other studies that describe large scale structural changes of MNV capsids and differences in binding of neutralizing antibodies upon addition of GCDCA and/or Mg^2+^ or Ca^2+^. Here, we show that this synergistic behavior can also be observed in virus neutralization assays, thus providing a molecular explanation for the influence of bivalent metal ions in MNV neutralization.

The comparison with metal ion binding to HuNoV P-dimers reveals substantial differences (Figure 8). Under the conditions chosen (isolated P-dimers instead of complete virions) neither Mg^2+^ nor Ca^2+^ bind. The synergism between P-domain dimerization, binding of bivalent metal ions, and binding of GCDCA is absent in the HuNoV P-dimers studied here. Bivalent metal ions may nevertheless play a role in stabilizing the “resting” conformation of virions and could serve as co-factors in infection, but our results strongly suggest that the mechanisms behind this would be different from the ones operative in MNV infection. These findings call for systematic investigations into the role of metal ions in HuNoV infection.

### Experimental Section

#### Protein biosynthesis and purification

The synthesis of MNV P-domains^[21]^ and of HuNoV P-domains^[38]^ has been performed according to the published procedures^[21, 38]^. Briefly, E. coli was transformed with a pMAL-c2X expression plasmid^[52]^ containing genes for ampicillin resistance and a fusion protein consisting of maltose-binding protein (MBP) and the GV CW1 murine norovirus VP1 P-domain (referred to as MNV P-domain, GenBank ID DQ285629, amino acids 228-530, N-terminal addition of GP peptide), or the non-deamidating N373D GII.4 Saga4/2006 P-domain (referred to as HuNoV P-domain, GenBank ID AB447457, amino acids 225-530, N-terminal addition of GPGS peptide). The resulting MBP and P-domain proteins are separated by two His8-tags and a HRV 3C protease cleavage site (LEVLFQGP) for protein purification. Plasmids for expression and protein biosynthesis of MNV P-domain point mutants were generated by site-directed mutagenesis using primers listed in Table S4.

Biosynthesis and protein purification of [U-^2^H,^15^N] and [^1^H,^13^C] MILVA methyl group labeled HuNoV P-domains, as well as the production of stable, non-deamidating N3l3D HuNoV P-domains have been explained in detail previously^[21, 38]^. Unlabeled MNV P-domains were expressed in terrific broth medium. Protein expression was induced using 0.1 mM IPTG at an OD^600^ of 1.3. Cells were harvested after 24 h at 16 °C and the pellets were stored at -70 °C until further processing. [^1^H, ^13^C] MILVA- and [^1^H,^13^C] MI*LVA-methyl group labeled proteins (Table S5) were expressed in D_2_O based minimal medium containing 3 g/L of D-glucose-d_7_ (Deutero) and 3 g/L ^14^NH_4_Cl, respectively. *E. coli* BL21 (DE3) cells carrying the pMAL-c2X plasmid encoding wild type MNV P-domain, point mutant MNV P-domain or N373D HuNoV P-domain proteins were used to inoculate 40 ml LB medium with 100 μg/ml ampicillin. Cells were grown over night at 37 °C. On the next day, 40 ml D_2_O based minimal medium with 100 μg/ml ampicillin was inoculated with a cell pellet obtained from the overnight culture resulting in an OD_600_ of 0.05. Cells were grown at 37 °C until an OD_600_ of 0.5. Cells were centrifuged at room temperature and resuspended in a volume of D_2_O based minimal medium with 100 μg/ml ampicillin corresponding to 4/5 of the final culture volume. The cells were grown at 37 °C until an OD_600_ of 0.75. A volume of D_2_O based minimal medium with 100 μg/ml ampicillin corresponding to 1/5 of the final culture volume was used to solve precursors listed in Table S5. The pH was adjusted to 7 and the solution was sterile filtrated. The temperature of the culture was decreased to 16 °C and the solution containing the precursors were added to the culture. Protein biosynthesis was induced after 1 h by addition of 0.1 mM IPTG. Cells were grown until the stationary phase was reached, harvested, and stored at -70 °C until further processing.

Protein biosynthesis of [U-^2^H,^15^N] and [^1^H,^13^C] MILVA methyl group labeled MNV P-domain proteins for measurement of molecular correlation times (see Supplementary methods) was carried out as explained above. The only difference was the substitution of ^14^NH_4_Cl with ^15^NH_4_Cl (Deutero) in D_2_O based minimal medium preparation.

Cell pellets for purification of MNV P-domain were resuspended in chilled buffer containing 20 mM sodium acetate and 100 mM NaCl at pH 5.3 for MNV P-domains or PBS pH 7.3 for HuNoV P-domains. 10 mg/ L chicken egg white lysozyme (Merck Millipore), 160 U/L benzoase (Novagen), 0.48 mg/L leupeptin (Sigma) and 0.48 mg/L aprotinin (Roth) were added, and cells were subjected to two lysis cycles using a high-pressure homogenizer (Thermo) applying a pressure of 1400 psi. Cell lysate purification was centrifuged for 90 min with 7000 xg at 4 °C, and the supernatant was than subjected to further purification steps.

Protein purification was carried out on an ÄKTA pure system (GE Healthcare) at 4 °C. The MBP P-domain fusion protein was separated from the *E. coli* lysate employing two coupled 5 ml Ni-NTA columns (GE Healthcare). For MNV P-domains, the columns were equilibrated using 20 mM sodium acetate, 100 mM NaCl (pH 5.3). The lysate was loaded, and the columns were washed with equilibration buffer. A stepwise elution supplementing the equilibration buffer with 10 % 20 mM NaPO_4_, 100 mM NaCl, 500 mM Imidazole (pH 7.4) elution buffer was used to clear the columns from remaining *E. coli* proteins. Supplementation with 100 % elution buffer yielded the MNV fusion protein.

Fractions containing the fusion proteins were pooled and filled up with 20 mM sodium acetate, 100 mM NaCL (pH 5.3) to a final volume of 40 ml. 200 μl of 4 mg/ml His-tagged HRV 3C protease was added to cleave the fusion protein. The solution was dialyzed against 20 mM sodium acetate, 100 mM NaCl, 5 mM β-mercaptoethanol (pH 5.3) over night at 4 °C. On the next day, a further purification step employing Ni-NTA chromatography was performed. The columns were equilibrated as described before. The solution containing the His-tagged protease, the His-tagged MBP, and the MNV P-domain were loaded on the columns, with the P-domain not able to attach to the resin anymore. P-domain containing fractions were pooled and concentrated using 10 kDa centrifugal filters (Merck) to a final volume of 4 ml. Remaining proteins were eluted from the columns using stepwise gradients of 10 % and 100 % elution buffer as explained before. For a last purification step, concentrated MNV P-domain proteins were subjected to size exclusion chromatography (SEC) employing a preparative Superdex 16/600 200 pg size exclusion column (GE Life Science). 20 mM sodium acetate, 100 mM NaCl (pH 5.3) was used as equilibration and running buffer. The P-domain eluted from the column at an elution volume of ca. 80 to 90 ml.

### NMR sample preparation

Sample pH values of H_2_O-based buffers were determined using an Orion 8220BNWP pH-electrode (Thermo Fisher) with an Orion Star A221 pH-meter (Thermo Fisher). The electrode was calibrated with H_2_O-based pH 4, 7 and 10 standard solutions (Roth) prior to measurements. For D_2_O-based buffers, the obtained pH values (pH*) were converted into pH_corr_ values using Eq. 3 ^[39]^:

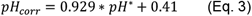

Note that in the literature, mostly pH* values are given for D_2_O-based solutions. Using pH_corr_ values instead of pH* values serve the comparability to other studies carried out in H_2_O. For solutions constituting out of a H_2_O/D_2_O mixture, the pH* is given.

MNV and HuNoV P-domain proteins were concentrated using 10 kDa centrifugal filters (Merck Millipore). Buffer exchange into the proteins’ respective analysis buffers (see below) was performed using ZebraTM Spin Desalting columns (MWCO 40 kDa, Thermo Scientific) after the manufacturer’s instructions or 10 kDa centrifugal filters (Merck, Millipore). Concentrations were determined using UV absorption at 280 nm measured with an UV spectrometer (Peqlab) using a molecular extinction coefficient of 46.870 M^-1^cm^-1^ and a molecular weight of 33.2 kDa for unlabeled MNV P-domains, and 35.410 M^-1^cm^-1^ and a molecular weight of 36.6 kDa for MILVA HuNoV P-domains. Potential ligand molecules or metal ions were titrated to the protein samples from highly concentrated stocks in the same buffer with carefully adjusted pH, or pH_corr_, or pH* values. The ^1^H NMR signals of imidazole were used to control the pH in the samples during NMR experiments.

### Analytical size exclusion chromatography (SEC) with MNV P-domains

Analytical SEC was carried out on an ÄKTA pure system (GE Healthcare). Protein solutions containing the P-domain in 20 mM sodium acetate, 100 mM NaCl (pH 5.3) with P-domain concentrations listed in Table S6 were subjected to a Superdex 75 Increase 3.2/300 column (GE Life Sciences) pre-equilibrated in the respective buffers using a 10 μL sample loop by filling the entire loop with the protein solution. UV absorption was measured at 280 nm. The SEC run was performed at 4 °C employing a flow rate of 0.075 ml/min using the respective buffer listed in Table S6.

### NMR spectroscopy

If not stated otherwise, all spectra were acquired on a Bruker Avance III HD spectrometer equipped with a TCI cryogenic probe. All spectra were processed and analysed using *TopSpin* v3.6 (Bruker) if not stated otherwise. Peak intensities and chemical shifts were extracted using CcpNmr v2.4.2 ^[53]^

In the case of [^1^H,^13^C] MILVA or [^1^H,^13^C] MI*LVA methyl group labeled MNV P-domains, methyl TROSY experiments with were carried out using 512 increments, 0.8 ppm as center of the spectral windows, and a sweep width of 3.5 ppm in the direct dimension. The respective parameters for the indirect dimension were 256 data points, 17 ppm and 18 ppm. For [U-^2^H,^15^N], [^1^H,^13^C] MILVA methyl group labeled N373D HuNoV P-dimers, methyl TROSY experiments with were carried out using 512 increments, 0.75 ppm as center of the spectral windows, and a sweep width of 3.75 ppm in the direct dimension. The respective parameters for the indirect dimension were 512 data points, 17.5 ppm and 18 ppm. The number of scans was either 4 or 8, and the relaxation delay was set to 1.5 s in all experiments. Further parameters and protein concentrations are specified in the respective figure legends. If not stated otherwise, the following buffers were used: i) for MNV P-domains 20 mM sodium acetate-d_4_ (Cambridge Isotope Laboratories), 100 mM NaCl (pH_corr_ 5.3). ii) For HuNoV P-domains 50 mM bis-tris-d_19_ (Cambridge Isotope Laboratories), 150 mM NaCl (pH_corr_ 6.80) (c.f. sample preparation section).

### Measurement of CSPs

CSPs in Hz Δv_eucl_ between resonances in [^1^H, ^13^C] HMQC spectra at different conditions are calculated using:

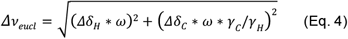

with Δδ_H_ and Δδ_C_ being the chemical shift differences in the respective dimensions in ppm. γ_H_ and γ_C_ are the gyromagnetic ratios of the respective nuclei. ω is the spectrometer’s frequency for ^1^H nuclei (i.e. 600 MHz for all spectra shown here).

### Measurement of PCSs

^1^H PCSs δ^PCS^ in ppm between certain resonances in [^1^H,^13^C] HMQC spectra of diamagnetic and paramagnetic samples were calculated using:

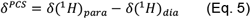

with δ(^1^H)_para_ and δ(^1^H)_dia_ being the ^1^H chemical shifts under paramagnetic and diamagnetic conditions in ppm.

Magnetic susceptibility tensor parameters and paramagnetic centers of anisotropic samples were fitted using the *Paramagpy* software package^[40]^ according to equations 6 to 8.

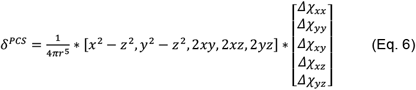

r is the distance between the paramagnetic center and the nucleus of interest. x, y, and z are the cartesian coordinates of the paramagnetic center in the protein frame. Δχ_kk_ with k ϵ [x,y,z] characterizes the anisotropic part of the magnetic susceptibility tensor Δχ.

Coordinates of the respective nuclei for calculation of r were obtained from crystal structure models, as described in ^[21]^. x, y, z, and Δχ_xx_, Δχ_yy_, Δχ_xy_, Δχ_xz_, and Δχ_yz_ were fitted using experimentally derived ^1^H PCSs according to Eq. 6. Uncertainties in x, y, and z were obtained using 1000 bootstrap iterations sampling 90 % of the data. In a second fitting round, x, y, and z were held constant on the values derived before and the fitting was repeated to derive values for Δχ_xx_, Δχ_yy_, Δχ_xy_, Δχ_xz_, and Δχ_yz_. The axial and rhombic components of the magnetic susceptibility tensor can be calculated from the obtained parameters using Eqs. 7 and 8.

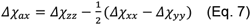

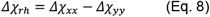

The orientation of Δχ’s frame to the protein frame is characterized by the Euler angles α, β, and γ. Uncertainties in Δχ_αx_, Δχ_rh_, α, β, and γ were determined using 1000 bootstrap iterations sampling 90 % of the data with x, y, and z hold constant as described above.

Quality factors (Q-factors) were derived using *Paramagpy* employing calculated PCSs δ^PCS,calc^ based on the above mentioned best fit’s parameters and experimental PCSs δ^PCS,exp^ according to Eq. 9:

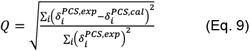

The index i is for summation over all spins.

Lines of best fit for correlation plots between experimental and calculated PCSs or experimental PCSs at different conditions were obtained using the linear regression tool implemented in *LibreOffice* v6.4.7.2.

### Estimation of off-rates and dissociation constants using 2D line shape analysis with TITAN

2D line shape analysis of [^1^H,^13^C] HMQC spectra was performed using the program TITAN ^[19]^. Spectra were processed using nmrPipe ^[54]^. Parameter uncertainties were obtained by 100 bootstrap iterations using TITAN’s error estimation tools.

Analysis of MNV P-domain dimerization at at pH_corr_ of 7.4 as shown in Figure 5 was carried out using *TITAN’s* built-in protein dimerization model and spectra of 13 μM, 25 μM, 50 μM, 75 μM, 100 μM, 150 μM, and 230 μM MNV P-domain. Chemical shifts and line widths of the monomeric P-domain (M) were fitted using the spectrum of the 13 μM P-domain sample. Chemical shifts and line widths of dimeric P-domain (D) were fitted using the spectrum of the 230 μM P-domain sample. In a last step, chemical shifts of the dimeric state, line widths, the off-rate k_off;Dimerization_, and the dissociation constant K_D;Dimerization_ were fitted using spectra of all P-domain concentrations with the line widths of all states and the chemical shift of the dimeric state derived above as starting conditions. The chemical shifts of the monomeric state were hold constant.

MNV P-domain metal ion binding analysis was performed employing a three-state binding model^[16]^. Spectra of D410A MNV P-domain in presence of 0 mM, 2.5 mM, 5 mM, 10 mM, 15 mM, 20 mM, and 25 mM MgCl_2_ or in presence of 0 mM, 5 mM, 10 mM, 15 mM, 20 mM, and 25 mM CaCl_2_ or spectra of MNV P-domain at a pH_corr_ of 7.4 in presence of 0 mM, 0.5 mM, 3 mM, 5 mM, 7.5 mM, 12.5 mM, and 20 mM CaCl_2_ were used. Chemical shifts and line widths of the monomer and dimer states were fitted using the spectra of the apo P-domain. Chemical shifts and line widths of the metal ion bound state were fitted using spectra with the maximal metal ion concentration. In the final run, all spectra were used to fit chemical shifts of the bound states with the values derived before as staring values. Additionally, line widths of all states, off-rates, and dissociation constants were fitted. Chemical shifts of the monomeric and dimeric states were kept constant at the values derived before. K_D;Dimerization_ and k_off;Dimerization_ were kept constant to the values obtained before for the corresponding pH_corr_ values (see ref.^[16]^ and Figure 5). All line widths and chemical shifts of the bound states obtained before were used as start conditions for the fit. A similar strategy was used for the analysis of Cd^2+^ binding to N373D HuNoV P-domain, although with a two-state binding model.

### Estimation dissociation constants from binding isotherms

Dissociation constants K_D_ were derived by fitting experimentally obtained NMR observables Δ_obs_ (CSPs or intensities) of n signals at certain ligand and protein concentrations L_t_ and P_t_ using the following system of equations^[30]^:

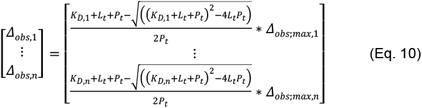

Δ_obs;max_ is the NMR observable at saturating ligand concentrations and is fitted for each signal individually. K_D_ values were globally fitted by constraining K_D,1_=…=K_D,n_ Errors were estimated using the standard error based on the residuals of the fit. Fitting of equations was carried out using *python*’s *symfit*, and *numpy* libraries. The *matplotlib* library was used for plotting binding isotherms.

### Deriving apparent pK_a_ values from CSPs

CSPs in Hz Δv_eucl_ (see above) of methyl group resonances at different pH_corr_ values (c.f. Figure S17) were used to fit apparent ionization constants pK_a_ to an adopted version of the Henderson-Hasselbalch equation^[55]^:

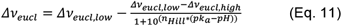

with Δv_eucl,low_ and Δv_eucl,high_ being the low pH and high pH CSP plateaus. n_Hill_ is the apparent Hill coefficient. pk_a_ values are regarded as apparent because methyl groups are not directly protonated. *Python*’s *numpy* and *symfit* libraries were used for the calculations.

### Measurement of increase in relaxation rates induced by metal binding

The increase of transverse ^1^H_M_-Г_2_ relaxation rates induced by binding of Co^2+^, Cu^2+^, Mn^2+^, Eu^3+^, and Ce^3+^ and the increase of ^1^H_M_-Г_2_ relaxation rates induced by binding of Zn^2+^ to N373D MILVA HuNoV P-dimers were obtained using the ^1^H,^13^C-HMQC-based pulse scheme described elsewhere ^[56]^. Seven relaxation delays (t=1, 6, 12, 25, 36, 50 and 100 ms) were acquired in an interleaved manner, with 4 scans (ca. 1 h) and with 512×512 data points in the direct and indirect dimensions, respectively. Measurements were repeat twice, and decays in cross-peak intensity were fitted to the function f(t)=I_0_ exp(-t/*T*_*2*_) using Dynamics Center V.2.5.2 (Bruker). The fit parameter error estimation was calculated from a Monte Carlo simulation with a 95% confidence level and the systematic error was estimated from the worst case per peak scenario, as defined in the Dynamics Center. Relaxation rates were calculated as the differences in transverse relaxation rates (^1^H_M_-Г_2_) between a sample containing 60 μM [U-^2^H,^15^N], [^1^H,^13^C] MILVA methyl group labeled N373D HuNoV P-domains and a second sample containing 60 μM of the same protein and ZnCl_2_, CoCl_2_, CuCl_2_, MnCl_2_, EuCl_2_ or CeCl_2_. For metal concentrations see Table S2.

### Analysis of structural models

Structural models were graphically illustrated using *pymol* (The PyMOL Molecular Graphics System, Version 2.5.4 Schrödinger, LLC.). Alignment of structural models and calculation of corresponding RMSD values was carried out using the “align” command. Three dimensional rotations were performed using the “turn” command. Distances were determined using the “distance” command.

### Cell culture

Murine microglial cells (BV-2) were maintained in Dulbecco’s Modified Eagle Medium (Gibco) supplemented with 5% FCS (Capricorn), 1x L-glutamine (Biozym), 0.1 mM non-essential amino acids (Biozym), and 100 units/ml penicillin and streptomycin (DMEM-5) incubated at 37 °C with 5% CO2 and 95% humidity as described^[57]^. Hybridoma suspension cell lines producing A6.2 MAb were maintained in Iscove’s Modified Dulbecco’s Medium (IMDM, Life Technologies) supplemented with 10% FCS (Capricorn), 1x L-glutamine, and 100 units/ml penicillin and streptomycin (IMDM-10) in spinner flasks at 30 rpm at 37 °C with 5% CO2 and 95% humidity as described previously^[16]^.

### Virus production, purification and titration

MNV-1.CW1 (GV/MNV1/2002/USA)^[58]^ (Karst 2003) was cultivated in BV-2 cells as previously described^[16]^ (Creutznacher et al 2022). Recombinant MNV viruses were produced using an adapted RNA-based reverse-genetics system^[59]^. Briefly, site-directed mutagenesis was performed using an adapted QuickChange approach^[60]^ using following primer sets: D410A-for: 5’ GAA TAC AAC GcT GGG CTA CTG GTT CCC CTT GC 3’ and D410A-rev: 5’ CCC AgC GTT GTA TTC AGG GAT GGT GTC CTG AAA AC 3’ as well as D440A-for: 5’ CAG ATC GcC ACC GCT GAC GCC GCA GCA GAG GCG 3’ and D440A-rev: 5’ CAG CGG TGg CGA TCT GAC GCA TGT AGG TCC GGA ACC 3’. Mutations in the recombinant viruses were sequence-confirmed at passage 3 and all recombinant viruses (including recombinant wild type) were used at passage 4 post-transfection. To obtain a cation-free stocks, virions were precipitated from clear-centrifuged cell lysate with a 45 % saturated solution of ammonium sulfate (v/v) and centrifuged for 15 min at 10,000 g. The supernatant was discarded and the pellet containing the concentrated virus was resuspended in 5 ml PBS pH 7.4 and sterile filtered using a 0.2 μM syringe filter. Stocks were further purified by ultracentrifugation through a 30 % sucrose cushion at 175,000 g for 6 h at 4 °C. The supernatant was again discarded and the pellet containing the purified virus was resuspended in 5 ml PBS pH 7.4 overnight at 4 °C and sterile filtered. Viral titers were determined by TCID_50_ as previously described^[59]^ and stored at -80 °C.

### Enzyme-linked Immunosorbent Assay (ELISA)

An indirect ELISA was used to measure the combined effect of ions and bile acid on MAb antibody binding. All assays were performed on Immulon® 2HB microtiter 96-well plates (Thermo Fisher Scientific). Wells were coated in 0.1 ml phosphate buffered saline (PBS pH 7.4) with 1 × 10^6^ TCID_50_ at 4 °C overnight (at least 17 h). After two washes with ELISA-wash buffer pH 7.0 (25 mM NaH_2_PO_4_ + 500 mM NaCl + 0.05 % Tween20), wells were blocked with 0.2 ml ELISA-block buffer pH 7.0 (ELISA-wash buffer + 3 % BSA) at 4 °C overnight. After washing four times with ELISA-wash buffer 0.1 ml of the Primary-Antibody-Mix was added (50 mM Tris pH 7.0 with 7.5 ng IgG A6.2 MAb plus bile acid (GCDCA, Sigma-Adrich), MgCl_2_, CaCl_2_, NaCl at indicated concentrations and incubated for 1 hour at 37 °C. The 50 mM Tris pH 7.0 was favored over PBS buffers to prevent metal ions from precipitating in phosphate buffer. The Primary-Antibody-Mix was removed, and wells were rigorously washed 8x with ELISA-wash buffer pH 7.0. A secondary HRP-conjugated anti-mouse IgG antibody (1:2,000 dilution in ELISA-Wash Buffer) was added and incubated overnight. After 8 washes plates were developed using 0.1 ml ABTS (2,2’-Azinobis [3-ethylbenzothiazoline-6-sulfonic acid]-diammonium salt, Pierce) and quenched with 0.05 ml Stop-Solution (1 % SDS) after exactly 20 min. A Tecan M Plex plate reader was used measuring the absorbance at 410 nm for the substrate using 490 nm as a reference. Analysis was performed using the Prism software package (GraphPad Software).

### IgG A6.2 antibody

The IgG Antibody A6.2^[61]^ was produced from suspension cultures as described previously^[16]^. Hybridoma cells producing A6.2 antibodies were a kind gift from Christiane Wobus (University of Michigan, USA). Briefly, hybridoma cells^[62-63]^ were grown in suspension and cells were removed by centrifugation. Antibody was precipitated with a 45 % (v/v) saturated solution of ammonium sulfate and purified using a protein G column according to the manufacturer’s recommendations.

## Supporting Information

The authors have cited additional references within the Supporting Information.^[11, 16-17, 21, 41-42, 64][30, 31]^

## Acknowledgements

This research was funded by the Deutsche Forschungsgemeinschaft (DFG) via grants Pe494/12-2 (T.P.) and TA1093-2 (S.T.) within the research unit FOR2327 (ViroCarb).

T.P. thanks the State of Schleswig-Holstein for supplying the NMR infrastructure (European Funds for Regional Development, LPW-E/1.1.2/857).

## Entry for the Table of Contents

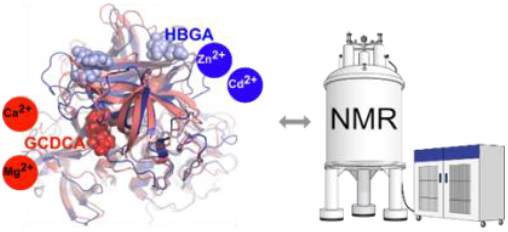

NMR experiments reveal distinct synergistic effects of the binding of bivalent metal ions to human and murine norovirus P-domains with other co-factors (glycochenodeoxycholic acid, histo-blood-group antigens). This has implications for structural models of human vs. murine norovirus infection.

## Notes

### Competing Interest Statement

The authors have declared no competing interest.

