## Supplemental figures, tables and methods for "NMR Reveals the Synergistic Roles of Bivalent Metal Ions in Norovirus Infections"

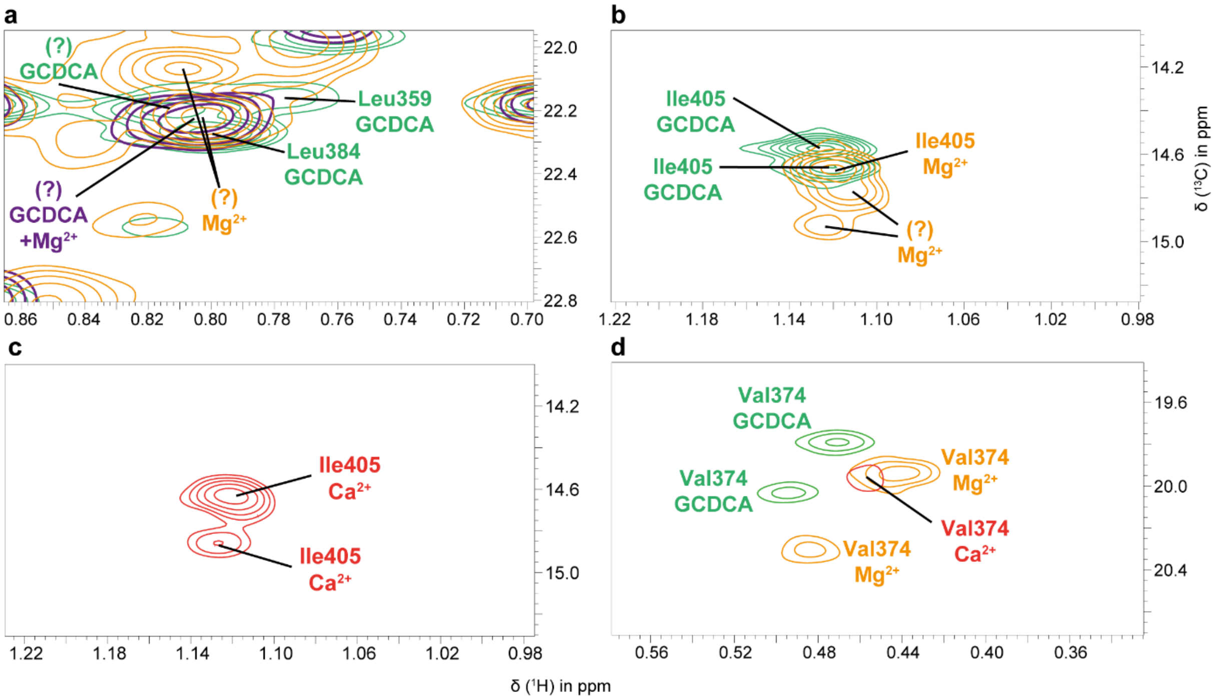


**Figure S1: Transfer of assignment of methyl resonance peaks is impossible for L359, L384, I405, and V374.**

The figure shows superpositions of sections of methyl TROSY spectra of $^{13}$C-methyl labeled MNV P-domain samples in the presence of GCDCA (green), with $Mg^{2+}$ bound (orange), with $Ca^{2+}$ bound (red), or with GCDCA and $Mg^{2+}$ bound (purple). a) Massive overlap of L384 and L359 cross peaks prevents transfer of assignment (exemplarily shown for $Mg^{2+}$). b) I405 is split into two resonances due to E/Z-isomerism of nearby P361[1]. In the presence of $Mg^{2+}$ a third peak of unknown origin is observed. c) For the $Ca^{2+}$ bound form two I405 peaks are observed as expected. d) In the presence of $Ca^{2+}$ the V374 methyl resonance is broadened beyond detection. Unassigned peaks are labeled with a question mark. Details of the $^{13}$C-methyl labeling schemes used for the preparation of the samples and conditions for the acquisition of methyl TROSY spectra are found in the legends to Figure 2 ($Mg^{2+}$ bound P-domain), Figure 4 (P-domain in the presence of GCDCA and $Mg^{2+}$), and Figure S11 (in the presence of $Ca^{2+}$ and in the presence of $Ca^{2+}$ and GCDCA). The methyl TROSY spectrum of MNV P-domain in the presence of GCDCA is taken from Creutznacher et al.[2] (see Figure 4 within that reference).

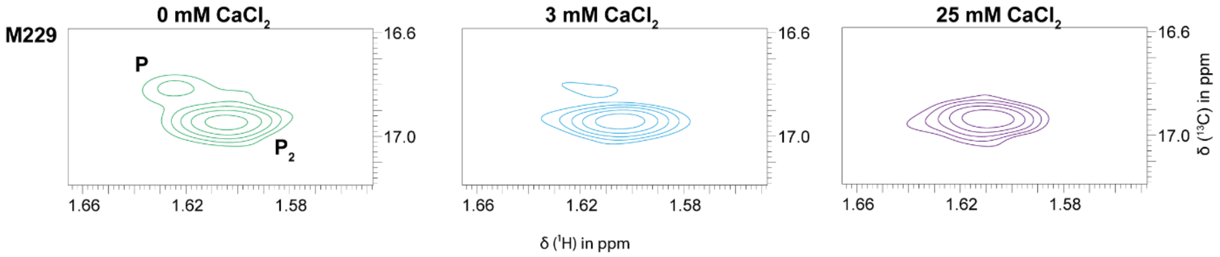


**Figure S2: $Ca^{2+}$ dependent depletion of monomer cross peaks in methyl TROSY spectra of the MNV P-domain.**

Addition of $CaCl_2$ to a MILVA labeled sample of the MNV P-domain leads to the disappearance of monomer resonances (M) whereas dimer resonances (D) remain. This is exemplarily shown for the M229 $^{13}C$ methyl resonance. The methyl TROSY spectra were acquired at 298 K and pH* 5.3. The samples contained a 39 μM solution of CW1 P-domain with $Ca^{2+}$ concentrations of 0, 3, and 25 mM.

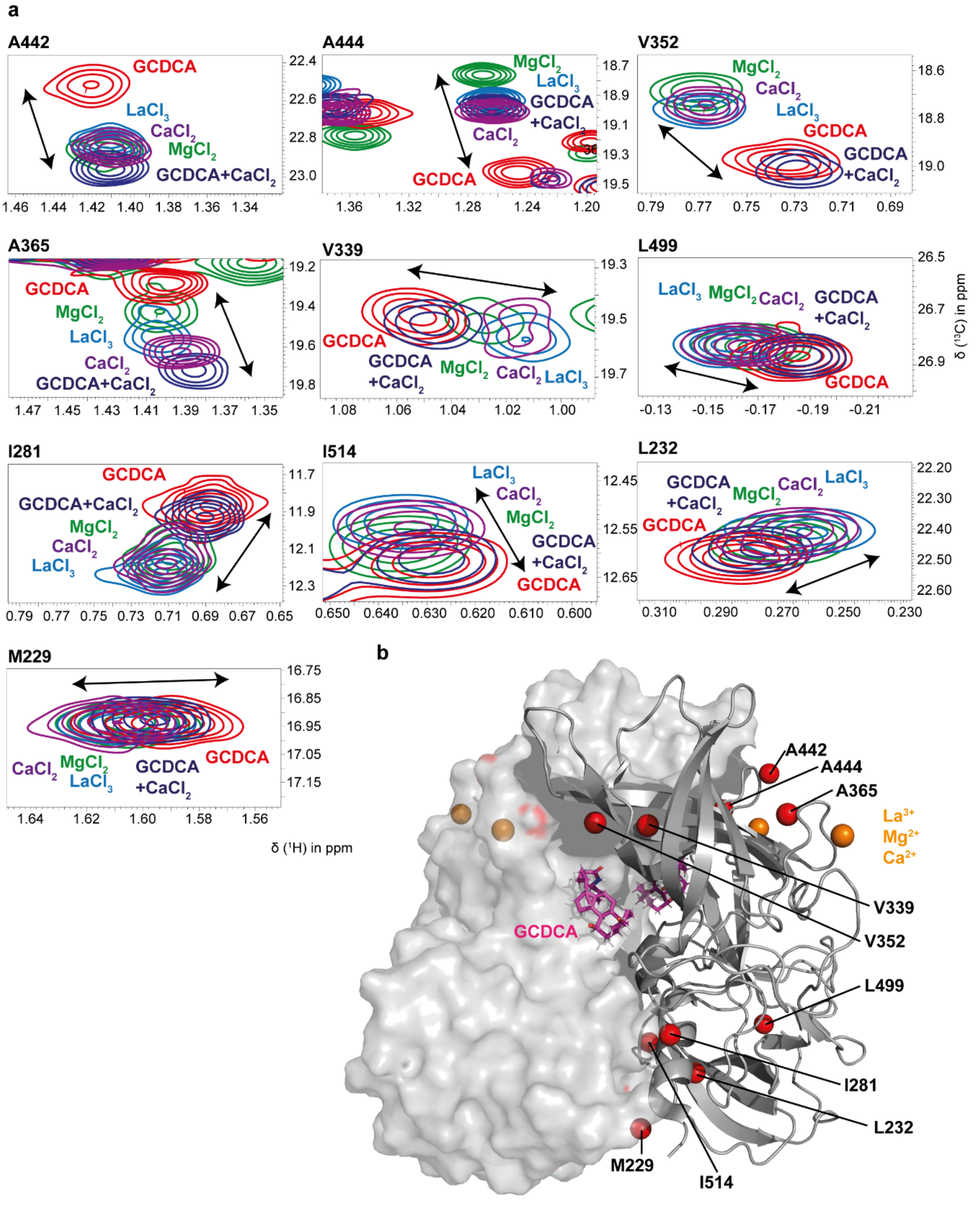


**Figure S3: CSPs caused by binding of $Ca^{2+}$, $Mg^{2+}$, $La^{3+}$, or GCDCA follow linear vectors.** a) Examples of long-range CSPs observed upon addition of $Ca^{2+}$ (25 mM), $Mg^{2+}$ (25 mM), $La^{3+}$ (0.4 mM), GCDCA (0.55 mM) or $Ca^{2+}$ and GCDCA (25 mM and 0.3 mM) follow linear vectors. Details of sample preparation and NMR data acquisition are found in the legends to Figures 2, S11, and S15 as well as in the legend to Figure 4 of Creutznacher et al.[2]. b) The methyl groups affected by ligand binding are distributed throughout the P-domain. The linearity of CSPs suggests the presence of an allosteric network, likely reflecting subtle global conformational and/or dynamic changes of amino acid side chains upon metal ion or GCDCA binding.

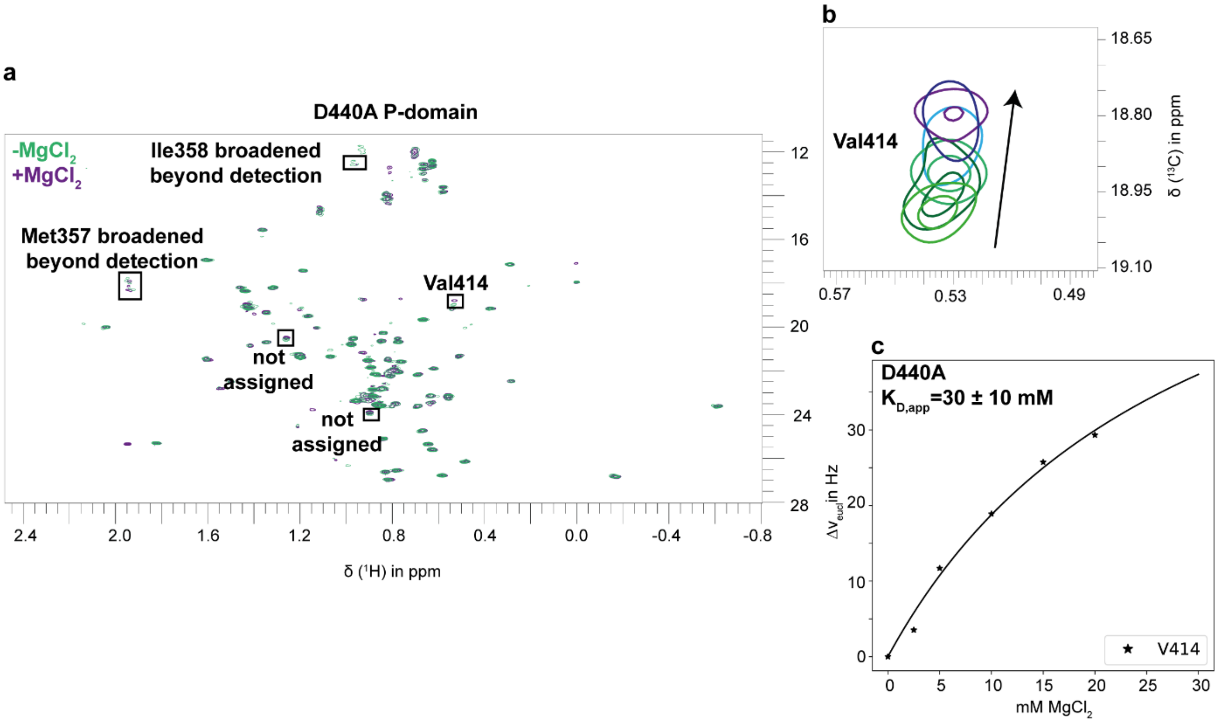


**Figure S4: $Mg^{2+}$ binding to the D440A P-domain.** a) Methyl TROSY spectra of the D440A P-domain in the presence and absence of $MgCl_2$ indicate $Mg^{2+}$ binding at the D410 metal ion binding site, which is part of the receptor binding epitope. Many peaks are broadened beyond detection, as indicated for Ile358 and Met357. b) The only assigned resonance showing CSPs is Val414, which is close to the D410 metal ion binding site. c) An apparent dissociation constant $K_{D,app}$ of 30 ± 10 mM was derived assuming a two-state binding model and using the CSPs measured for Val414. Spectra were acquired at 298 K on a 600 MHz NMR spectrometer with a cryogenic probe using a sample containing 27 µM MI*LVA labeled MNV CW1 P-domain (for details on MI*LVA labeling see the methods section).

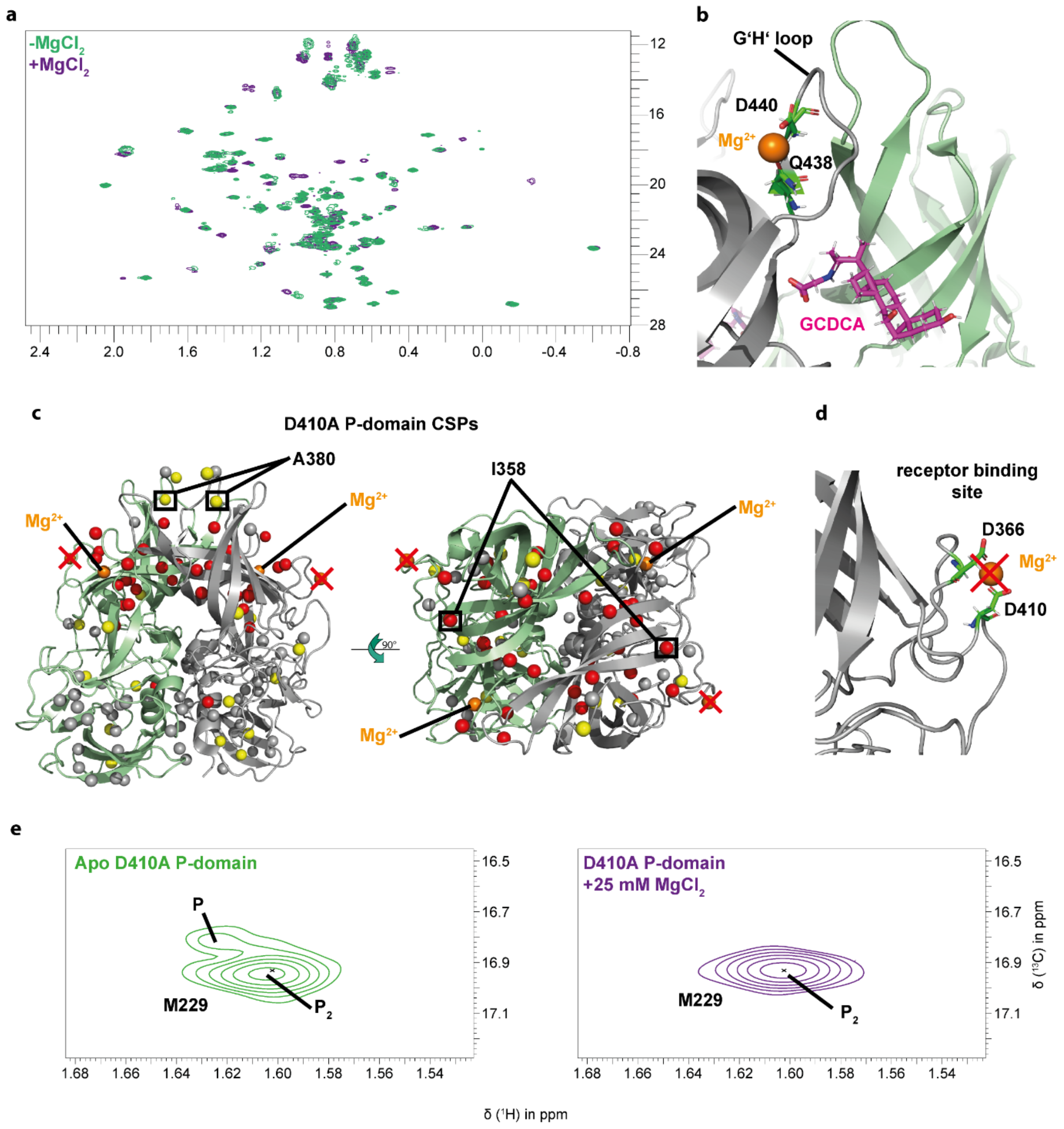


**Figure S5:** Methyl TROSY spectra of the D410A MNV P-domain in the presence and absence of 25 mM $Mg^{2+}$. Methyl TROSY CSPs observed upon adding $MgCl_2$ to a 62 µM solution of MI*LVA labeled D410A P-domain (a) at a temperature of 298 K and a pH* of 5.3. The color coding and significance thresholds are identical with the values given in the legend to Fig. 2. In (c) the CSPs are mapped onto the structure of the MNV P-dimer (pdb 6e47). Long-range CSPs affecting I358 (CD330lf binding site) and A380 (binding site for neutralizing antibodies) are highlighted. The active D440/Q438 metal ion binding site is shown in (b), and the deactivated D410/D366 site is shown in (d). Addition of $Mg^{2+}$ causes a stabilization of Pdimers ($P_2$) as shown for the M229 methyl TROSY cross peak (e).

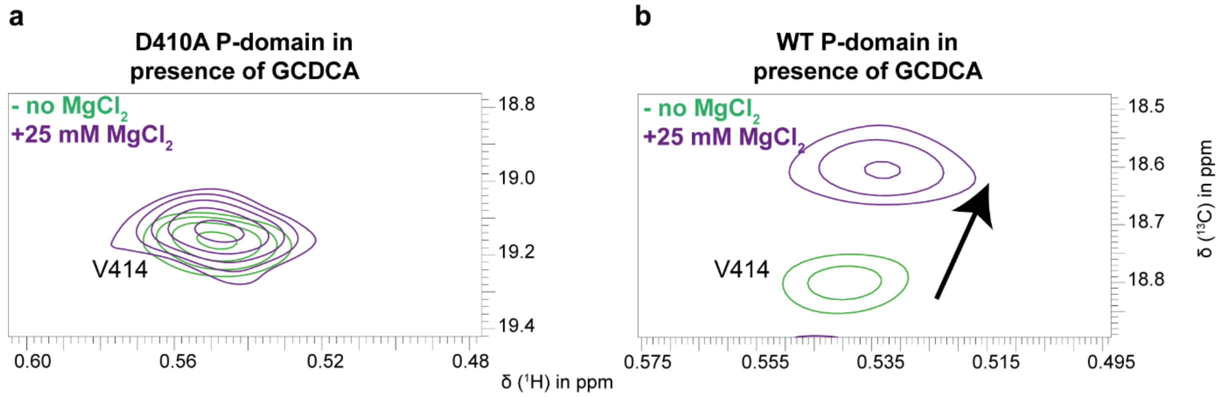


**Figure S6: The D410A mutant shows no binding of $Mg^{2+}$ to the binding site at the receptor epitope.** (a) shows the resonance corresponding to Val414 of the D410A P-domain in the presence and absence of $MgCl_2$. In comparison to the WT (wildtype) P-domain (b), the resonance of Val414 does not move significantly due to $MgCl_2$ addition indicating no significant binding to D410A P-domain at this binding site. Note that Val414 is close to Asp410, coordinating the metal ion (Fig. S5d). Spectra were acquired at 298 K on a 600 MHz spectrometer with cryo probe using 36 µM MI*LVA labeled MNV CW1 D410A P-domain or 75 µM MILVA labeled MNV CW1 WT P-domain in presence of saturating amounts of GCDCA (for details on MI*LVA labelling, see methods section).

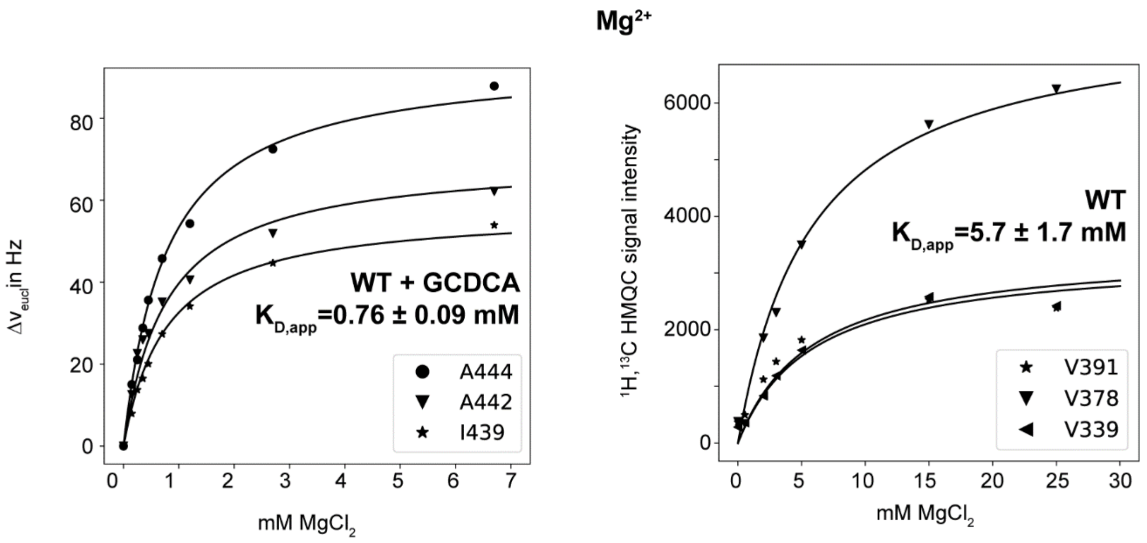


**Figure S7: Dissociation constants of $Mg^{2+}$ binding to the G'H' loop of the WT (wildtype) P-domain protein in the presence and absence of GCDCA.** The corresponding methyl TROSY spectra were acquired at 298 K on a 600 MHz spectrometer with cryo probe using 75 µM MILVA labeled MNV CW1 P-domain in absence of GCDCA (right panel) and 25 µM MILVA labeled MNV CW1 P-domain in presence of GCDCA (left panel).

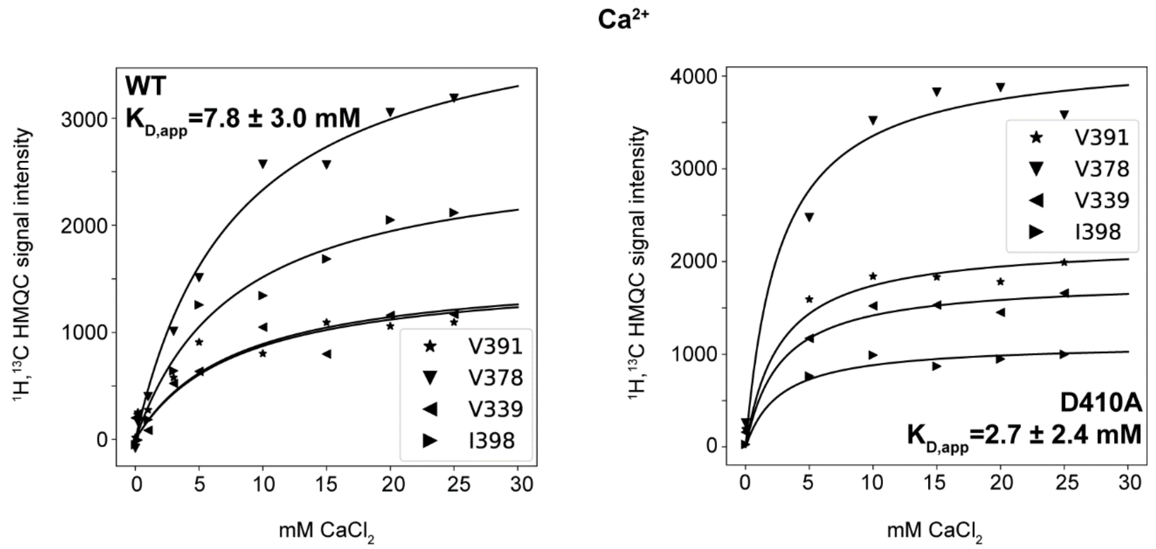


**Fig. S8: Dissociation constants of $Ca^{2+}$ binding to the G'H' loop of WT (wildtype) and D410A P-domain proteins in the absence of GCDCA.** The corresponding methyl TROSY spectra were acquired at 298 K on a 600 MHz spectrometer with cryo probe using 39 µM MILVA labeled MNV CW1 P-domain or 18 µM MI*LVA labeled D410A MNV CW1 P-domain (for details on MI*LVA labelling, see methods section).

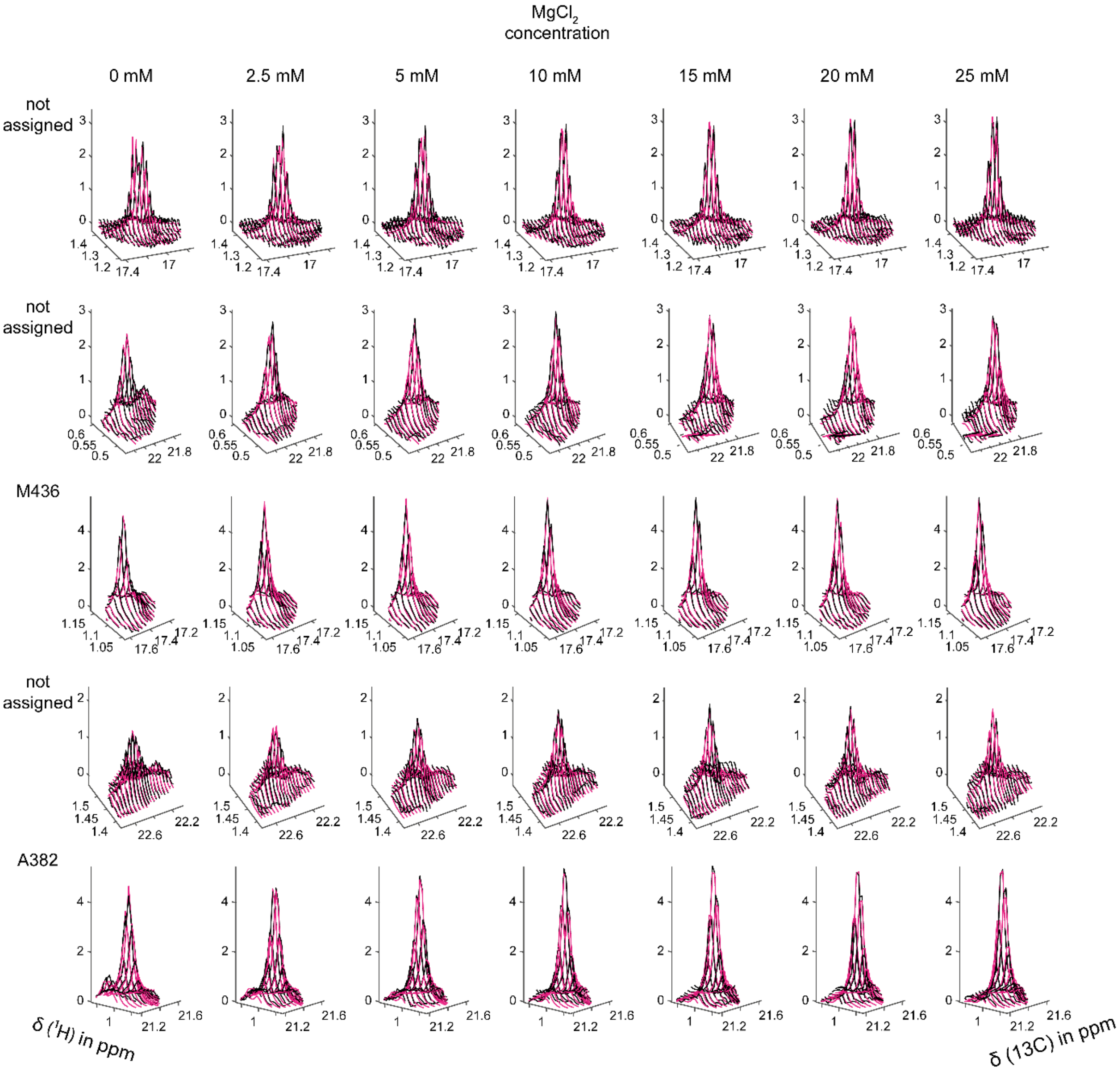


**Figure S9: Binding of $Mg^{2+}$ and $Ca^{2+}$ to the G'H'-loop of the D410A mutant P-domain in the absence of GCDCA - 2D line shape analyses of methyl TROSY cross peaks with TITAN**. Black and magenta spectra correspond to the measured and simulated spectra, respectively. For details of the acquisition of spectra, see Figs. S5 and S8 The overlay visually reflects the good quality of the fit. **$MgCl_2$:** For $Mg^{2+}$ the analysis yielded a $K_{D,Mg}$ of 3.8 ± 0.2 mM and a $k_{off,\,Mg}$ of 336 ± 692 $s^{-1}$. The large error for the determination of $k_{off,\,Mg}$ prevents this value from being used. **$CaCl_2$:** For $Ca^{2+}$ the analysis yielded a $K_{D,Ca}$ of 1.7 ± 0.2 mM and a $k_{off,Ca}$ of 3052 ± 2998 $s^{-1}$. Again, the large error for the determination of $k_{off,\,Ca}$ prevents this value from being used.

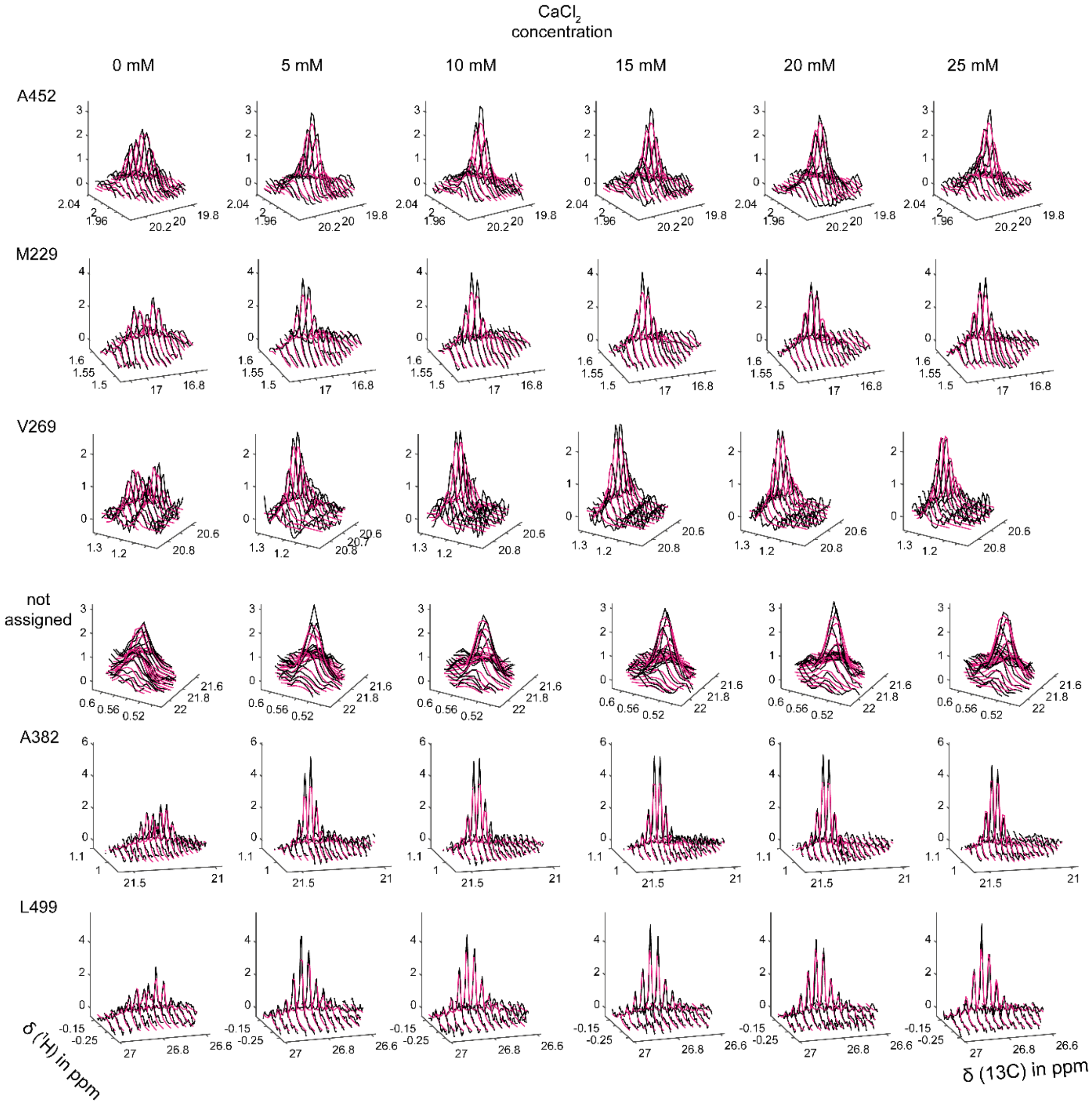


**Fig. S9 continued**

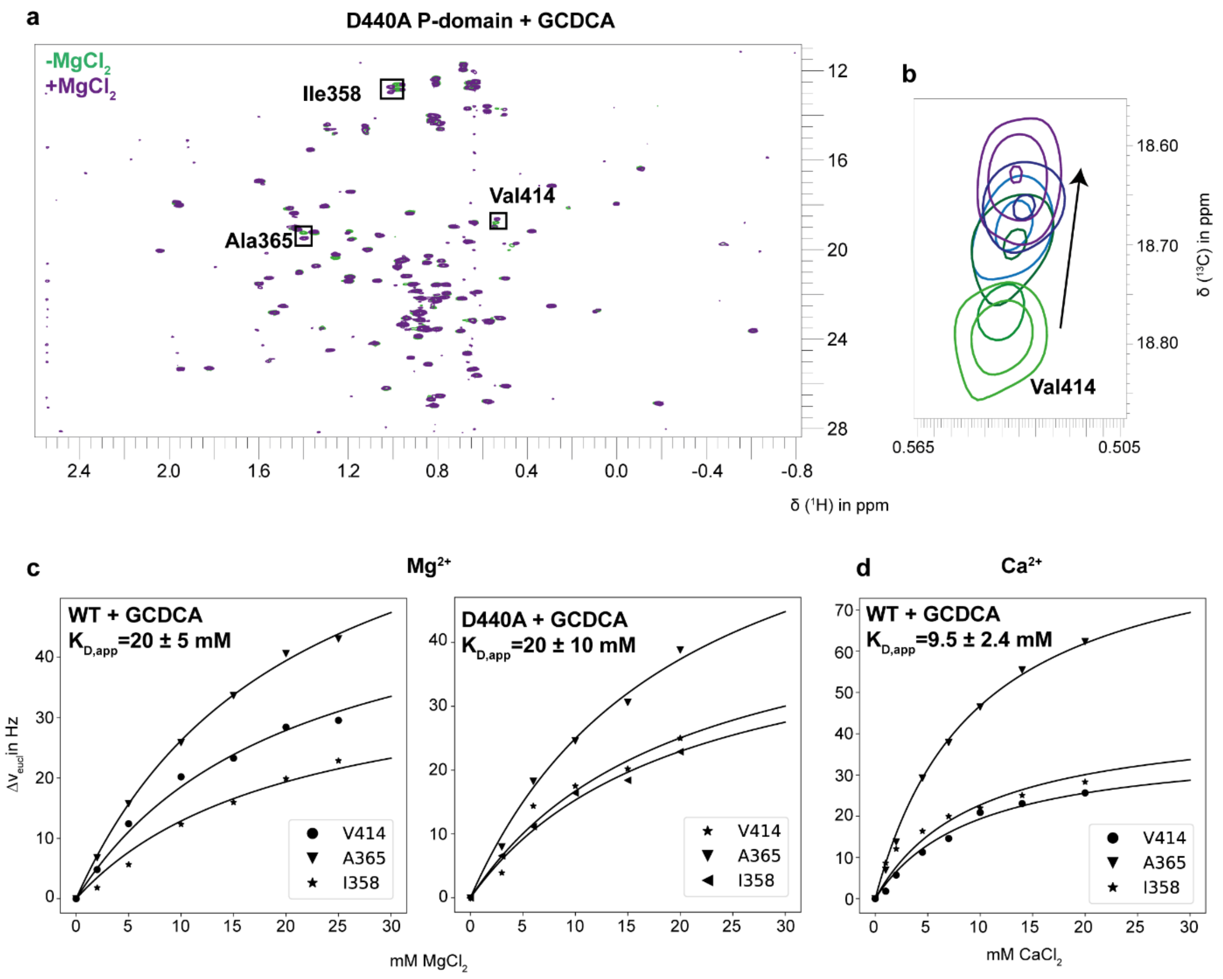


**Figure S10: Bivalent metal ion binding to the WT (wildtype) and D440A P-domain receptor binding site.** (a) Adding GCDCA to the D440A P-domain results in well resolved spectra. This contrasts with the spectra of the apo form (see Figure S4. Using chemical shift perturbations of suitable resonances (b) corresponding to methyl groups around the CD300lf receptor binding site yield dissociation constants $K_D$ for $Mg^{2+}$ around 20 mM in the presence of GCDCA (c) and a dissociation constant of ca. 10 mM for $Ca^{2+}$. The spectra were acquired at 298 K on a 600 MHz spectrometer with cryo probe using samples of 13 µM MI*LVA labeled D440A P-domain, 54 µM MILVA labeled MNV CW1 P-domain for the titration with $MgCl_2$ and 39 µM MILVA labeled MNV CW1 P-domain for the titration with $CaCl_2$ (for details on MI*LVA labelling, see methods section).

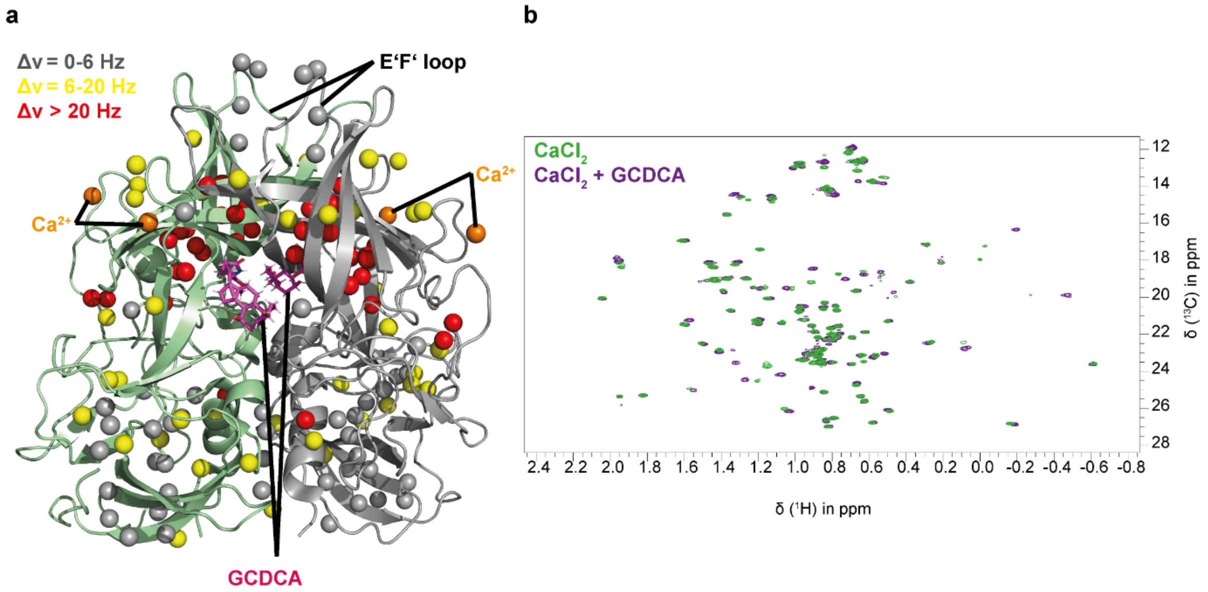


**Figure S11: Chemical shift perturbations (CSPs) upon adding GCDCA to MNV CW1 P-dimers in the presence of $Ca^{2+}$.** (a) For the illustration of CSPs a structural model of Nelson et al.[3] (pdb 6e47) was used. The positions of the $Mg^{2+}$ ions are labeled as "$Ca^{2+}$" for illustration purposes. (b) Methyl TROSY spectra of the P-domain in the presence of $CaCl_2$ with and without 300 µM GCDCA were acquired at 298 K on a 600 MHz spectrometer with cryo probe using 39 uM MILVA labelled MNV CW1 P-domain at 25 mM $CaCl_2$.

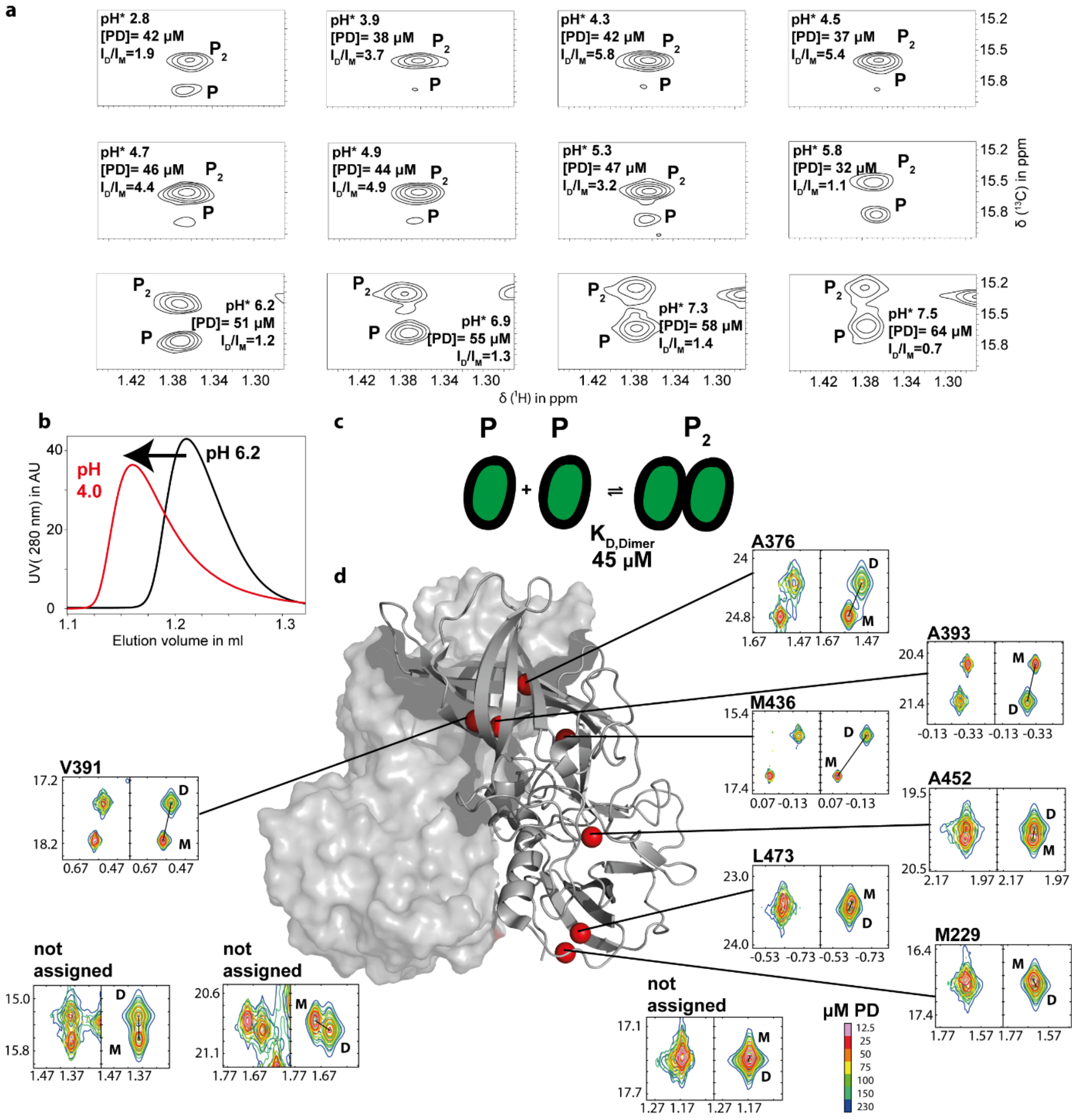


**Figure S12: The stability of MNV P-dimers depends on the pH.** (a) Relative intensities of cross peaks of monomer and of dimer resonances in methyl TROSY spectra as a function of pH (Samples contained 20 mM citric acid-d$_4$ , 100 mM NaCl for pH* values ranging from 2.8 to 5.8 (i.e. pH$_{corr}$ 2.9 to 5.8) and 20 mM Bis-Tris-d$_{19}$, 100 mM NaCl for pH* values ranging from 6.2 to 7.4 (i.e pH$_{corr}$ 6.2 to 7.4)). (b) Size exclusion chromatograms of MNV P-domain samples at two different pH values, reflecting a higher apparent molecular weight at pH 4.0. (c,d) 2D line shape analysis of cross peaks in methyl TROSY spectra of MNV Pdomain- at different protein concentrations and at pH* 7.5 (pH$_{corr}$ 7.4) yielded a dissociation constant $K_D$ of 45 ± 1.6 µM and a dissociation rate constant $k_{off}$ of 4.7 ± 0.3 s$^{-1}$. The left images show sections of the experimental spectra, and the right images show corresponding sections of simulated spectra. e) A structural model of the P-domain (pdb 6e47)[3] illustrates the position of the amino acid side chains used for line shape analysis.

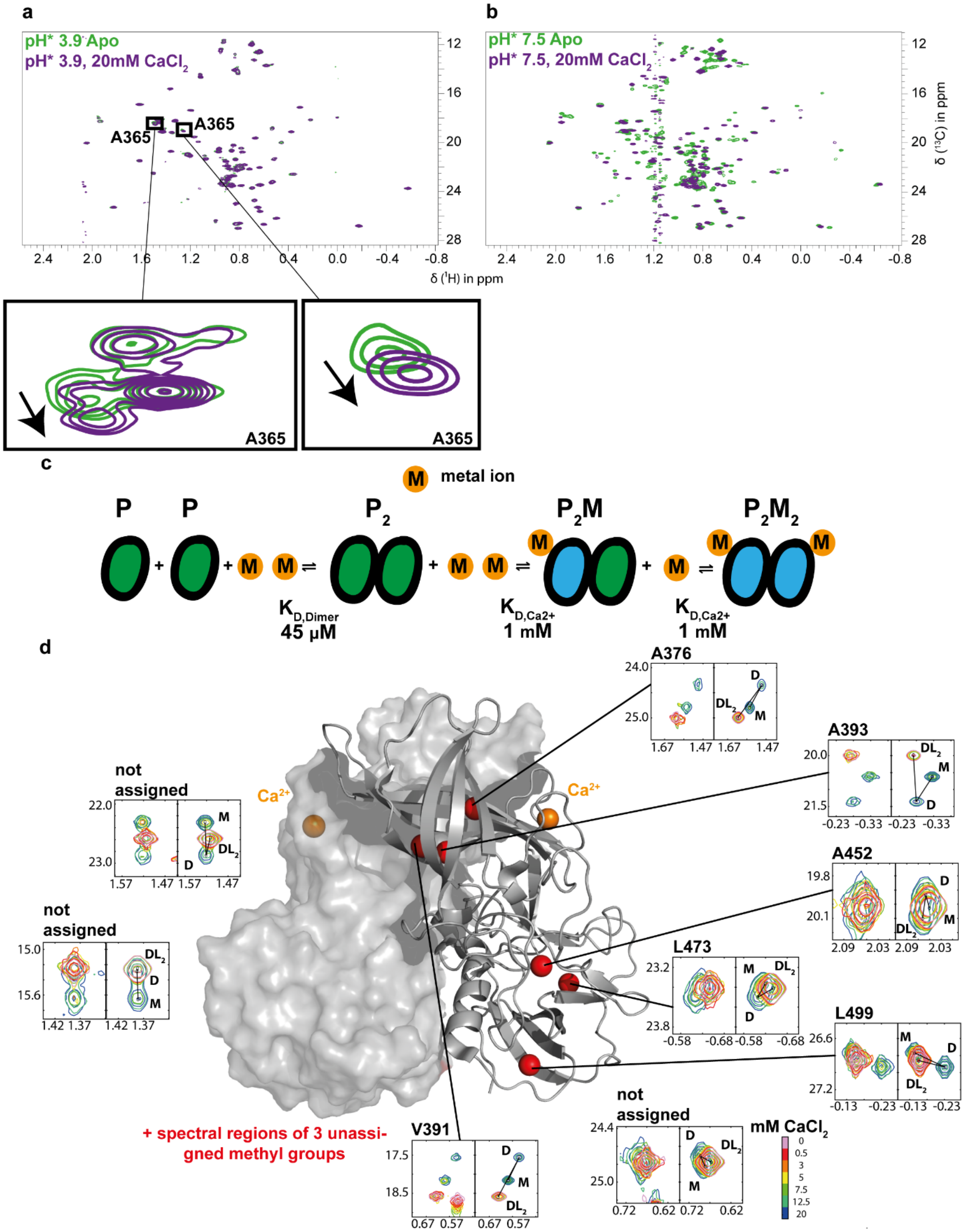


**Figure S13: Binding of $Ca^{2+}$ depends on the pH value. (a)** Methyl TROSY spectra of the MNV CW1 P-domain at a pH* of 3.9 ($pH_{corr}$ of 4.0) show no changes upon addition of 20 mM $CaCl_2$. Small CSPs observed for Ala365 reflect binding to the metal ion binding site that is close to the CD300lf receptor binding site. **(b)** At a pH* of 7.5 ($pH_{corr}$ of 7.4), titration with $Ca^{2+}$ allows observation of P-domain monomer peaks (P), of P-dimer peaks ($P_2$) and of peaks of P-dimer with $Ca^{2+}$ bound ($P_2M_2$). This allows application of a three-state model (c) for the simulation with TITAN. **(c, d)** Methyl TROSY spectra of the P-domain in the presence of different $Ca^{2+}$ concentrations yielded a dissociation constant $K_{D,Ca2+}$ of 1.09 ± 0.02 mM and a dissociation-rate constant $k_{off,Ca2+}$ of 27.5 ± 0.4 $s^{-1}$ employing 2D line shape analysis with TITAN. The resulting on-rate $k_{on,Ca2+}$ for the metal ion is approximately $2.8*10^4$ $M^{-1}s^{-1}$. The left panels in (d) show sections of the experimental methyl TROSY spectra, and the right panels show the simulated spectra. Spectra were acquired at 298 K on a 600 MHz spectrometer with cryo probe using 52 µM or 58 µM MILVA labeled MNV CW1 P-domain at pH* values of 3.9 (20 mM acetate-$d_4$, 100 mM NaCl) or of 7.5 (20 mM BisTris-$d_{19}$, 100 mM NaCl). A structural model of the P-domain by Nelson et al., 2018, (pdb 6e47) was used for the illustration.

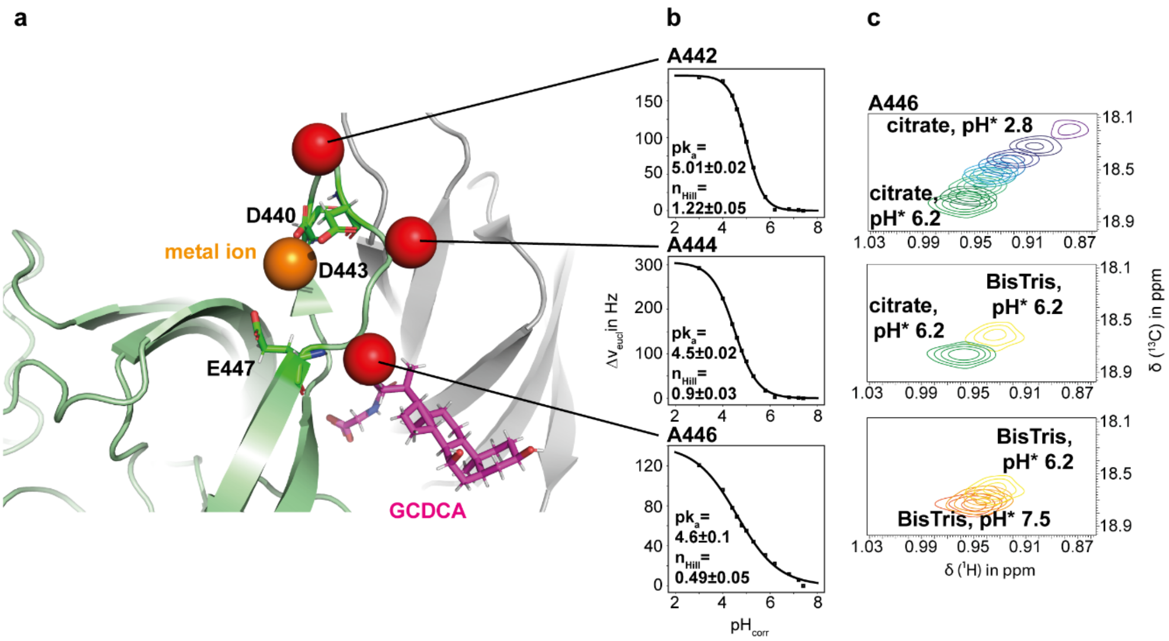


**Figure S14: Protonation of G'H' loop parallels with dimer formation.** (a) shows the G'H' loop which is involved in metal ion and GCDCA binding (pdb 6e47, Nelson et al., 2018). Both events lead to dimerization of the P-domain. The loop contains three acidic residues (Asp440, Asp443 and Glu447). Alanine methyl groups within the loop are highlighted as red spheres. The corresponding resonances show CSPs due to the alteration of the pH* value. CSPs can be used to fit the Henderson-Hasselbalch equation (Croke et al., 2011, see the methods section) to derive ionization constants $pk_a$. Even though the CSPs probably do not report for protonation of a single side, it can be concluded that the G'H' loop changes its protonation state between $pH_{corr}$ 4 and 6 (b). (c) shows CSPs due to changes of pH* exemplarily for A446. As origin, the position of the resonance at a pH* of 2.8 ($pH_{corr}$ 3 ) was used and the pH* was increased to 6.2 ($pH_{corr}$ 6.2 ) (upper panel). BisTris buffer was used to follow CSPs in the neutral range (pH* 6.2 to 7.5 or $pH_{corr}$ 6.2 to 7.4 lower panel). To account for the impact of the different buffers, spectra were acquired at a pH* of 6.2 in both, citrate buffer and BisTris buffer (panel in the middle). For details on the acquisition of the respective NMR spectra, see Fig. S12 and the methods section. In addition to the information given there, saturating amounts of GCDCA were added to the samples.

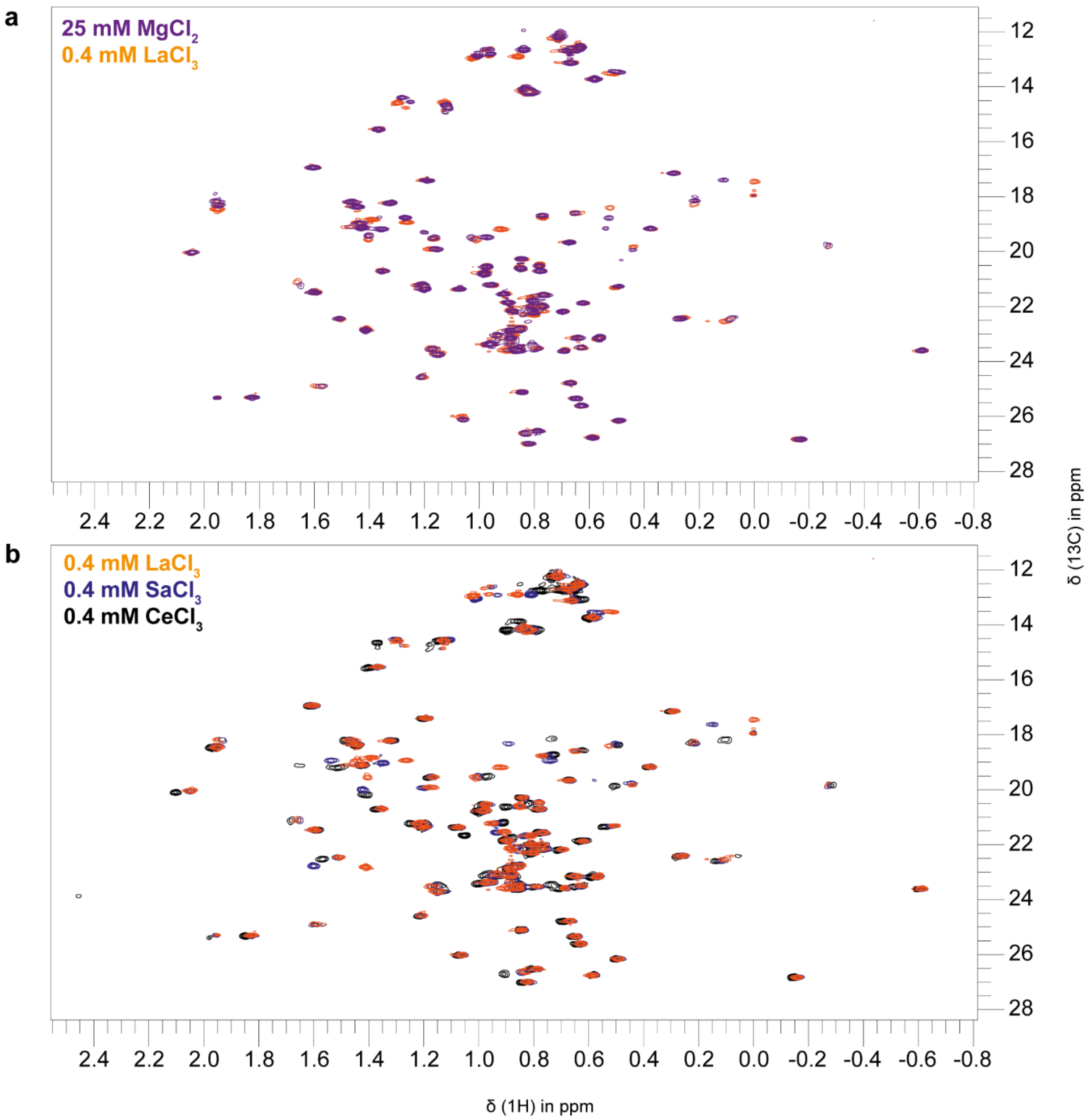


**Figure S15: P-domain methyl TROSY spectra with paramagnetic and diamagnetic metal ions.**

Methyl TROSY spectra were acquired at 298 K on a 600 MHz spectrometer with cryo probe using 38 μM MILVA labeled MNV CW1 P-domain. For sample conditions of the spectrum in presence of $MgCl_2$, see Fig. 2.

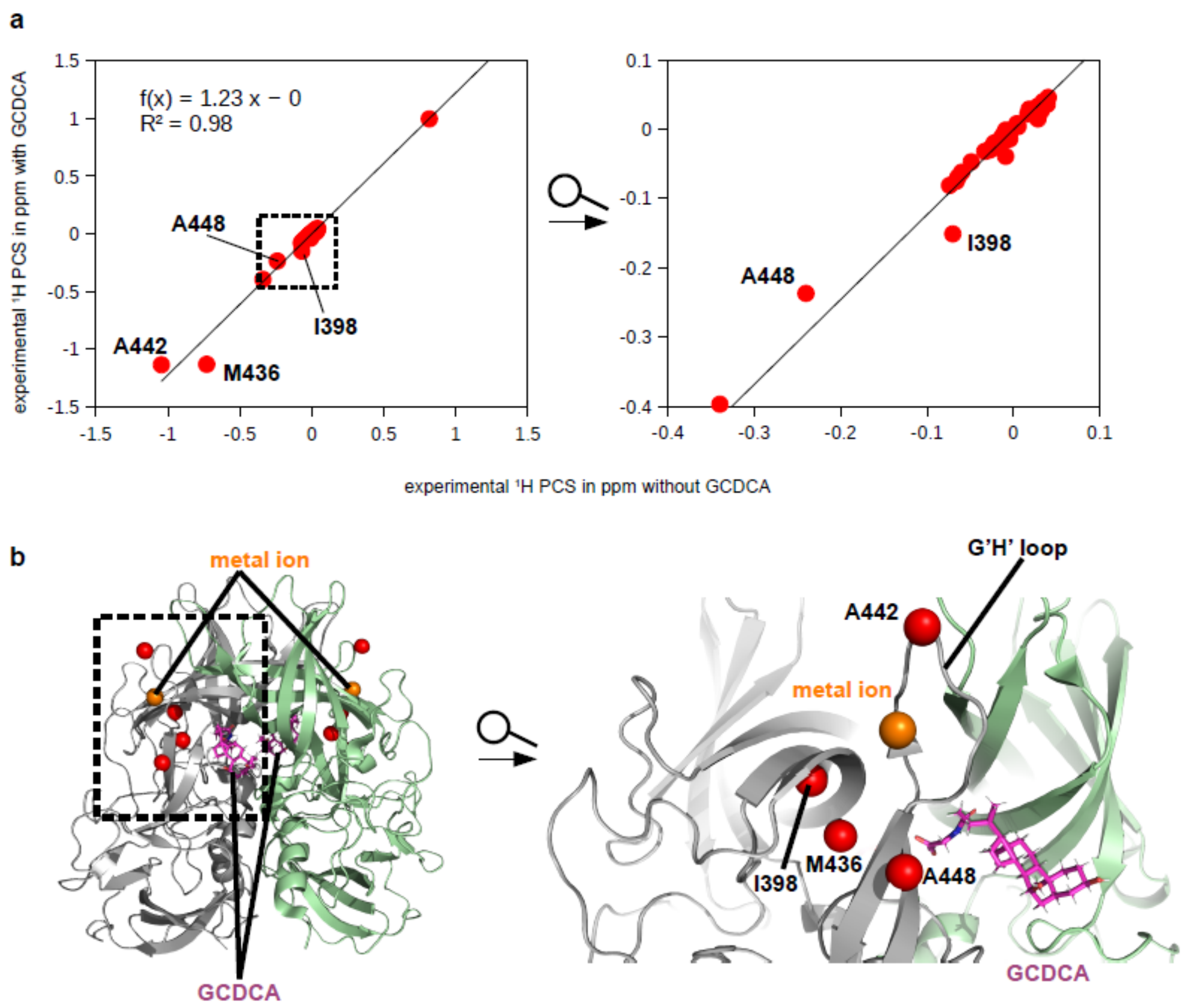


**Figure S16: $Ce^{3+}$ induced pseudo contact shifts (PCSs) in the absence and in the presence of GCDCA indicate no major conformational changes upon the addition of GCDCA.** (a) shows a correlation plot of PCSs in the presence and absence of GCDCA. PCSs are in general larger in the presence of GCDCA as can be seen by the slope of the line. This can be explained by stronger alignment in the presence of GCDCA as GCDCA enhances P-domain metal ion interactions. The correlation is good ($R^2$= 0.98), and outliers are located exclusively close to the metal ion binding site at the G'H' loop (b). As PCSs strongly depend on the distance to the metal ion, this region is susceptible for larger PCS differences. Slight rearrangements at the G'H' loop might be caused by GCDCA binding to it. No large-scale or global structural rearrangements by GCDCA binding to the metal ion-bound P-domain can be concluded. The structures with and without GCDCA must be highly similar. PCSs in the absence of GCDCA were obtained from [$^1H$,$^{13}C$] HMQC spectra shown in Fig. S15 PCSs in the presence of GCDCA are as shown in Maass et al.[4]. The structural model of Nelson et al., 2018 (pdb 6e47) was used to for the illustration.

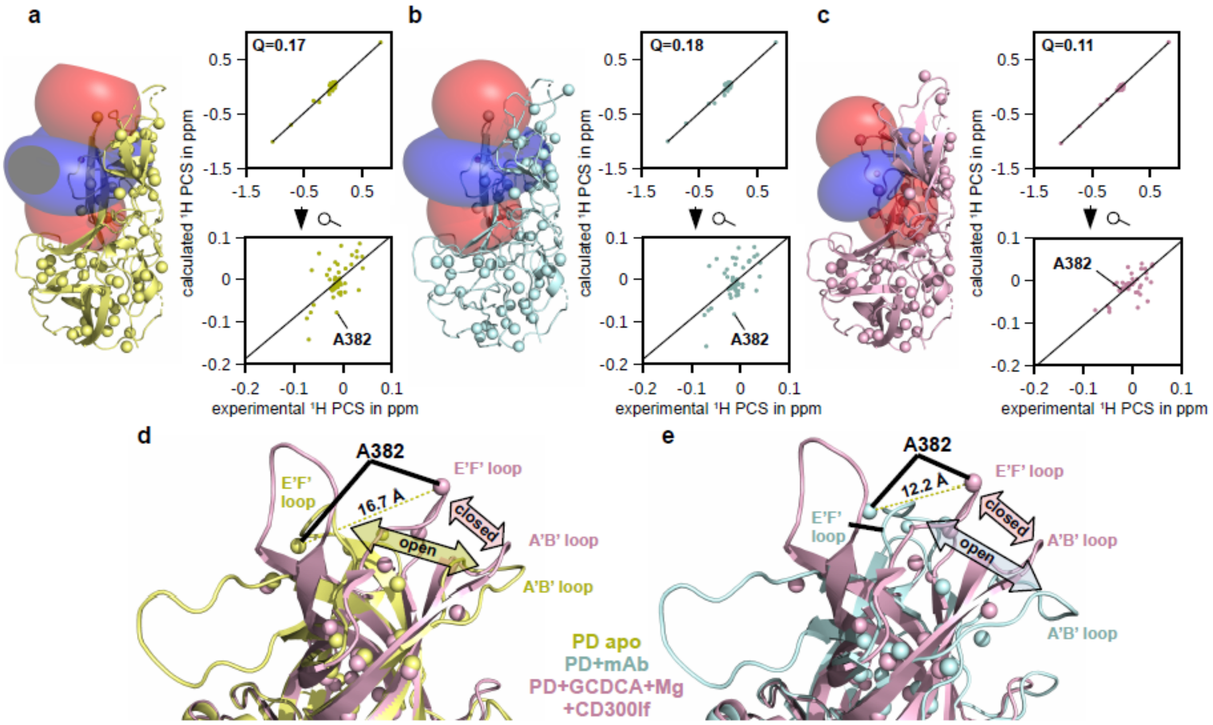


**Figure S17: P-domains in the presence of metal ions exists in a conformation that does not allow binding of neutralizing antibodies**. PCSs were measured in the presence of $CeCl_3$ (Fig. S15b) and Δχs and the position of paramagnetic centers were determined using structural models of the apo P-domain (a, pdb 3lq6, yellow), P-domain in the presence of a monoclonal antibody (mAb) (b, pdb 7l5j, cyan) and in the presence of GCDCA, metal ions and the CD300lf receptor (c, pdb 6e47, purple). Note that structural models of P-domain in complex with CD300lf and metal ions in the presence and absence of GCDCA are almost identical (pdb 6c6q, RSMD=0.248 $nm^2$). The best Q-factor was observed for the P-domain in the presence of GCDCA, metal ions, and CD300lf. For, e.g., Ala382, theoretical and measured PCSs do not match using structural models of the apo P-domain or of the P-domain in the presence of the mAb (a and b). Whereas the E'F' loop, where A382 is located, is in the open conformation for these two structural models, it is in a closed conformation in the presence of GCDCA, metal ions, and CD300lf (d, e). Note that the color coding in (d) and (e) is the same as in (a), (b), and (c). For Δχ tensor and paramagnetic center fitting, the structural model of P-domain in the presence of GCDCA, metal ions and the CD300lf receptor (c, pdb 6e47) was processed as explained in Maass et al.[4]. Identical atoms were included for the calculations with structural models of apo P-domain (a) and P-domain in complex with the mAb (b). See Tab. S3 for tensor and paramagnetic center parameters.

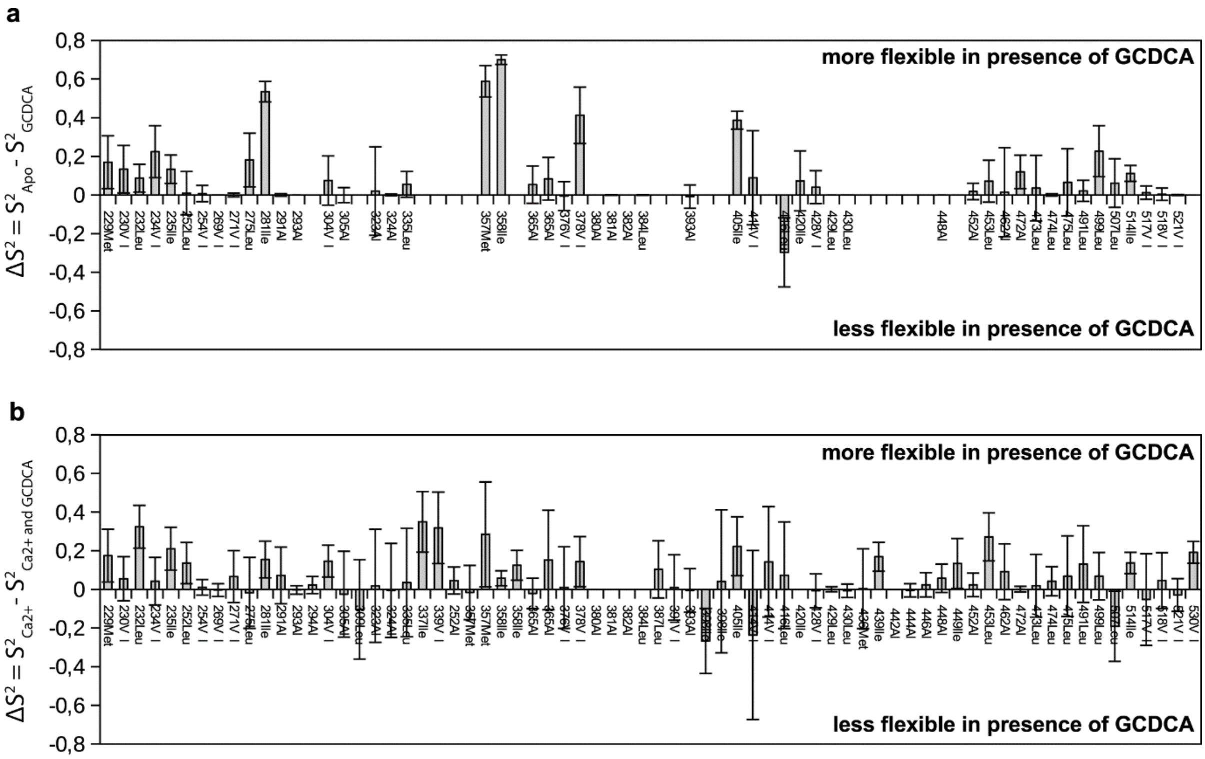


**Figure S18: Methyl group order parameters reflect GCDCA changing the dynamic profile of the apo P-domain upon binding to $Ca^{2+}$. (a)** apo P-domain (pH* 4.5) and **(b)** P-domain in the presence of $Ca^{2+}$ (pH* of 5.3). Bar plots of order parameter differences in the presence and absence of GCDCA. Notably, most methyl groups increase in flexibility due to GCDCA binding. The corresponding spectra were acquired at 298 K on a Bruker AV 600 MHz spectrometer equipped with a cryo-probe. Information about sample conditions is given in Table S8.

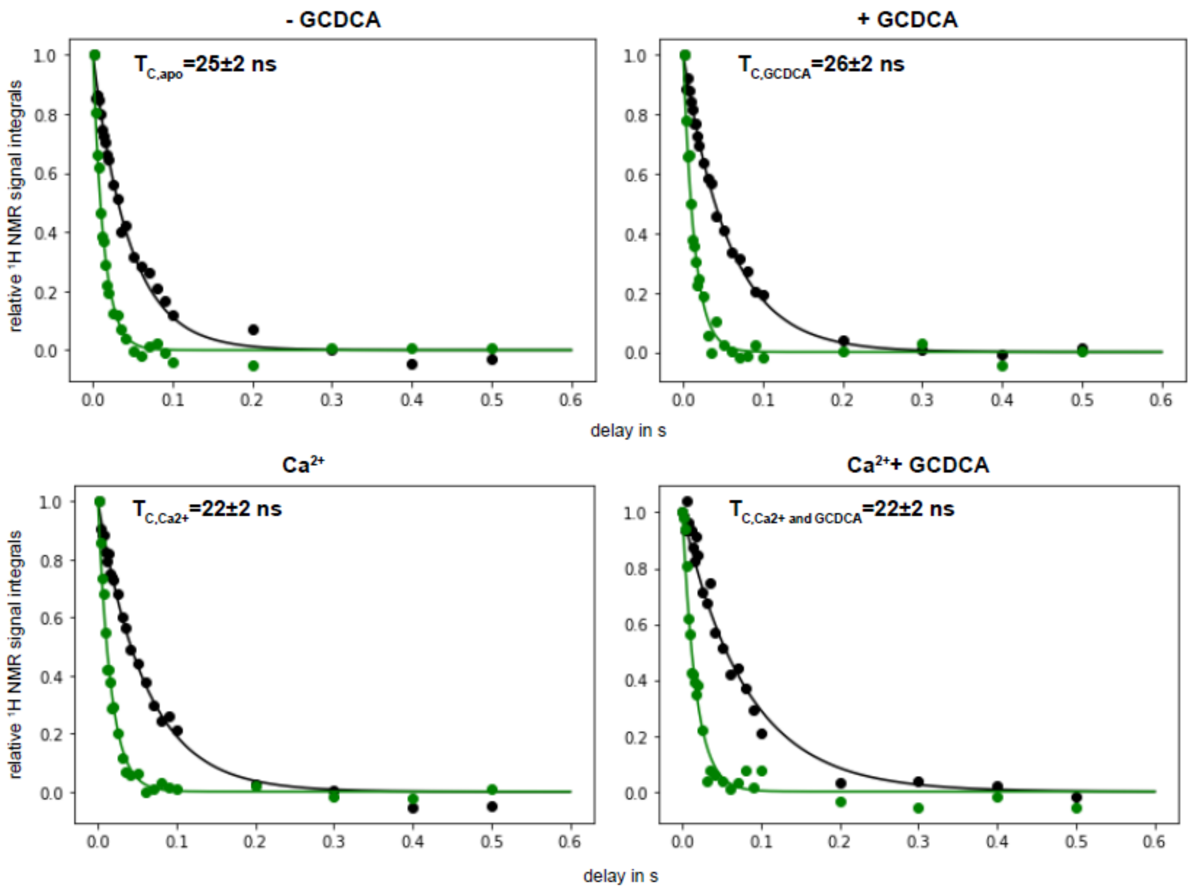


**Figure S19: TRACT experiments show identical molecular correlation times $\tau_C$ in the presence and absence of GCDCA at a pH* of 4.6 and at a pH* of 5.3 in the presence of $CaCl_2$.** Sample conditions, experimental details, and details on the analysis are given in Table S7.

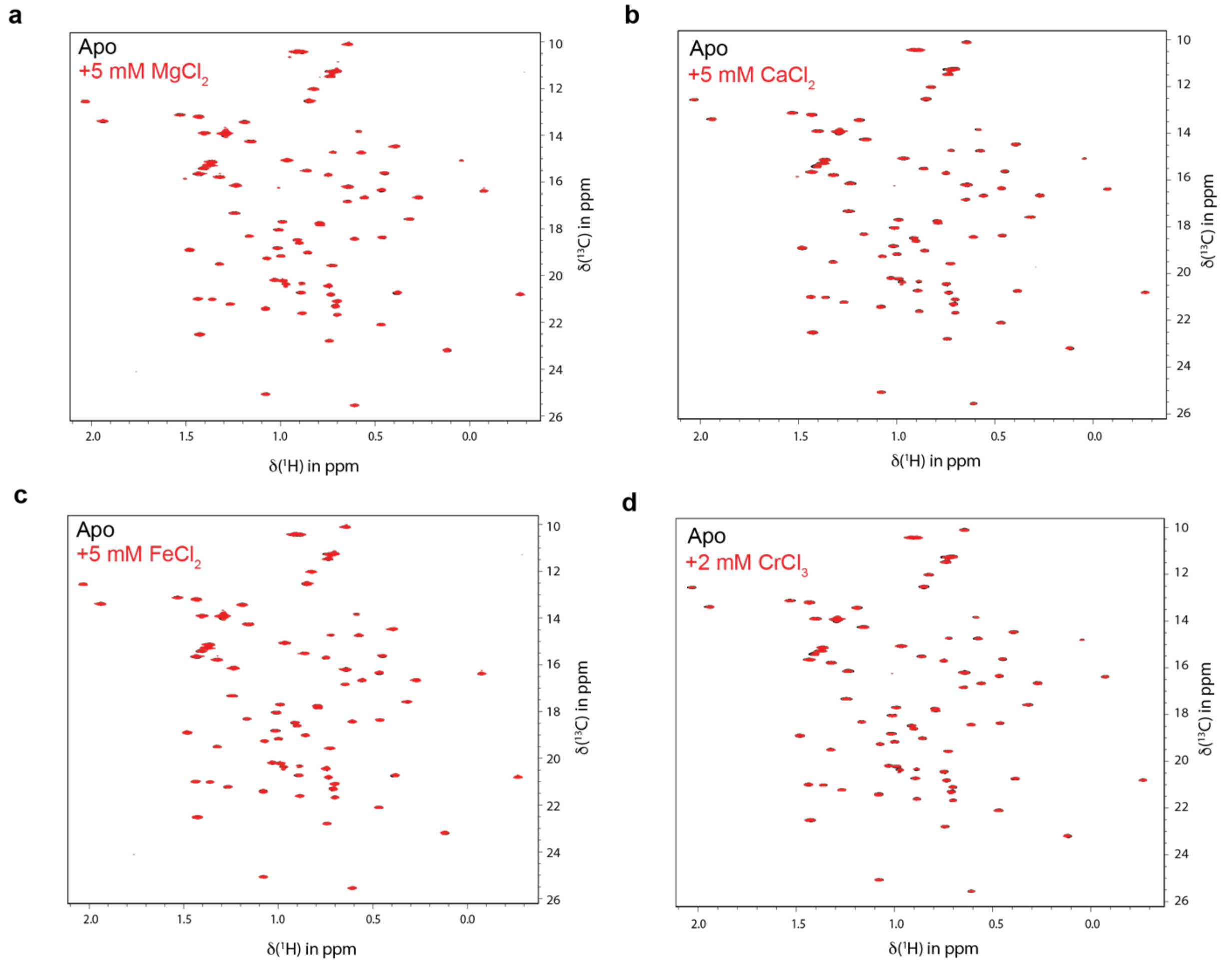


**Figure S20: $Mg^{2+}$, $Ca^{2+}$, $Fe^{2+}$ and $Cr^{3+}$ do not cause any perturbation in methyl TROSY spectra of GII.4 Saga P-dimers.** Superposition of methyl TROSY spectra of apo P-dimers (black) and P-dimers in the presence of **a)** 5 mM $MgCl_2$, **b)** 5 mM $CaCl_2$, **c)** 5 mM $FeCl_2$ and 10 mM ascorbic acid and **d)** 2 mM $CrCl_3$. The absence of any intensity changes or chemical shift perturbations (CSPs) and intensity changes shows that $Mg^{2+}$, $Ca^{2+}$, $Fe^{2+}$ and $Cr^{3+}$ do not bind to GII.4 Saga P-dimers.

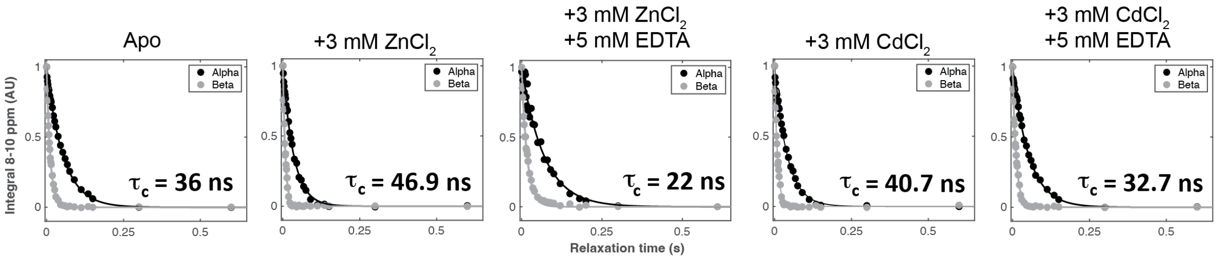


**Figure S21: TRACT experiments show changes in $\tau_C$ in upon titration of $ZnCl_2$ and $CdCl_2$, and after metal chelation with EDTA.** All experiments were measured at pH 6.88. Imidazole was used as pH reporter to ensure pH stability through titrations**.**

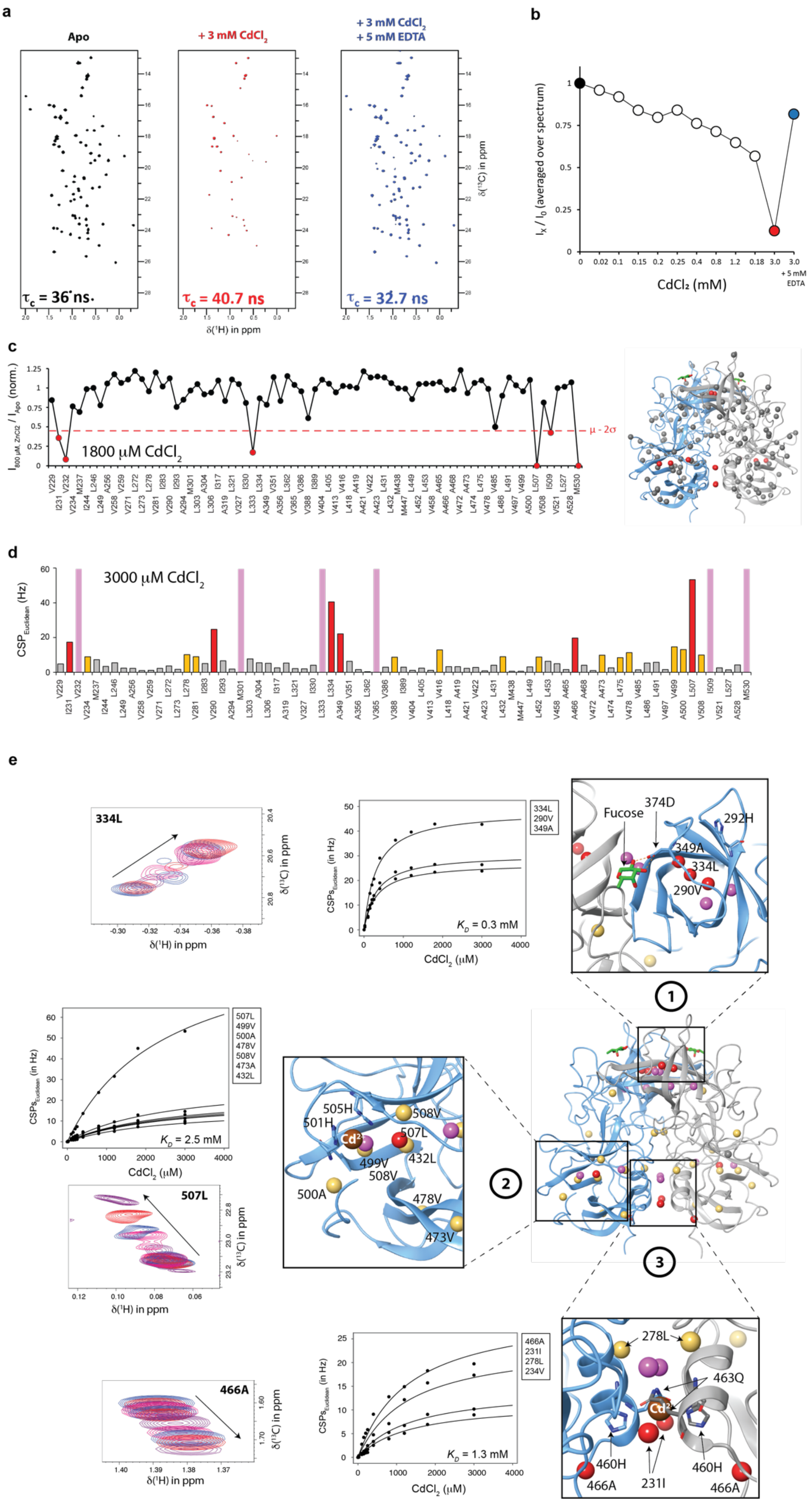

a
Apo
+ 3 mM CdCl2
+ 3 mM CdCl2
+ 5 mM EDTA
τc = 36 ns
τc = 40.7 ns
τc = 32.7 ns
δ(1H) in ppm
δ(13C) in ppm
b
Ix / I0 (averaged over spectrum)
CdCl₂ (mM)
+ 5 mM EDTA
c
I800 µM ZnCl2 / IApo (norm.)
1800 µM CdCl2
µ - 2σ
d
CSPEuclidean (Hz)
3000 µM CdCl2
e
334L
507L
466A
CSPsEuclidean (in Hz)
CdCl2 (µM)
KD = 0.3 mM
KD = 2.5 mM
KD = 1.3 mM
334L
290V
349A
507L
499V
500A
478V
508V
473A
432L
466A
231I
278L
234V
Fucose
374D
292H
349A
334L
290V
505H
501H
Cd2+
508V
507L
432L
499V
508V
500A
478V
473V
278L
463Q
460H
466A
231I
460H
466A
1
2
3

**Figure S22 Spectral effects upon addition of $CdCl_2$ to MILVA-labeled N373D huNoV GII.4 Saga P-dimers as observed in methyl TROSY spectra. a)** Methyl TROSY spectra of Saga P-dimers in the absence of $CdCl_2$ (black). Addition of 3 mM $CdCl_2$ (red) is associated with signal decrease accompanied by a light increase in $\tau_c$, suggesting protein aggregation. Signal intensities can be recovered by addition of 5 mM EDTA (blue), which is accompanied by a decrease in $\tau_c$ as compared to apo P-dimers. $\tau_c$ were experimentally obtained from $^{15}N$ TRACT experiments. **b)** Normalized sum of intensities of all cross peaks of methyl TROSY spectra of Saga P-dimers at increasing $CdCl_2$ concentrations and, finally, after the addition of 5 mM EDTA. **c)** *Left panel*: Methyl group specific decrease of methyl group signal intensities in a methyl TROSY spectrum of Saga P-dimers in the presence of 1800 mM $CdCl_2$. Red dots indicate a signal decrease of at least $\mu$-2$\sigma$. *Right panel:* Location of methyl groups showing a significant signal decrease of larger than $\mu$-2$\sigma$ upon addition of 1800 mM $CdCl_2$ mapped onto the crystal structure of Saga P-dimers. **d)** CSPs observed in methyl groups at 3000 mM $CdCl_2$ concentration. Methyl groups whose signals were broadened beyond detection in the HMQC spectra are indicated in pink. Orange and red denote CSPs over $\mu$+$\sigma$ and $\mu$+2$\sigma$, respectively. Gray indicates non-significant CSPs. **e)** Epitope mapping on Saga P-dimers crystal structure of significant CSPs observed at 3000 mM $CdCl_2$ concentration. Color code like in (d). Fitting of the law of mass action to the binding isotherms yields dissociation constants $K_D$ for the binding of $Cd^{2+}$ to the metal binding pockets 1, 2 and 3. Amino acid labels adjacent to binding isotherms are ordered according to largest to smallest CSPs at 3000 mM $CdCl_2$ concentration. A superimposition of spectra as a function of metal concentration is shown for the methyl group with the largest CSPs in each metal binding pocket. Metal positions are only approximated, and were obtained by superimposition with crystal structure pdb 76KV for binding site 3[5] or manually added in binding site 2. No fucose has been added to the sample but the position of the L-fucose residue in the crystal structure is shown (green sticks). For visualization, the crystal structure pdb 4X06 was used in all panels.

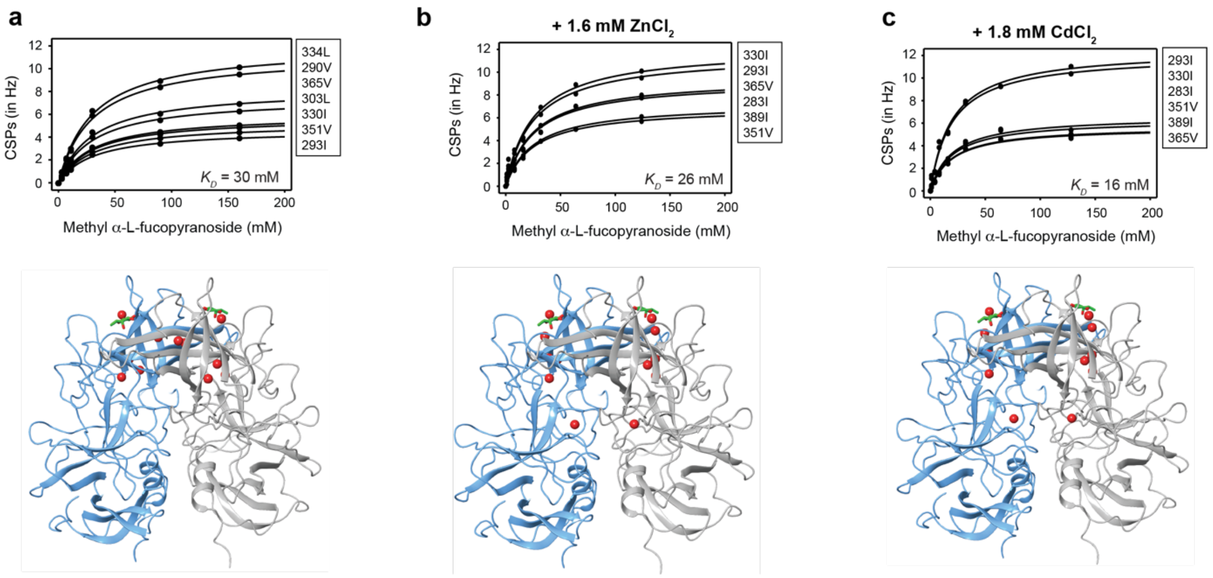


**Figure S23:** *Top panels* show binding isotherms and dissociation constants $K_D$ were obtained for the binding of methyl-α-L-fucopyranoside to N373D huNoV GII.4 Saga P-dimers in **a)** the absence of any bivalent metal ions, **b)** in the presence of 1.6 mM $ZnCl_2$ and **c)** in the presence of 1.8 mM $CdCl_2$. Only signals showing CSPs larger than $\mu+2\sigma$ at highest glycan concentrations were selected for fitting. Amino acid labels adjacent to binding isotherms are ordered according to largest to smallest CSPs at 160 mM methyl-α-L-fucopyranoside concentration. Metal concentrations were selected to prevent protein precipitation and were kept constant throughout the titration. *Bottom panels* show the location of the methyl groups used for calculation of $K_D$s as red balls (pdb 4X06).

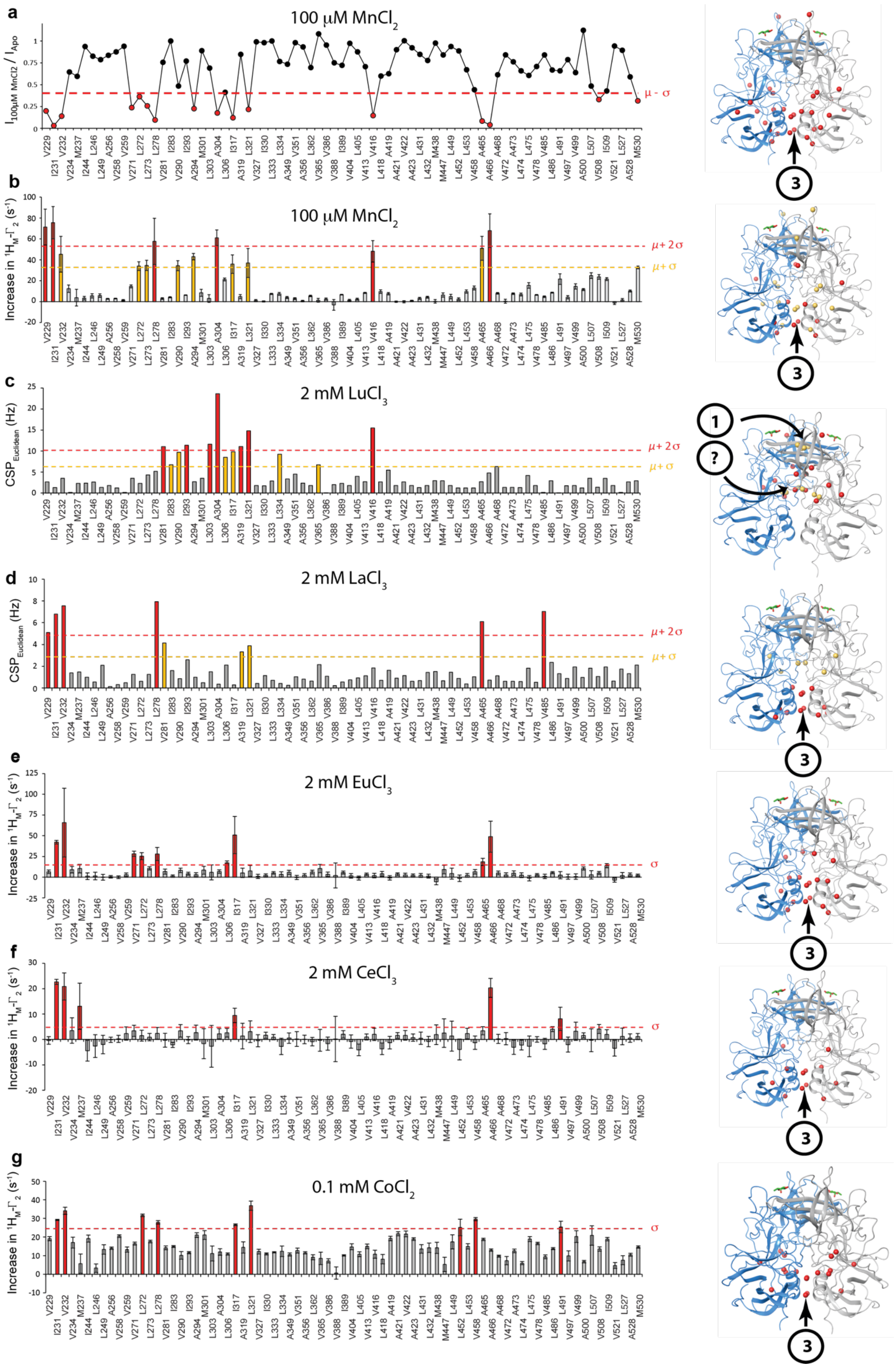
a
100 μM MnCl2
I100μM MnCl2 / IApo
μ - σ
b
100 μM MnCl2
Increase in 1HM-Γ2 (s-1)
μ+ 2σ
μ+ σ
c
2 mM LuCl3
CSPEuclidean (Hz)
μ+ 2σ
μ+ σ
d
2 mM LaCl3
CSPEuclidean (Hz)
μ+ 2σ
μ+ σ
e
2 mM EuCl3
Increase in 1HM-Γ2 (s-1)
σ
f
2 mM CeCl3
Increase in 1HM-Γ2 (s-1)
σ
g
0.1 mM CoCl2
Increase in 1HM-Γ2 (s-1)
σ
V229 I231 V232 V234 M237 I244 L246 L249 A256 V258 V259 V271 L272 L273 L278 V281 I283 V290 I293 A294 M301 L303 A304 L306 I317 A319 L321 V327 I330 L333 L334 A349 V351 A356 L362 V365 V386 V388 I389 V404 L405 V413 V416 L418 A419 A421 V422 A423 L431 L432 M438 M447 L449 L452 L453 V458 A465 A466 A468 V472 A473 L474 L475 V478 V485 L486 L491 V497 V499 A500 L507 V508 I509 V521 L527 A528 M530
1
?
3

**Figure S24: Spectral effects upon addition of $MnCl_2$, $CoCl_2$ and lanthanoids. a)** *Left panel*: Methyl group specific decrease of signal intensities in methyl TROSY spectra of Saga P-dimers upon addition of 100 $\mu$M $MnCl_2$. Red dots indicate a signal decrease of at least $\mu$-$\sigma$. *Right panel*: Location of methyl groups with a significant signal reduction of larger than $\mu$-$\sigma$ upon addition of 100 $\mu$M $MnCl_2$ mapped onto the crystal structure of GII.4 Saga P-dimers. $MnCl_2$ binds to a region labeled as region 3. This region is different from the binding regions 1 and 2 for $ZnCl_2$ and $CdCl_2$ (Figs. S4 and S5). **b)** *Left panel*: Increase in $^1H_M$-$\Gamma_2$ rates of methyl groups in P-dimers in the presence of 100 $\mu$M $MnCl_2$. Yellow and red indicate an increase of $\mu$+$\sigma$ and $\mu$+2$\sigma$, respectively. For the calculation of errors see the Methods section. *Right panel*: Location of methyl groups showing a significant ($\mu$+$\sigma$ and $\mu$+2$\sigma$) increase in $^1H_M$-$\Gamma_2$ rates upon addition of 100 $\mu$M $MnCl_2$ mapped onto the crystal structure of GII.4 Saga P-dimers. **c)**, **d)** *Left panels:* Chemical shift perturbances (CSPs) observed at 2 mM $LuCl_3$ (c) and 2 mM $LaCl_3$ (d) concentrations. Yellow and red indicate an increase CSPs larger than $\mu$+$\sigma$ and $\mu$+2$\sigma$, respectively. *Right panels:* Location of methyl groups showing significant CSPs. The question mark in (c) denotes that CSPs were observed for methyl groups not located in metal binding regions 1, 2 or 3. **e)**, **f), g)** Increase in $^1H_M$-$\Gamma_2$ rates of methyl groups in P-dimers in the presence of 2 mM $EuCl_3$ (e), 2 mM of $CeCl_3$ (f) and 0.1 mM $CoCl_2$ (g). Red indicates an increase of $\sigma$ in $^1H_M$-$\Gamma_2$ rates. No fucose has been added to the samples. However, the position of the L-fucose residue is shown in every crystal structure for illustrative purposes (green sticks). All GII.4 crystal structures correspond to pdb 4X06.

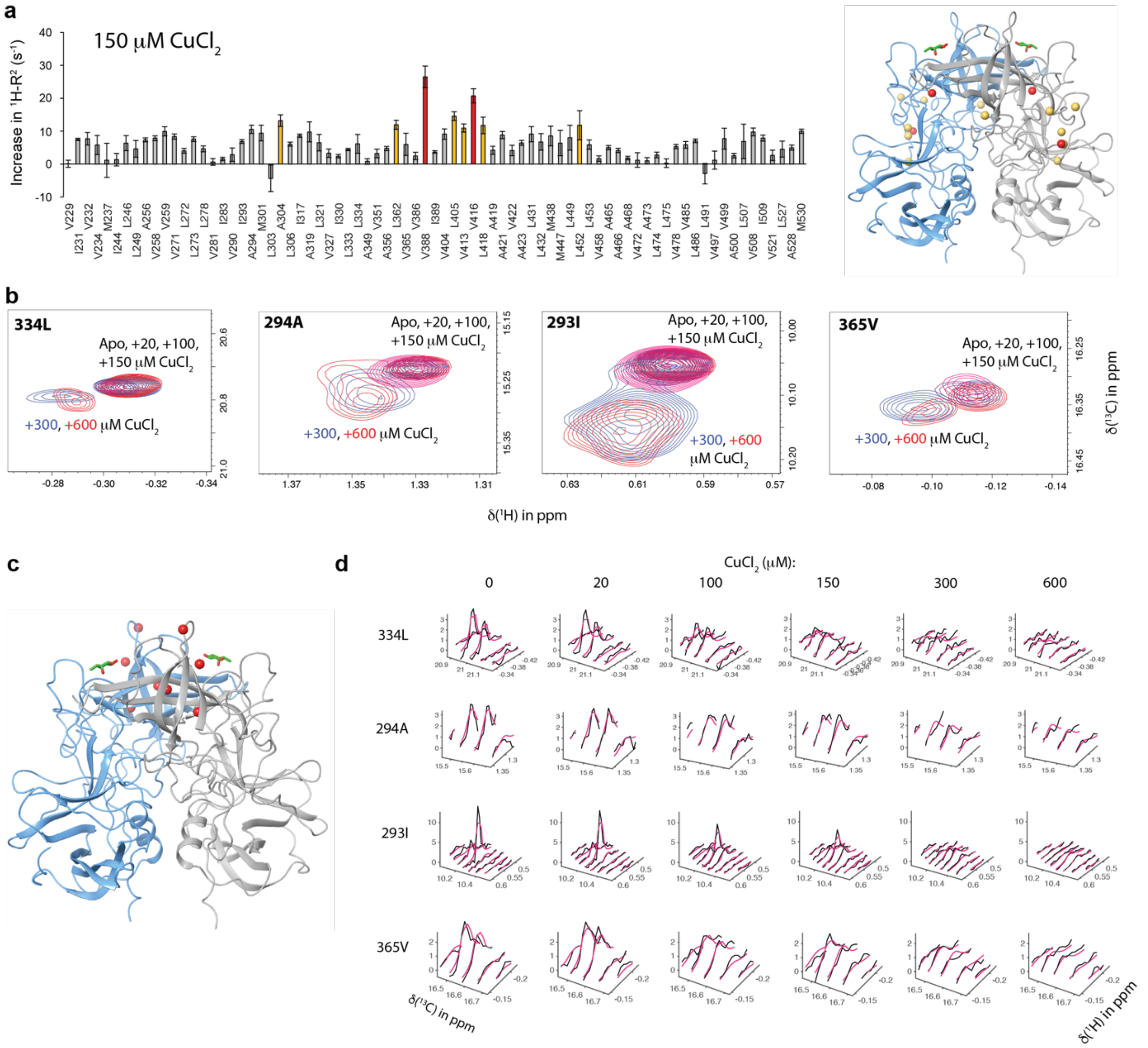


**Figure S25: Spectral effects upon addition of $CuCl_2$ to MILVA-labeled N373D huNoV GII.4 Saga P-dimers as observed in methyl TROSY spectra. a)** *Left panel*: Increase of $^{1}H_{M}$-$\Gamma_2$ rates of methyl groups in Saga P-dimers at 150 μM $CuCl_2$ concentration. Yellow and red indicate an increase of $\mu+\sigma$ and $\mu+2\sigma$, respectively. For the calculation of errors see the Methods section. *Right panel*: Location of methyl groups showing a significant increase of $^{1}H_{M}$-$\Gamma_2$ rates upon addition of 150 μM $CuCl_2$ mapped onto the crystal structure of GII.4 Saga P-dimers. **b)** Superimposition of methyl TROSY spectra of methyl groups selected for line shape fitting with TITAN. Note that free and metal bound protein states are in fast-to-intermediate exchange on the NMR chemical shift time scale. Indicated are the metal concentrations. **c)** Red balls indicate the location of the methyl groups from (b) in the crystal structure of Saga P-dimers (pdb 4X06). **d)** 2D line shape analyses of methyl TROSY cross peaks with TITAN. Black and magenta spectra correspond to the measured and simulated spectra, respectively. The overlay visually reflects the good quality of the fit.

**Table S1: $^1$H and $^{13}$C chemical shifts δ of the apo MNV P-domain and CSPs due to the presence of saturating amounts of $Mg^{2+}$.**

The assignment of $^{13}$C methyl chemical shifts of the apo MNV P-domain was based on a previous assignment by Creutznacher et al.[2] (see Table S3 of that reference). Total CSPs ($\Delta\nu_{Eucl}$) were calculated using Eq. 4 (main text). Rows colored in gray correspond to signals where CSPs could not be determined. Rows colored in red designate resonances where CSPs were assumed to be larger than 20 Hz since no corresponding peak could be identified upon addition of $Mg^{2+}$. The chemical shifts and CSPs tabulated for Ile405 correspond only to one of the cross peaks (see legend to Figure S1 and cf. Table S3 of Creutznacher et al.[2]) observed. The chemical shifts of L359 and L384 of the apo form are not tabulated in this table but are given in Table S3 of Creutznacher et al.[2] (see also legend to Figure S1). See legend to Figure 2 of the main text for more details on the sample and acquisition conditions.

| | Apo P-domain | | P-domain with $Mg^{2+}$ bound | | CSP / Hz | | |
|---|---|---|---|---|---|---|---|
| Amino acid | δ($^1$H) / ppm | δ($^{13}$C) / ppm | δ($^1$H) / ppm | δ($^{13}$C) / ppm | $\Delta\nu_H$ | $\Delta\nu_C$ | $\Delta\nu_{Eucl}$ |
| 229Met | 1.61 | 16.9 | 1.6 | 16.9 | 1.5 | 2.1 | 2.6 |
| 230Val | 0.7 | 22.2 | 0.7 | 22.2 | 0.7 | 1.6 | 1.7 |
| 232Leu | 0.31 | 22.5 | 0.27 | 22.4 | 23.8 | 13 | 27.1 |
| 234Val | 0.84 | 20.2 | 0.85 | 20.3 | 1.7 | 7.3 | 7.5 |
| 235Ile | 0.58 | 13.7 | 0.58 | 13.7 | 1 | 0.3 | 1 |
| 252Leu | 0.49 | 26.1 | 0.49 | 26.2 | 3.2 | 2.3 | 4 |
| 254Val | 0.38 | 19.2 | 0.38 | 19.2 | 0.6 | 1.9 | 2 |
| 269Val | 1.34 | 20.7 | 1.35 | 20.7 | 5 | 0.5 | 5 |
| 271Val | 0.63 | 23.1 | 0.64 | 23.1 | 7.1 | 3.9 | 8.1 |
| 275Leu | 0.7 | 23.7 | 0.69 | 23.6 | 2.6 | 5.3 | 6 |
| 281Ile | 0.7 | 12 | 0.71 | 12.2 | 7.9 | 25.1 | 26.3 |
| 291Ala | 1.22 | 24.6 | 1.21 | 24.6 | 4 | 1.9 | 4.5 |
| 293Ala | 1.15 | 23.8 | 1.15 | 23.7 | n.d. | n.d. | n.d. |
| 294Ala | n.d. | n.d. | 1.21 | 21.2 | n.d. | n.d. | n.d. |
| 304Val | 0.85 | 20.6 | 0.85 | 20.6 | 0.6 | 3.7 | 3.7 |
| 305Ala | 0.08 | 22.4 | 0.09 | 22.5 | n.d. | n.d. | n.d. |
| 309Leu | 0.62 | 23.4 | 0.63 | 23.5 | 4.6 | 10.7 | 11.6 |
| 310Ile | n.d. | n.d. | 0.81 | 14.2 | n.d. | n.d. | n.d. |
| 323Ala | 0.22 | 18 | 0.21 | 18.1 | 3.1 | 18.9 | 19.1 |
| 324Ala | 1.16 | 19.5 | 1.17 | 19.5 | 3.4 | 6 | 6.9 |
| 335Leu | 1 | 26.4 | 1.06 | 26.1 | 35.9 | 37.1 | 51.6 |
| 337Ile | 0.67 | 13.2 | 0.67 | 13.1 | 1.3 | 5 | 5.2 |
| 339Val | n.d. | n.d. | 1.03 | 19.5 | n.d. | n.d. | >20 |
| 352Val | n.d. | n.d. | 0.77 | 18.7 | n.d. | n.d. | >20 |
| 357Met | 1.94 | 18 | 1.95 | 18.3 | 6.2 | 48.7 | 49.1 |
| 357Met | 1.95 | 17.8 | 1.96 | 18.2 | 2.5 | 47.3 | 47.4 |
| 358Ile | 0.96 | 12.6 | 1.01 | 12.9 | 29 | 40.3 | 49.7 |
| 358Ile | 0.96 | 12.6 | 0.96 | 12.8 | 0.1 | 29 | 29 |
| 365Ala | n.d. | n.d. | 1.4 | 19.4 | n.d. | n.d. | >20 |
| 365Ala | 1.22 | 19.1 | 1.2 | 19.3 | 14.4 | 34.8 | 37.6 |
| 374Val | 0.47 | 19.9 | 0.44 | 19.9 | 15.9 | 11.9 | 19.8 |
| 374Val | 0.49 | 20.1 | 0.48 | 20.3 | 2.9 | 37.8 | 37.9 |
| 376Ala | 1.54 | 24.9 | 1.57 | 24.9 | n.d. | n.d. | n.d. |
| 378Val | n.d. | n.d. | 0.96 | 21.2 | n.d. | n.d. | >20 |
| 380Ala | 1.61 | 21.4 | 1.6 | 21.5 | 7.5 | 10.5 | 12.9 |

| | | | | | | | |
|---|---|---|---|---|---|---|---|
| 381Ala | 1.43 | 19.1 | 1.43 | 19.1 | 0 | 2.5 | 2.5 |
| 382Ala | 1.07 | 21.3 | 1.07 | 21.4 | 0.4 | 7.8 | 7.8 |
| 387Val | n.d. | n.d. | 0.81 | 21.8 | n.d. | n.d. | n.d. |
| 391Val | 0.63 | 18.3 | 0.65 | 18.6 | 13.5 | 43.6 | 45.6 |
| 393Ala | -0.28 | 19.8 | -0.27 | 19.7 | n.d. | n.d. | n.d. |
| 398Ile | n.d. | n.d. | 1.28 | 14.4 | n.d. | n.d. | >20 |
| 398Ile | n.d. | n.d. | 1.25 | 14.5 | n.d. | n.d. | >20 |
| 405Ile | 1.11 | 14.7 | 1.12 | 14.7 | 4.3 | 11.8 | 12.5 |
| 414Val | 0.54 | 19 | 0.53 | 18.8 | 4.1 | 36.6 | 36.8 |
| 414Val | 0.55 | 19.2 | 0.54 | 19.2 | 3.1 | 6.1 | 6.9 |
| 416Leu | n.d. | n.d. | 0.81 | 23.53 | n.d. | n.d. | n.d. |
| 420Ile | 0.83 | 14 | 0.83 | 14.1 | 2.2 | 8.6 | 8.8 |
| 428Val | 0.98 | 20.8 | 0.98 | 20.8 | 0.7 | 3.2 | 3.3 |
| 429Leu | 0.83 | 26.6 | 0.83 | 26.6 | 1.1 | 2 | 2.3 |
| 430Leu | 0.82 | 27 | 0.82 | 27 | 0.8 | 3.2 | 3.3 |
| 436Met | 0.01 | 16.9 | 0.11 | 17.4 | 58.4 | 71.4 | 92.2 |
| 439Ile | n.d. | n.d. | 0.84 | 12.6 | n.d. | n.d. | >20 |
| 442Ala | n.d. | n.d. | 1.41 | 22.9 | n.d. | n.d. | >20 |
| 444Ala | 1.25 | 18.9 | 1.27 | 18.8 | 12.1 | 14.9 | 19.2 |
| 446Ala | n.d. | n.d. | 0.97 | 19.5 | n.d. | n.d. | >20 |
| 448Ala | 1.14 | 19.9 | 1.16 | 19.9 | 7.3 | 3.7 | 8.1 |
| 449Ile | 0.44 | 13.7 | 0.49 | 13.5 | 31.4 | 32.6 | 45.3 |
| 449Ile | 0.46 | 13.7 | 0.51 | 13.4 | 30.7 | 38.2 | 49 |
| 452Ala | 2.04 | 20 | 2.04 | 20 | 0 | 3.5 | 3.5 |
| 453Leu | 0.96 | 23.4 | 0.96 | 23.4 | 0.9 | 1.1 | 1.4 |
| 462Ala | 1.46 | 18.2 | 1.47 | 18.2 | 4.4 | 2.4 | 5 |
| 472Ala | 0.76 | 21.6 | 0.76 | 21.6 | 2.4 | 2.1 | 3.2 |
| 473Leu | -0.6 | 23.6 | -0.61 | 23.6 | 3.2 | 4.3 | 5.4 |
| 474Leu | 0.65 | 25.4 | 0.65 | 25.4 | 0.4 | 0.8 | 0.9 |
| 475Leu | 0.63 | 25.6 | 0.63 | 25.6 | 2.8 | 3.1 | 4.1 |
| 491Leu | 0.78 | 26.6 | 0.79 | 26.5 | 2.2 | 4.5 | 5 |
| 497Ile | 0.66 | 12.7 | 0.67 | 12.7 | 3.3 | 1.7 | 3.7 |
| 499Leu | -0.17 | 26.8 | -0.17 | 26.8 | 1.5 | 0.1 | 1.5 |
| 507Leu | 0.59 | 26.8 | 0.59 | 26.8 | 0.3 | 1 | 1 |
| 514Ile | 0.63 | 12.6 | 0.64 | 12.6 | 5 | 1.6 | 5.2 |
| 517Val | 0.63 | 21.9 | 0.62 | 21.9 | 3.2 | 7.9 | 8.5 |
| 518Val | 0.78 | 20.7 | 0.78 | 20.7 | 0.4 | 2.5 | 2.5 |
| 521Val | 0.29 | 17.1 | 0.29 | 17.1 | 1.2 | 0.3 | 1.2 |
| 530Val | 1.2 | 21.4 | 1.2 | 21.4 | 0.9 | 3.3 | 3.4 |

**Table S2: Parameters from Δχ fitting for $Ce^{3+}$.**

| | | Apo P-domain | P-domain with mAb | P-domain with CD300lf, $Mg^{2+}$ and GCDCA* |
|---|---|---|---|---|
| pdb code | | 3lq6 (Taube et al., 2010) | 7l5j (Williams et al. 2021a) | 6e47 (Nelson et al. 2018) |
| $\Delta X_{ax}$ in $10^{-32}$ $m^3$ | | -2.81±0.36 | -2.60±0.45 | -1.325±0.018 |
| $\Delta X_{rh}$ in $10^{-32}$ $m^3$ | | -1.01±0.20 | -0.57±0.28 | -0.679±0.025 |
| Coordinates of origin in Å | X | 3.94±2.2 | 94.9±1.0 | -16.3±0.5 |
| | Y | -6.45±0.43 | 90.8±0.9 | 2.9±0.4 |
| | Z | -5.27±1.3 | 99.4±1.5 | -45.3±0.7 |
| Orientation of principal axis of tensor in ° | α | 20.3±5.7 | 40±15 | 109.88±0.19 |
| | β | 94.5±3.0 | 24.1±8.3 | 74.1±1.3 |
| | γ | 163.3±9.3 | 24±27 | 170.4±7.6 |
| Q-factor | | 0.17 | 0.18 | 0.11 |

*similar to structure with P-domain with CD300lf and $Mg^{2+}$ (RMSD = 0.248 $nm^2$)

**Table S3: Metal ion binding to huNoV GII.4 Saga P-dimers**

| Metal | Concentrations titrated (mM) | Binding detected | N° binding pockets observed | Binding parameters | | |
|---|---|---|---|---|---|---|
| | | | | Binding site 1 | Binding site 2 | Binding site 3 |
| $Zn^{2+}$ | 0, 0.15, 0.25, 0.35, 0.5, 0.8, 1.6, 3.0 | Yes | 2 | $K_{D,approx}$ = 3.5 ± 0.4 mM | $K_D$ = 1.0 ± 0.1 mM | - |
| $Cd^{2+}$ | 0, 0.02, 0.1, 0.15, 0.2, 0.25, 0.4, 0.8, 1.2, 1.8, 3.0 | Yes | 3 | $K_D$ = 0.28 ± 0.1 mM<br>$k_{off}$ = 360 ± 30 $s^{-1}$<br>$k_{on}$ = 1.29x$10^6$ $M^{-1}s^{-1}$ | $K_D$ = 2.5 ± 0.2 mM | $K_D$ = 1.3 ± 0.1 mM |
| $Cu^{2+}$ | 0, 0.02, 0.1, 0.15, 0.3, 0.6, 0.9 | Yes | 1 | Not clearly characterized, mapped by PREs.<br>$K_D$ = 0.31 ± 0.08 mM; $k_{off}$ = 80.7 ± 30 $s^{-1}$; $k_{on}$ = 2.58x$10^5$ $M^{-1}s^{-1}$ | | |
| $Mn^{2+}$ | 0.1 | Yes | 1 | - | - | Mapped by PRE |
| $Co^{2+}$ | 0.1 | Yes | 1 | - | - | Mapped by PRE |
| $Ca^{2+}$ | 5 | No | - | - | - | - |
| $Fe^{2+}$ | 5 | No | - | - | - | - |
| $Mg^{2+}$ | 5 | No | - | - | - | - |
| $Cr^{3+}$ | 2 | No | - | - | - | - |
| $La^{3+}$ | 2 | Yes | - | - | - | Mapped by CSPs |
| $Lu^{3+}$ | 2 | Yes | 1 or 2 | Mapped by CSPs | - | - |
| $Eu^{3+}$ | 2 | Yes | 1 | - | - | Mapped by PRE |
| $Ce^{3+}$ | 2 | Yes | 1 | - | - | Mapped by PRE |

The locations of metal binding pockets were mapped by CSPs if not stated otherwise.

**Table S4: Primers for site-directed mutagenesis of MNV P-domain.** All primers were obtained from Eurofins.

| Mutant | Primer Sequence |
|---|---|
| D410A | Forward: 5’-CGAATACAACGCTGGCCTGCTCG-3’ |
| | Reverse: 5’-CGAGCAGGCCAGCGTTGTATTCG-3’ |
| D440A | Forward: 5’-GCGCCAGATTGCTACTGCGGATG-3’ |
| | Reverse: 5’-CATCCGCAGTAGCAATCTGGCGC-3’ |

**Table S5: Final concentrations of precursors for [$^1$H,$^{13}$C] MILVA and MI*LVA methyl group labeling.**

| Labelling scheme | Precursor | Concentration |
|---|---|---|
| $L^{ProS}$ , $V^{ProS}$ (L, V) | 2-$^{13}$C-methyl-4-$d_3$-acetolactate (in-house synthesis) | 195 mg/L |
| I | 2-ketobutyricacid-4-$^{13}$C-3, 3-$d_2$ (in-house synthesis) | 72 mg/L |
| I* | 2-ketobutyricacid-$^{13}C_4$ (Sigma Aldrich) | 72 mg/L |
| A | L-alanine-$^{13}$C-$d_2$ (Cambridge Isotope Laboratories) | 0.6 g/L |
| | succinate-$d_4$ (Cambridge Isotope Laboratories) | 3.75 g/L |
| M | L-methionine-(methyl-$^{13}$C) (Sigma Aldrich) | 130 mg/L |

Note that I* labelling leads to coupling in the $^{13}$C dimension of [$^1$H,$^{13}$C] HMQC spectra. The labelling was used as the corresponding precursor was available in large quantities.

**Table S6: Buffers used for analytical SEC.**

| Equilibration and running buffer | MNV P-domain concentration in µM | Chromatograms shown in |
|---|---|---|
| 20 mM sodium acetate, 100 mM NaCl (pH 5.3) | 5 | Fig. 2b, green |
| 20 mM sodium acetate, 100 mM NaCl, 5 mM $MgCl_2$ (pH 5.3) | 5 | Fig. 2b, blue |
| 20 mM sodium acetate, 100 mM NaCl, 25 mM $MgCl_2$ (pH 5.3) | 5 | Fig. 2b, magenta |
| 20 mM citric acid, 100 mM NaCl (pH 4) | 21 | Fig. S12b, red |
| 20 mM citric acid, 100 mM NaCl (pH 6.2) | 21 | Fig. S12b, black |

**Table S7: Sample conditions of TRACT experiments.**

| Sample | Shown in |
|---|---|
| 140 µM P-domain, 20 mM soudium acetate, 100 mM NaCl, 30 mM $CaCl_2$, 10 % $D_2O$ (pH* 5.3) | Fig. S 19 |
| 133 µM P-domain, 20 mM soudium acetate, 100 mM NaCl, 30 mM $CaCl_2$, 500 µM GCDCA, 10 % $D_2O$ (pH* 5.3) | Fig. S 19 |
| 81 µM P-domain, 20 mM citric acid, 100 mM NaCl, 10% $D_2O$ (pH*4.6) | Fig. S 19 |
| 177 µM P-domain, 20 mM citric acid, 100 mM NaCl, 500 µM GCDCA, 10% $D_2O$ (pH*4.6) | Fig. S 19 |

**Table S8: Sample conditions for [$^{1}H$,$^{13}C$] HMQC and HSQC spectra used for order parameter determination.**

| Sample | Shown in |
|---|---|
| 143 µM P-domain, 20 mM soudium acetate, 100 mM NaCl, 29 mM $CaCl_2$ ($pH_{corr}$ 5.3) | Fig. S18 |
| 136 µM P-domain, 20 mM soudium acetate, 100 mM NaCl, 29 mM $CaCl_2$, 500 µM GCDCA ($pH_{corr}$ 5.3) | Fig. S18 |
| 180 µM P-domain, 20 mM citric acid, 100 mM NaCl ($pH_{corr}$ 4.6) | Fig. S18 |
| 171 µM P-domain, 20 mM citric acid, 100 mM NaCl, 500 µM GCDCA ($pH_{corr}$ 4.6) | Fig. S18 |

### Supplementary methods - Estimating methyl group order parameters

#### NMR spectroscopy

TRACT experiments[6] were carried out using [U-$^2$H,$^{15}$N] and [$^1$H,$^{13}$C] MILVA methyl group labeled MNV P-domain proteins. Buffers and protein concentrations are given in Table 7. The relaxation delay was set to 2s. 16 scans were measured for P-domain at a pH* of 5.3. 64 scans were measured for P-domain at s pH* of 4.6. The pulse sequence was repeated with 25 increasing delays up to 0.5 s (see respective figures).

Adapted versions of [$^1$H,$^{13}$C] HMQC and HSQC experiments and HMQC experiments with an additional delay to obtain $R^S_{MQ}$ relaxation rates for determination of methyl group order parameters[7] were carried out using [$^1$H,$^{13}$C] MILVA methyl group labeled MNV P-domain proteins. Sample compositions are given in Table S8. The relaxation delay was set to 1.5 s. The sweep width and spectral window centers were set to 3.75 ppm and 0.75 ppm for the direct dimension and to 18 ppm and to 17.5 ppm for the indirect dimension. For both dimensions, 512 increments were acquired. The number of scans was 8 except of the sample containing $CaCl_2$ and GCDCA, where 80 scans were acquired for HMQC and HSQC spectra and 32 for HMQC experiments with additional delays.

**Calculating S2 order parameters**

Methyl group order parameters were determined as suggested by Tugarinov and Kay[7], using the equations given below with a more extensive error propagation analysis.

$$I_{HSQC}(t_1, t_2) =$$

$$\left[\frac{9}{4} * \exp\left(-2\tau R_{2,H}^{F}\right) + \frac{9}{4} * \exp\left(-2\tau R_{2,H}^{S}\right)\right]$$

$$* \exp\left(-2\tau R_{2,H}^{F}\right) * \exp\left(-t_1 R_{2,C}^{F}\right) * \exp\left(-t_2 R_{2,H}^{F}\right)$$

$$+ [\frac{9}{4} * \exp\left(-2\tau R_{2,H}^{F}\right) - \frac{3}{4} * \exp\left(-2\tau R_{2,H}^{S}\right)]$$

$$* exp\left(-2\tau R_{2,H}^{F}\right) * \exp\left(-t_1 R_{2,C}^{S}\right) * \exp\left(-t_2 R_{2,H}^{F}\right)$$

$$+ \left[\frac{9}{4} * \exp\left(-2\tau R_{2,H}^{F}\right) + \frac{9}{4} * \exp\left(-2\tau R_{2,H}^{S}\right)\right]$$

$$* \exp\left(-2\tau R_{2,H}^{S}\right) * \exp\left(-t_1 R_{2,C}^{F}\right) * \exp\left(-t_2 R_{2,H}^{S}\right)$$

$$+ [-\frac{3}{4} * \exp\left(-2\tau R_{2,H}^{F}\right) + \frac{9}{4} * \exp\left(-2\tau R_{2,H}^{S}\right)]$$

$$* \exp\left(-2\tau R_{2,H}^{S}\right) * exp\left(-t_1 R_{2,C}^{S}\right) * exp\left(-t_2 R_{2,H}^{S}\right)$$

**(Eq. S1)**

$$I_{HMQC}(t_1, t_2) = 6 * \exp\left(-4\tau R_{2,H}^{F}\right) * \exp\left(-t_1 R_{MQ}^{F}\right) * \exp\left(-t_2 R_{2,H}^{F}\right)$$

$$+6 * \exp\left(-4\tau R_{2,H}^{S}\right) * \exp\left(-t_1 R_{MQ}^{S}\right) * \exp\left(-t_2 R_{2,H}^{S}\right)$$

**(Eq. S2)**

R denotes relaxation rates. The superscripts “F” and “S” correspond to fast and slow relaxation rates, respectively. The subscripts “2,H” and “2,C” characterize $^{1}$H and $^{13}$C single quantum coherence relaxation rates. The subscript “MQ” corresponds to the $^{1}$H-$^{13}$C multiple quantum coherence relaxation rate. τ is a delay in the pulse program ($1/2*^{1}J_{CH}$= 3.6 ms). $t_1$ and $t_2$ are the acquisition times in the indirect and direct dimensions, respectively. It is assumed that the product of the $^{13}$C resonance frequency $\omega_C$ and the global correlation time $\tau_C$ $\omega_C * \tau_C >> 1$, and that methyl rotation is infinitely fast. The relaxation rates in equations S1 and S2 are given with

$$R_{2,H}^{F} = \left(\frac{9}{20} * \frac{\gamma_H^4 \hbar^2 \tau_C}{r_{HH}^6} + \frac{1}{45} * \frac{\gamma_H^2 \gamma_C^2 \hbar^2 \tau_C}{r_{HC}^6}\right) * \left(\frac{\mu_0}{4\pi}\right)^2 * S^2 + R_{2,ext}$$

**(Eq. S3)**

$$R_{2,H}^{S} = \left(\frac{1}{45} * \left(\frac{\mu_0}{4\pi}\right)^2 * \frac{\gamma_H^2 \gamma_C^2 \hbar^2 \tau_C}{r_{HC}^6}\right) * S^2 + R_{2,ext}$$

**(Eq. S4)**

$$R_{2,C}^{F} = \left(\frac{1}{5} * \left(\frac{\mu_0}{4\pi}\right)^2 * \frac{\gamma_H^2 \gamma_C^2 \hbar^2 \tau_C}{r_{HC}^6}\right) * S^2 + R_{1,ext}$$

**(Eq. S5)**

$$R_{2,C}^{S} = \left(\frac{1}{45} * \left(\frac{\mu_0}{4\pi}\right)^2 * \frac{\gamma_H^2 \gamma_C^2 \hbar^2 \tau_C}{r_{HC}^6}\right) * S^2 + R_{1,ext}$$

**(Eq. S6)**

$$R_{MQ}^{F} = \left(\frac{4}{45} * \frac{\gamma_H^2 \gamma_C^2 \hbar^2 \tau_C}{r_{HC}^6} + \frac{9}{20} * \frac{\gamma_H^4 \hbar^2 \tau_C}{r_{HH}^6}\right) * \left(\frac{\mu_0}{4\pi}\right)^2 * S^2 + R_{2,ext}$$

**(Eq. S7)**

$$R_{MQ}^{S} = R_{2,ext} = \frac{8}{15} * \left(\frac{\mu_0}{4\pi}\right)^2 \frac{\gamma_H^2 \gamma_C^2 \hbar^2 \tau_C}{r_{HDext}^6} + \frac{9}{20} * \left(\frac{\mu_0}{4\pi}\right)^2 \frac{\gamma_H^4 \hbar^2 \tau_C}{r_{HHext}^6}$$

**(Eq. S8)**

$$R_{1,ext} = \frac{3}{20} * \left(\frac{\mu_0}{4\pi}\right)^2 \frac{\gamma_H^4 \hbar^2 \tau_C}{r_{HHext}^6}$$

**(Eq. S9)**

where $\mu^0$ is the vacuum permeability constant and $\hbar$ is the Planck constant. $\gamma_H$ and $\gamma_C$ are the gyromagnetic rations of $^1$H and $^{13}$C spins. $r_{HH}$ and $r_{HC}$ are the distances between $^{13}$C and $^1$H (1.135 Å) or $^1$H and $^1$H spins (1.813 Å) within a methyl group. $R_{1,ext}$ and $R_{2,ext}$ describe dipolar relaxation with external deuterons and protons. $r_{HHext}$ and $r_{HDext}$ are dependent of distances to external protons and deuterons (see below).

To account for the intensity in the final spectra, equations for $I_{HMQC}$ and $I_{HSQC}$ were integrated over the acquisitions times $t_{1,max}$ (0.0942302 s) and $t_{2,max}$ (0.1136640 s) of the indirect and direct dimensions. This is achieved by substituting the exp(-$t_1$R) and exp(-$t_2$R) terms in Eqs. S1 and S2 with (1-exp(-$t_{1,max}$R)/R and (1-exp(-$t_{2,max}$R)/R terms.

**Gauss distributions of $R^S_{MQ}$**

$R^S_{MQ}$ was derived experimentally using the pulse program *2020.R.mq.am.tb.V2* (see above) with relaxation delays Δt of 0 ms, 4 ms, 8 ms, 12 ms, 20 ms, 28 ms, 36 ms, 50 ms, 76 ms, 100 ms, and 200 ms and using resonance intensities and relaxation delays to fit $R^S_{MQ}$ and $A_0$ to an exponential decay:

$$I(\Delta t) = A_0 * \exp\left(-R^S_{MQ}\Delta t\right)$$

**(Eq. S10)**

For error estimation, standard deviations and means of intensities of methyl group resonances were estimated from duplicate intensity measurements for each methyl group resonance and relaxation delay. Standard deviations and mean of the spectral noise were calculated from 100 noise peaks. Noise peaks were obtained from the spectrum with the longest relaxation delay. For each methyl group resonance and relaxation delay, 1000 synthetical intensity values were generated by summing up randomly drawn values from synthetic Gauss distributions of the spectral noise and the respective resonance's intensity (estimated by the respective standard deviation and mean, see above). 1000 bootstrap iterations were carried out for the fitting process. In each iteration, one intensity value for each relaxation delay was drawn randomly from the corresponding synthetic intensity values. The obtained values and the corresponding relaxation delays were than used to fit $R^S_{MQ}$. This resulted in a Gauss distribution of $R^S_{MQ}$ values defined by the standard deviation and mean of the bootstrap procedure. Spectra with different relaxation delays were acquired in a random order to avoid artifacts.

**Gauss distributions of $I_{HMQC}$ and $I_{HSQC}$**

Gauss distributions of $I_{HMQC}$ and $I_{HSQC}$ values were estimated by the means and standard deviations of triplicate measurements.

**Gauss distributions of the global correlation time $\tau_C$**

Determination of $^{15}N$ $R_\alpha$ and $R_\beta$ relaxation rates was carried out by integrating the 8 ppm to 10 ppm region of TRACT spectra with 25 increasing delays $\Delta t_{TRACT}$ and fitting the integral areas and delays to an exponential decay:

$$I(\Delta t) = A_0 * \exp\left(-R_i \ \Delta t_{TRACT}\right)$$

**(Eq. S11)**

with i ϵ [α,β]. Gauss distributions of $R_\alpha$ and $R_\beta$ resulted from the obtained values and standard errors of the fit. A Gauss distribution of $\tau_C$ was generated by 1000 Monte Carlo iterations of calculating $\tau_C$ from randomly drawn values of $R_\alpha$ and $R_\beta$ Gauss distributions by solving the following equation[6]:

$$R_\beta - R_\alpha = 2\frac{\mu_0 \gamma_H \gamma_N h}{16\pi^2\sqrt{2} r_{HN}^3} * \frac{\gamma_N B_0 \Delta\delta_N}{3\sqrt{2}} * \left(1.6\tau_C + \frac{1.2\tau_c}{1+\tau_C^2\omega_N^2}\right) * (3 * \cos^2(\theta) - 1)$$

**(Eq. S12)**

where $\gamma_N$ is the gyromagnetic ration of $^{15}N$ spins, h is the Planck constant, $r_{HN}$ is the distance between $^1H$ and $^{15}N$ spins within the N-H bond (1.02 Å), $B_0$ is the external magnetic field strength, $\Delta\delta_N$ is the difference of the two principal components of the axially symmetric $^{15}N$ chemical shift tensor (160 ppm), and $\omega_N$ is the $^{15}N$ resonance frequency. The angle Θ is assumed to be 17°.

**Gauss distribution of $r_{HHext}$**

Gauss distributions of $r_{HHext}$ were obtained by calculating means and standard deviations resulting from 1000 Monte Carlo iterations of solving equation S8 with values for $\tau_C$ and $R^S_{MQ}$ randomly drawn from their respective Gauss distributions.

**Calculation of $r_{HDext}$ from a crystal structure model**

The crystal structural model of MNV P-domain in presence of GCDCA, metal ions, and CD300lf (pdb 6e47, Nelson et al.[3]) was used to obtain the mean distance $r_{HDext}$ of the three protons of a certain methyl groups to the protein's deuterons using:

$$r_{HDext} = \frac{1}{3} \sum_{j=1-3} \sum_{i} \frac{1}{r^6_{H_j D_i}}$$

**(Eq. S13)**

where proton "j" corresponds to the methyl group of question and deuteron "i" can be any deuteron of the protein.

**Calculating $S^2$ methyl group order parameters**

For the calculation of methyl group order parameters, 1000 Monte Carlo iterations were performed employing the corresponding $r_{HDext}$ and drawing random values from Gauss distributions of $\tau_C$, $R^S_{MQ}$, $r_{HHext}$, $I_{HMQC}$, and $I_{HSQC}$. The values were used to fit $S^2$ globally according to equations S1 to S9. The resulting mean and standard deviation are given as the final $S^2$ value and its corresponding error, respectively. *Python*'s *lmfit* and *numpy* libraries were used for the calculations.
